# Astrocytic GABA transport controls fidelity of temporal processing

**DOI:** 10.1101/2022.02.08.479618

**Authors:** Victoria A. Wagner, Ritika Thapa, Anna O. Norman, Alexandra Varallo, Jordan Wimberly, Maham Rais, Dmytro Gerasymchuk, Timo P. Piepponen, Khaleel A. Razak, Maija L. Castren, Iryna M. Ethell

**Affiliations:** Neuroscience Graduate Program, University of California Riverside, Riverside, CA, USA; Division of Biomedical Sciences and Biomedical Sciences Graduate Program, School of Medicine, University of California Riverside, Riverside, CA 92521; Department of Physiology, Medical Faculty, and Division of Pharmacology and Pharmacotherapy, University of Helsinki, Helsinki, Finland; Department of Psychology, University of California Riverside, Riverside, CA, USA

**Keywords:** Autism, Fragile X Syndrome, Neurodevelopmental Disorders, Astrocytes, GABA Transport, Parvalbumin Inhibitory Interneurons, Electrocortical Activity, Cortical Circuits

## Abstract

Astrocytes control neural communications by influencing GABAergic transmission through uptake and synthesis of GABA. Impaired GABAergic signaling is thought to underlie cortical hyperexcitability in autism. Here we show that dysregulation of astrocyte GABA transport in Fragile X syndrome (FXS), a leading genetic cause of autism, contributes to circuit hyperexcitability. Human FXS astrocytes derived from patient-specific induced pluripotent stem cells and mouse *Fmr1* knockout (KO) astrocytes display a significant increase in levels of GABA and GABA-synthesizing enzyme GAD65/67. Our astrocyte-specific *Fmr1* KO (cKO) mouse model reveals reduced inhibitory connectivity and impaired cortical responses to sound. Reverse GABA transport in cortical astrocytes contributes to impaired fidelity of temporal processing and hyperactive behaviors in cKO mice. Blocking astrocyte GABA transport is sufficient to restore PV expression, cortical activity, EEG responses, and locomotor behavior. Our findings suggest astrocyte GABA transport plays a key role in regulating cortical inhibition, and contributes to autism-associated phenotypes.

**Highlights:**

- FXS astrocytes show elevated levels of GABA and its synthesizing enzyme GAD65/67
- *Fmr1* KO cortical astrocytes suppress PV expression leading to enhanced overall cell activity through reverse GABA transport
- Astrocyte-specific postnatal deletion of *Fmr1* results in reduced inhibitory cortical connectivity, impaired fidelity of temporal processing and behavioral hyperactivity
- Acute blockade of astrocytic GABA transport is sufficient to restore cortical responses and correct hyperactive mouse behaviors

**Graphic Abstract:** 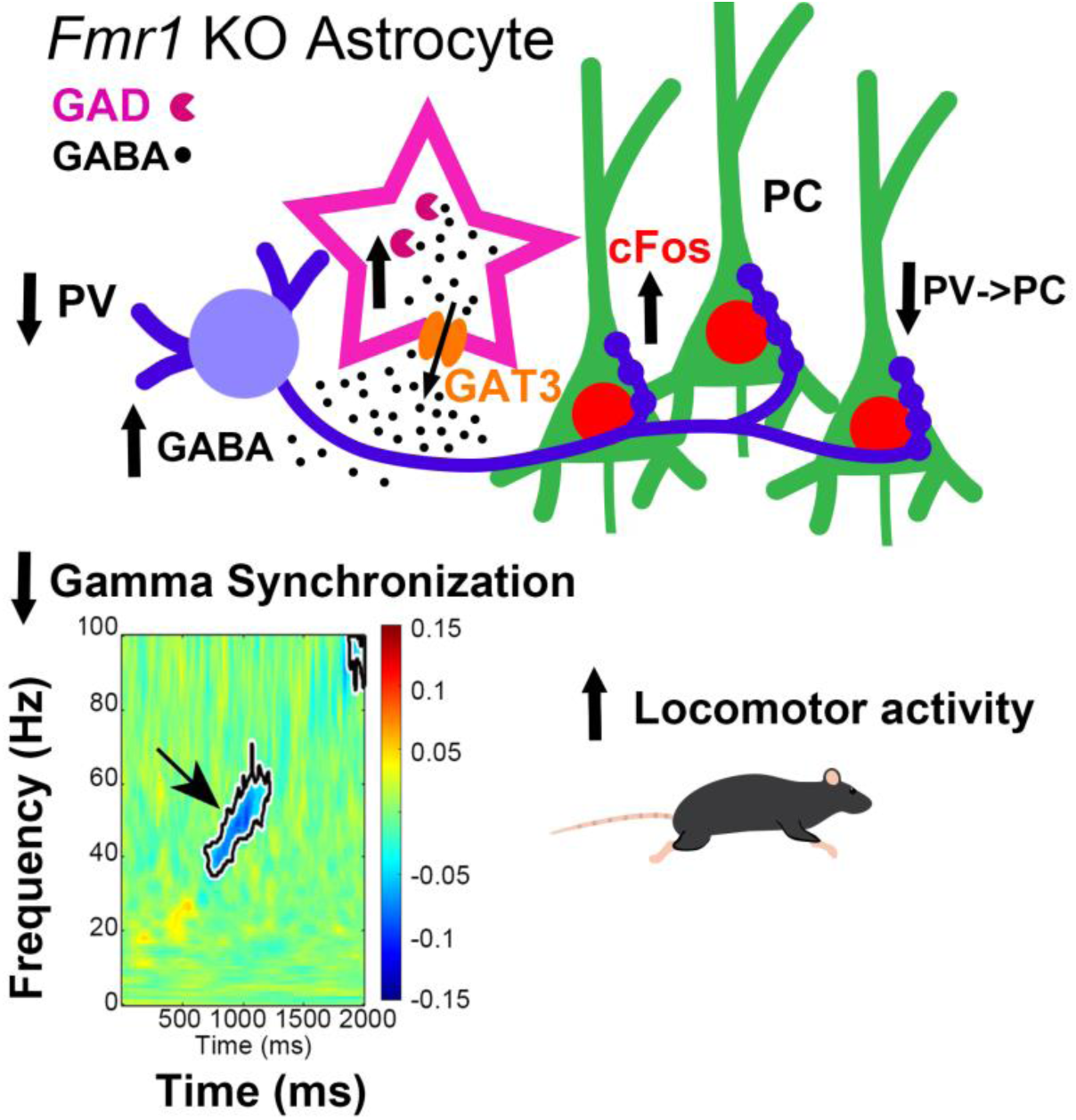

## Introduction

Regulation of cortical excitability is important for sensory processing and motor behaviors and is supported by dynamic interactions between excitatory neurons, interneurons, and astrocytes^1^. Increased ratio of excitation/inhibition (E/I) in the cortex is thought to contribute to features of autism spectrum disorders (ASD) including hypersensitivity, sensory processing deficits, language delays, repetitive behaviors, anxiety, and sleep disturbances^2,3^. Impaired inhibition is proposed to mediate cortical hyperexcitability in ASD, based on observations including delayed developmental shift from depolarizing to hyperpolarizing gamma-aminobutyric acid (GABA), parvalbumin (PV) interneuron hypofunction, and reduced expression of GABA receptors^4–8^. However, the complex etiology of ASD cases can introduce challenges to unraveling the emergence of inhibitory impairments.

Fragile X Syndrome (FXS) is the most common genetic form of ASD^9^. FXS is most often caused by a CGG repeat expansion in the 5’-untranslated region of the *Fragile X messenger ribonucleoprotein 1* (*Fmr1*) *gene* followed by promoter methylation, leading to the loss of Fragile X Messenger RibonucleoProtein (FMRP), translational dysregulation, and abnormal protein synthesis^10,11^. Prominent symptoms of FXS include intellectual disability, developmental delays, repetitive behavior, and hyperarousal seen as sensory hypersensitivity^12,13^, particularly in the auditory domain^14–16^. Sensory hypersensitivity is observed in both humans with FXS and mouse models of FXS and can stem from hyperexcitable cortical circuits^13–16^.

The *Fmr1* knockout (KO) mouse is an established FXS model that is well suited to study cellular mechanisms of cortical hyperexcitability. FMRP loss increases network-level excitability in the rodent cortex through impaired inhibition and altered excitatory/inhibitory (E/I) balance^17^ that most likely affects neural synchrony^18,19^. The *Fmr1* KO mice show decreased density and function of parvalbumin (PV)-expressing inhibitory interneurons^20–23^, reduced GABA receptor levels, altered GABA metabolism, and overall decreased GABAergic input in several areas of the brain^24–27^. Studies to date suggest defective communications between excitatory pyramidal neurons and inhibitory interneurons^20,28^ in the absence of FMRP that result in abnormal E/I balance^29^. Although FMRP is involved in the regulation of synaptic function and neural communication^30–32^, little is known how astrocytes may contribute to abnormal inhibition.

Astrocytes control neural communication, not only by regulating glutamate metabolism, but also through uptake and synthesis of GABA, thereby influencing GABAergic transmission^33,34^. However, most studies of astrocytes are focused on excitatory synapses and pyramidal cells (PC), leaving inhibitory synapses, and GABAergic interneurons less explored^35^. *Fmr1* KO astrocytes are known to trigger FXS-specific changes in wild-type (WT) excitatory neurons^36–38^. The enhanced excitability was also linked to reduced expression of glutamate transporter in FMRP-deficient astrocytes and reduced glutamate uptake by the astrocytes^36,39^. Whether astrocytes contribute to abnormal inhibitory cell development and circuit hyperexcitability by altering GABAergic signaling in FXS is unclear.

Therefore, the main goal of the present study was to determine whether cell-specific deletion of *Fmr1* from astrocytes during the critical developmental period of inhibitory circuit refinement^40–43^ would alter GABAergic signaling and PV cell development, leading to cortical hyperexcitability and behavioral alterations, and further, whether these features could be rescued by selectively blocking astrocyte GAT3-mediated GABA transport. We used a multidisciplinary approach including transcriptional, cellular, molecular, electrophysiological, and behavioral methods to delineate the astrocyte-mediated mechanisms of abnormal inhibition in FXS. We utilized both human and mouse astrocytes, and the analysis of translationally relevant electroencephalogram (EEG) phenotypes that are remarkably similar between the mouse model of FXS and the human condition across different brain areas implicated in FXS-associated behaviors^44^. Treatment to block GAT3-mediated GABA transport in astrocytes improved cortical responses, reversing EEG and behavioral phenotypes. Our findings provide novel insight into the role of astrocyte GABA transport in the development of cortical circuits and fidelity of temporal processing in the cortex, and suggest a non-neuronal mechanism of inhibitory circuit dysregulation in FXS with potential for translating the results of the mouse study into successful clinical trials.

## Results

### GABA levels are elevated in human iPSC-derived FXS astrocytes

To investigate the role of astrocytes in abnormal inhibition in FXS, we first investigated GABA levels in human astrocytes. Human astrocytes were differentiated from FXS-(SO02b, SO04a, SO10c3, HEL70.3, HEL70.6, HEL100.2, and HEL69.6) and control-derived (V36C, V44D, sc176c3a, HEL23.3, HEL24.3, and P04) iPSCs using a forebrain astrocyte model as previously described^45^ (**Figure 1A**) and collected after day (D)95-98 for analysis. Differentiated FXS and control cells displayed mature astrocyte morphology with over 90% of cells immunopositive for astrocyte markers GFAP and S100B (**Figure 1B, Table S1**).

**Figure 1.**
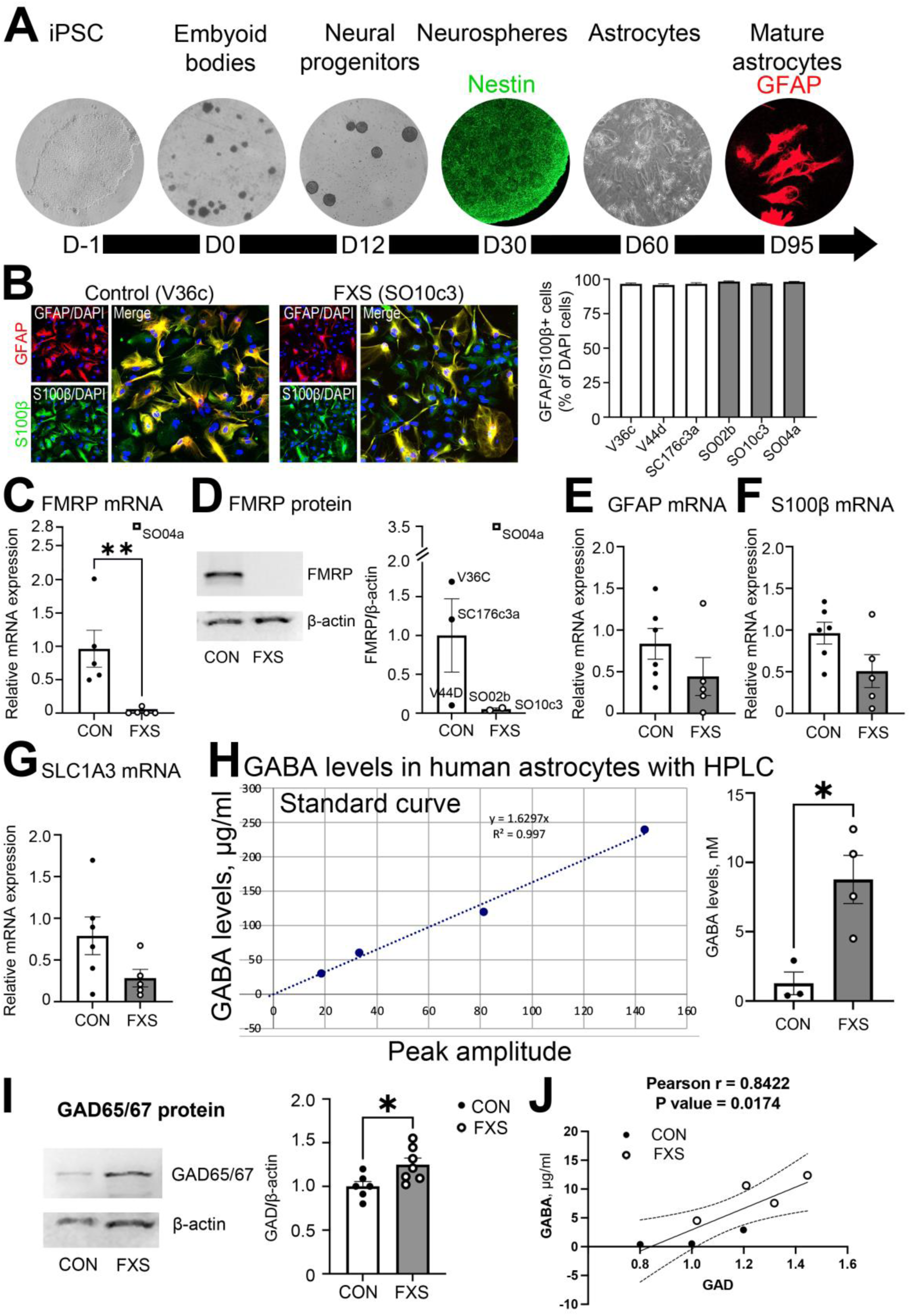
GABA levels are elevated in human iPSC-derived FXS astrocytes. A, Graphics showing timeline of astrocyte differentiation from human iPSC lines, including representative bright field images of cells, and confocal images of cells immunolabeled for neural progenitors with Nestin and astrocytes with GFAP. B, Representative images of day (D)95 differentiated FXS and control astrocytes immunolabeled with astrocytic markers S100b and GFAP, and overlaid with DAPI. Graph shows proportion of cells immunoreactive for astrocytic markers for each cell line (n=3 cell lines/group). C, Graph shows mRNA expression of *Fmr1* in FXS and control astrocytes. *Fmr1* mRNA expression is significantly reduced in FXS astrocytes, except one FXS cell line (SO04a) with abnormally high *Fmr1* expression, which was determined to be an outlier. Graph shows mean ± SEM (n=5-6 cell lines/group, **p<0.01; t-test). D, Immunoblots for FMRP and beta-actin in lysates from FXS and control (CON) astrocytes. One cell line (SO04a) had abnormally high FMRP expression, which was determined to be an outlier, and was excluded from subsequent analysis. Graph shows mean ± SEM (n= 3 cell lines/group, p=0.05; t-test). E-G, Graphs show mRNA expression of GFAP (E), S100B (F), and SLC1A3 (G) in FXS and control astrocytes. Graph shows mean ± SEM (n= 5-6 cell lines/group, p=0.05; t-test). No significant differences were observed in expression of astrocyte-specific genes. H, GABA detection in FXS and control astrocytes with HPLC analysis. Graphs show standard curve and average GABA levels in CON and FXS samples ± SEM (n= 3-4 cell lines/group, *p<0.05; t-test). FXS iPSC-derived astrocytes have increased GABA levels compared to controls. I, Immunoblots for GAD65/67 and beta-actin in lysates from FXS and CON astrocytes. Graph shows mean ± SEM (n= 6-7 cell lines/group, *p<0.05; t-test). GAD65/67 levels are significantly increased in FXS astrocytes compared to controls. J, Correlation between GABA and GAD65/67 levels in FXS and CON astrocytes (n= 3-4 cell lines/group, *p<0.05; Pearson correlation). GABA levels are strongly, positively correlated with GAD65/67 levels.

We observed only traces of FMRP expression in astrocytes differentiated from most FXS iPSC lines compared to control astrocytes (**Figure 1C, 1D, Table S1**). However, astrocytes derived from FXS iPSC line SO04a showed significant FMRP levels that were higher than in controls and were excluded from subsequent analysis (**Figure 1C, 1D, Table S1**). Real-time qPCR analysis showed that the expression levels of astrocyte-specific genes encoding GFAP, S100B, and GLAST did not differ in FXS and control astrocytes (**Figure 1E-1G, Table S1**).

We first assessed GABA levels in human astrocytes using HPLC and found a 7-fold increase in GABA in FXS astrocytes compared with control astrocytes (**Figure 1H, Table S1**). In addition, the levels of glutamic acid decarboxylase (GAD65/67), GABA synthesizing enzyme, were significantly upregulated in astrocytes derived from FXS iPSCs, assessed by western blot analysis (**Figure 1I, Table S1**). Furthermore, GABA levels were significantly correlated with GAD65/67 levels by cell line (Pearson *r* = 0.8422; **Figure 1J, Table S1**), suggesting that increased GABA levels in FXS astrocytes may arise from increased GABA synthesis from glutamate via GAD65/67.

In summary, iPSC-derived FXS astrocytes demonstrate elevated GABA levels potentially due to enhanced GABA synthesis from glutamate.

### Levels of GABA and its synthesizing enzyme are significantly upregulated in *Fmr1* KO astrocytes

Our data indicate that FXS human astrocytes, where *Fmr1 gene* is silenced as a result of CGG repeat expansion and gene methylation, have elevated GABA levels. We next investigated if *Fmr1 gene* deletion in astrocytes would also result in elevated GABA levels. Indeed, we observed that cultured *Fmr1* KO mouse astrocytes had higher levels of GABA detected with immunolabeling (**Figure 2A, Table S2**). Similar to FXS human iPSC-derived astrocytes, *Fmr1* KO mouse astrocytes also showed increased protein levels of GABA-synthesizing enzyme GAD65/67 (**Figure 2B**). We found no changes in the protein levels of another GABA synthesizing enzyme Aldha1, that is known to convert putrescene to GABA (**Figure 2B, Table S2**). In brief, increased GABA levels and its synthesizing enzyme GAD65/67 were observed in both FMRP deficient human and mouse astrocytes *in vitro*.

**Figure 2.**
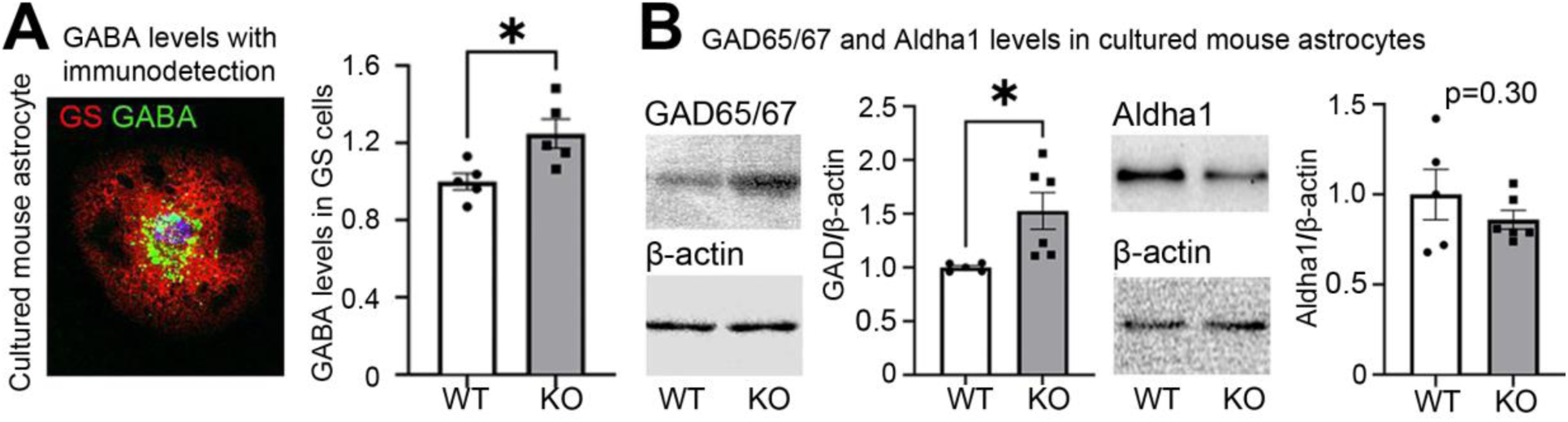
Levels of GABA and its synthesizing enzyme are significantly upregulated in Fmr1 KO astrocytes. A-B, GABA immunodetection (A) and immunoblot for GAD65/67, Aldha1 and beta-actin (B) in cultured astrocytes from WT and *Fmr1* KO mice. Graphs show mean ± SEM (n= 5-6 cultures/group, *p<0.05; t-test). GABA levels and GABA synthesizing enzyme GAD65/67 are significantly increased in KO astrocytes compared to controls.

### Astrocyte-specific FMRP deletion during postnatal developmental period results in excess GABA release from astrocytes

As astrocytes play an important role in regulating extracellular GABA levels in the brain, changes in astrocytic GABA levels may influence neuronal activity. Therefore, we next examined how increased GABA levels in *Fmr1* KO astrocytes may affect the development of neuronal circuits in the mouse cortex.

To achieve specific *Fmr1* deletion in astrocytes, ERT2-Cre^GFAP^*Fmr1*^flox/y^ conditional KO mice (cKO) were generated and ERT2-Cre^GFAP^wild-type (Ctrl WT) mice were used as controls. For analysis of FMRP levels in astrocytes, tdTomato was expressed in astrocytes using Rosa-CAG-LSL-tdTomato reporter mice to generate tdTomato-ERT2-Cre^GFAP^*Fmr1*^flox/y^ cKO mice and tdTomato-ERT2-Cre^GFAP^ Ctrl WT mice. Ctrl WT and cKO mice received tamoxifen at postnatal day (P)14 intraperitoneally (IP, 0.5 mg in 5 mg/ml of 1:9 ethanol/sunflower seed oil solution) once a day for 5 days, and analysis was performed at P28 (**Figure 3A, Table S3**).

**Figure 3.**
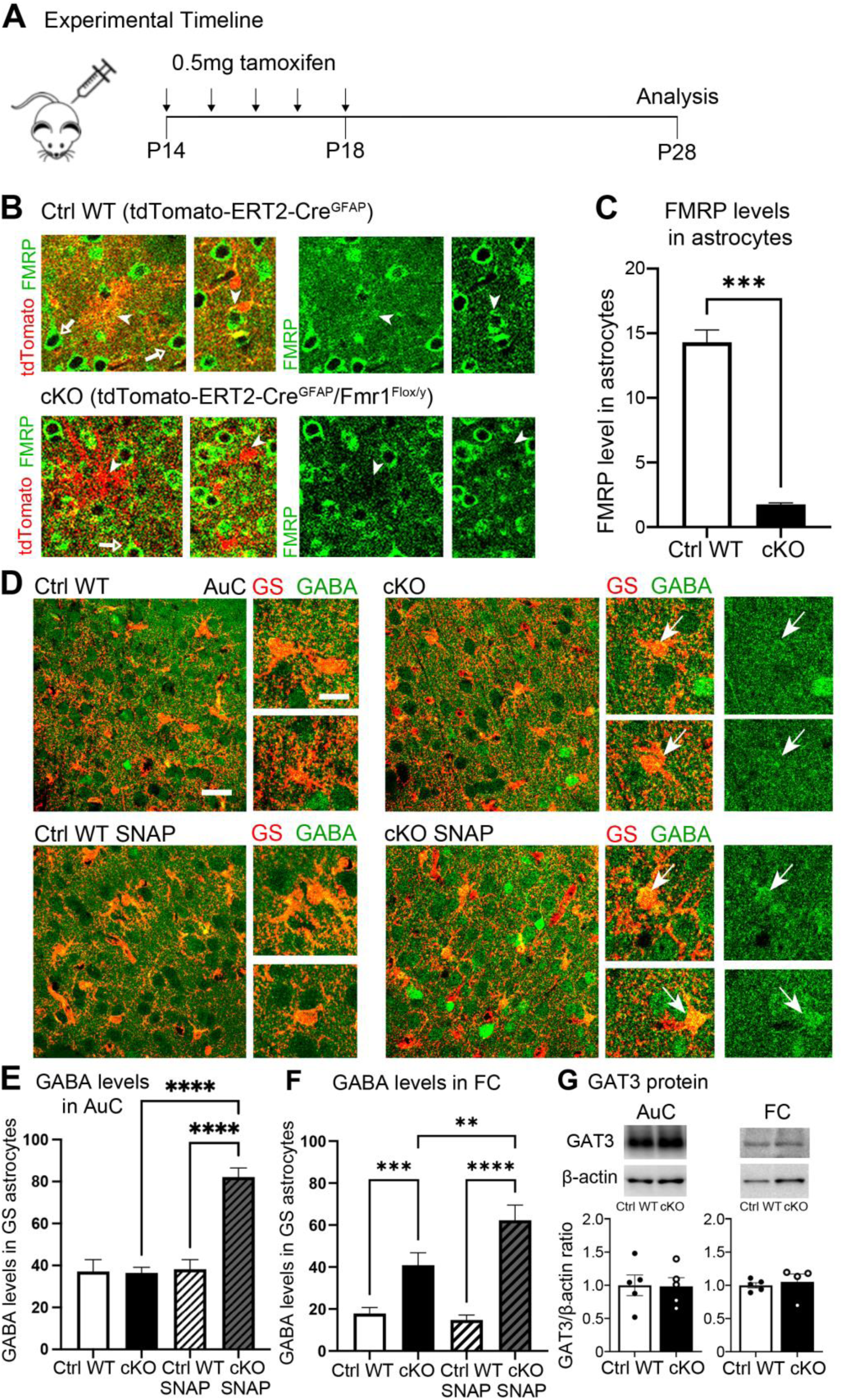
Astrocyte-specific FMRP deletion during postnatal developmental period results in excess GABA release from astrocytes. A, Graphics showing experimental timeline of *Fmr1* deletion in astrocytes. To achieve FMRP deletion in astrocytes tamoxifen (0.5 mg) was intraperitoneally (IP) injected at P14 for 5 days; experiments were performed at P28. The transgenic mouse groups used in this study were: (1) ERT2-Cre*^GFAP^* control (Ctrl WT) and ERT2-Cre*^GFAP^Fmr1*^flox/y^ conditional KO (cKO); (2) tdTomato-ERT2-Cre*^GFAP^* (Ctrl WT) and tdTomato-ERT2-Cre*^GFAP^Fmr1*^flox/y^ (cKO). B, Confocal images showing tdTomato astrocytes (red) and FMRP immunoreactivity (green) in neurons (arrow) and astrocytes (arrowhead) in auditory cortex (AuC) of Ctrl WT (upper panels) and cKO (lower panels) mice at P28. C, Quantitative analysis of FMRP levels in astrocytes in the AuC at P28. Graph shows mean ± SEM (n= 3 mice/group, ***p<0.001; t-test). FMRP levels in astrocytes were significantly reduced in cKO mice compared to controls. D, Confocal images showing Glutamine Synthetase (GS, red) and GABA (green) immunoreactivity in superficial layers of AuC of vehicle-treated and SNAP-treated Ctrl WT and cKO mice. Arrows indicate GS-labeled cell bodies of astrocytes. E-F, Quantitative analysis of the intensity of GABA immunoreactivity in GS-labeled astrocytes in AuC (E) and FC (F). Graphs show mean ± SEM (n= 3-5 mice/group, **p<0.01,***p<0.001, ****p<0.0001; two-way ANOVA, Fisher’s LSD post hoc test). GABA levels in GS-labeled astrocytes were significantly increased in cKO mice compared to controls in FC and in cKO SNAP mice compared to all other groups in AuC and FC. G, Western blots showing GAT3 and beta-actin protein levels in lysates from AuC (left) and FC (right) from Ctrl WT and cKO mice. Graphs show mean ± SEM (n=4-5/group; p=0.05; t-test). No differences were observed in GAT3 protein levels.

To confirm specific ablation of *Fmr1* in astrocytes, FMRP immunoreactivity was analyzed in the cortex of tdTomato-ERT2-Cre^GFAP^Ctrl WT and tdTomato-ERT2-Cre^GFAP^*Fmr1*^flox/y^ cKO mice at P28 (**Figure 3B, Table S3**). FMRP immunoreactivity was significantly decreased in cortical astrocytes of cKO mice compared to Ctrl WT (**Figure 3C, Table S3**), confirming successful deletion of FMRP specifically from cortical astrocytes during the postnatal P14-P28 window.

We next assessed whether cortical astrocytes in our cKO mouse model retained high GABA levels as observed in cultured *Fmr1* KO mouse and FXS human astrocytes. While astrocytic GABA levels were higher in cKO mice compared to Ctrl WT in L2-4 frontal cortex (FC), there were no genotype differences in astrocyte GABA levels in L2-4 auditory cortex (AuC) (**Figure 3D-3F, Table S3**). It is possible that elevated astrocyte GABA levels compared to extracellular space may result in reverse GABA transport in KO astrocytes. To address this possibility, we investigated the effects of acute blockage of astrocyte GABA transport through GAT3 using synthetic drug SNAP-5114 (SNAP). While we found no significant changes in GABA levels in WT astrocytes after SNAP treatment, KO astrocytes revealed a two-fold increase in astrocytic GABA in L2-4 in both AuC and FC of cKO mice treated with SNAP (**Figure 3D-3F, Table S3**). This observation confirms that cortical astrocytes in cKO mice are synthesizing more GABA than controls, similar to our FXS human and mouse culture models, leading to reverse GABA transport in astrocytes.

We also investigated if expression of astrocyte-specific GABA transporter GAT3 was altered in our cKO model, and could potentially account for any influence on astrocytic GABA trafficking. Our analysis determined no changes in GAT3 mRNA and protein levels in the FC and AuC in cKO mice compared to Ctrl WT (**Figures 3G, S1, Tables S3, S8**). Here we have shown that *Fmr1* KO astrocytes *in vivo* recapitulate elevated GABA synthesis observed in cultured astrocytes, which can be a source of elevated extracellular GABA via reverse GABA transport in astrocytes.

### Inhibitory and parvalbumin positive connectivity is reduced in astrocyte-specific *Fmr1* cKO mice

Extracellular GABA may serve as a cue for microglia to initiate removal of inhibitory synapses^46^. It is possible that reverse GABA transport from astrocytes can trigger pruning of GABAergic synapses, which may be responsible for deficits in inhibitory connectivity observed in FXS^47–49^. Therefore we next examined the role of astrocytes in the development of inhibitory connectivity in the cortex of astrocyte-specific *Fmr1* KO mice. We detected inhibitory synapses by immunolabeling pre-synaptic VGAT and postsynaptic gephyrin puncta and analyzing their colocalization (**Figure 4A, Table S4**). We observed a significant decrease in the density of colocalized VGAT/Gephyrin as well as VGAT puncta in L2-4 AuC and FC in cKO mice compared to Ctrl WT, suggesting reduced inhibitory innervation of pyramidal cells (PCs) (**Figure 4C-4D, Table S4**). As previous studies have shown reduced expression of GABA_A_ receptors in FXS^25,26,50–53^, we also analyzed GABA_A_ receptor expression in AuC and FC of cKO mice. However, we found no significant differences in GABA_A_R protein levels in our model (**Figure S2, Table S9**), suggesting a pre-synaptic mechanism.

**Figure 4.**
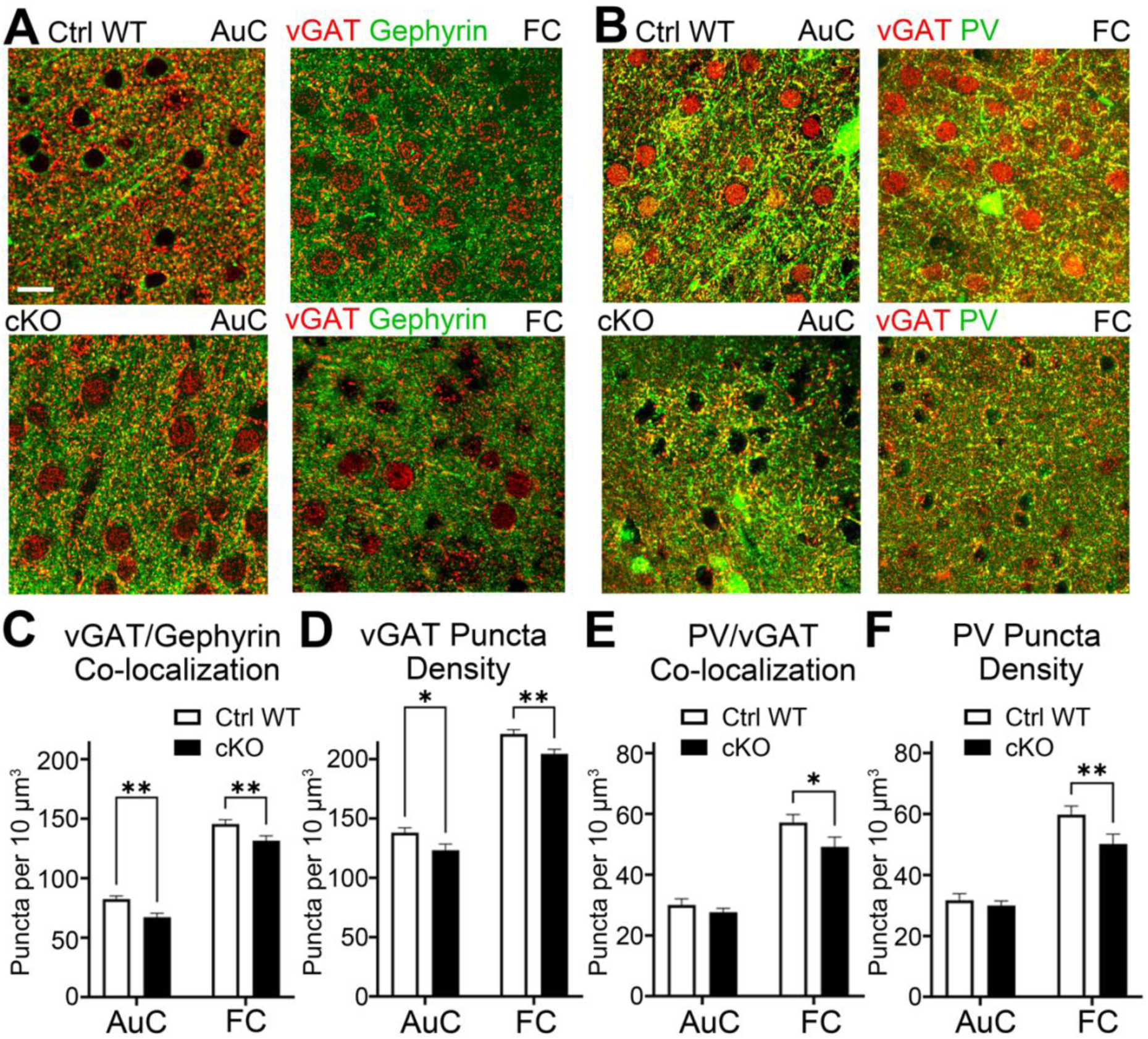
Inhibitory and parvalbumin positive connectivity is reduced in astrocyte-specific Fmr1 cKO mice. A-B, Confocal images showing immunolabeled inhibitory synaptic sites in L2/3 AuC (left) and FC (right) of P28 Ctrl WT (upper row) and cKO (lower row) mice. Presynaptic boutons were visualized with vGAT (red) and postsynaptic sites with Gephyrin (green) (A). Parvalbumin (green) positive inhibitory pre-synaptic sites were visualized with vGAT (red) (B). Scale bar 15 µm. C-F, Bar graphs show colocalized vGAT and Gephyrin puncta depicting structural inhibitory synapses (C) and density of vGAT puncta alone (D), colocalized vGAT and PV puncta to visualize PV-positive inhibitory pre-synaptic boutons (E) and density of PV immunoreactive puncta (F). Graphs show mean ± SEM (n= 3-5 mice/group, n = 5-8 images/mouse, *p<0.05, **p<0.01; two-way ANOVA, Fisher’s LSD post hoc test). Inhibitory synaptic sites and total VGAT puncta were reduced in AuC and FC of cKO mice compared to controls. Parvalbumin positive inhibitory synaptic sites were significantly reduced in FC, but not AuC of cKO mice compared to controls.

We next asked whether the deficit in inhibitory innervation was due to impaired PV cell innervation of PCs. PV-expressing cells are fast-spiking inhibitory interneurons that provide temporal control to excitatory activity and their loss or hypofunction was suggested to contribute to cortical hyperexcitability in individuals with autism and models of autism, including FXS^4,21–23,54–58^. Therefore, we next assessed PV-positive inhibitory pre-synaptic sites by analysis of VGAT and PV colocalization (**Figure 4B, Table S4**). We observed a significant decrease in the density of colocalized PV/VGAT puncta as well as overall PV puncta in L2-4 FC, with a similar trend in AuC, of cKO mice compared to Ctrl WT, suggesting reduced PV innervation of PC (**Figure 4E-4F, Table S4**). Taken together, our results demonstrate that the loss of astrocytic FMRP reduces inhibitory synapse formation onto pyramidal neurons, which may be accounted for, at least in part, by the reduction in PV synapses.

### Acute blockade of reverse GABA transport in astrocytes restores PV levels and overall cortical activity in astrocyte-specific cKO mice

Reverse GABA transport in astrocytes can elevate extracellular GABA levels, affecting neuronal activity via tonic inhibition^59–61^. Tonic inhibition differentially affects neuronal subtypes with cortical interneurons, particularly PV cells, being disproportionately suppressed by extracellular GABA; and GAT3 antagonism specifically enhances inhibitory neurotransmission in the cortex^59,62,63^. As PV protein is expressed in an activity-dependent manner, making it an effective indicator of PV neuronal activity^64,65^, we next examined whether the excess GABA release from astrocytes in our cKO model can attenuate PV activity by analyzing PV levels in cortical interneurons. We found PV levels to be significantly reduced in the AuC in cKO mice compared to Ctrl WT, without affecting density of PV cells (**Figure 5A, 5C-5D, Table S5**), more evident in L1-4 with a similar trend in L5/6 (**Figure S3A, Table S10**). There was also a significant increase in the proportion of low PV-expressing cells in cKO compared to Ctrl WT, suggesting lower PV cell activity (**Figure S3B, Table S10**). To examine whether GABA reverse transport contributes to reduced PV levels, we investigated effects of blocking GABA transport in astrocytes with acute 4h SNAP treatment. Indeed, we observed that PV levels were restored to WT levels in SNAP-treated cKO and were significantly elevated compared to vehicle-treated cKO groups in all layers of AuC (**Figures 5C, S3, Tables S5, S10**). Our findings suggest that excess GABA release from astrocytes suppresses PV cell activity; and PV cell activity is restored by blocking astrocytic GABA release in cKO mice.

**Figure 5.**
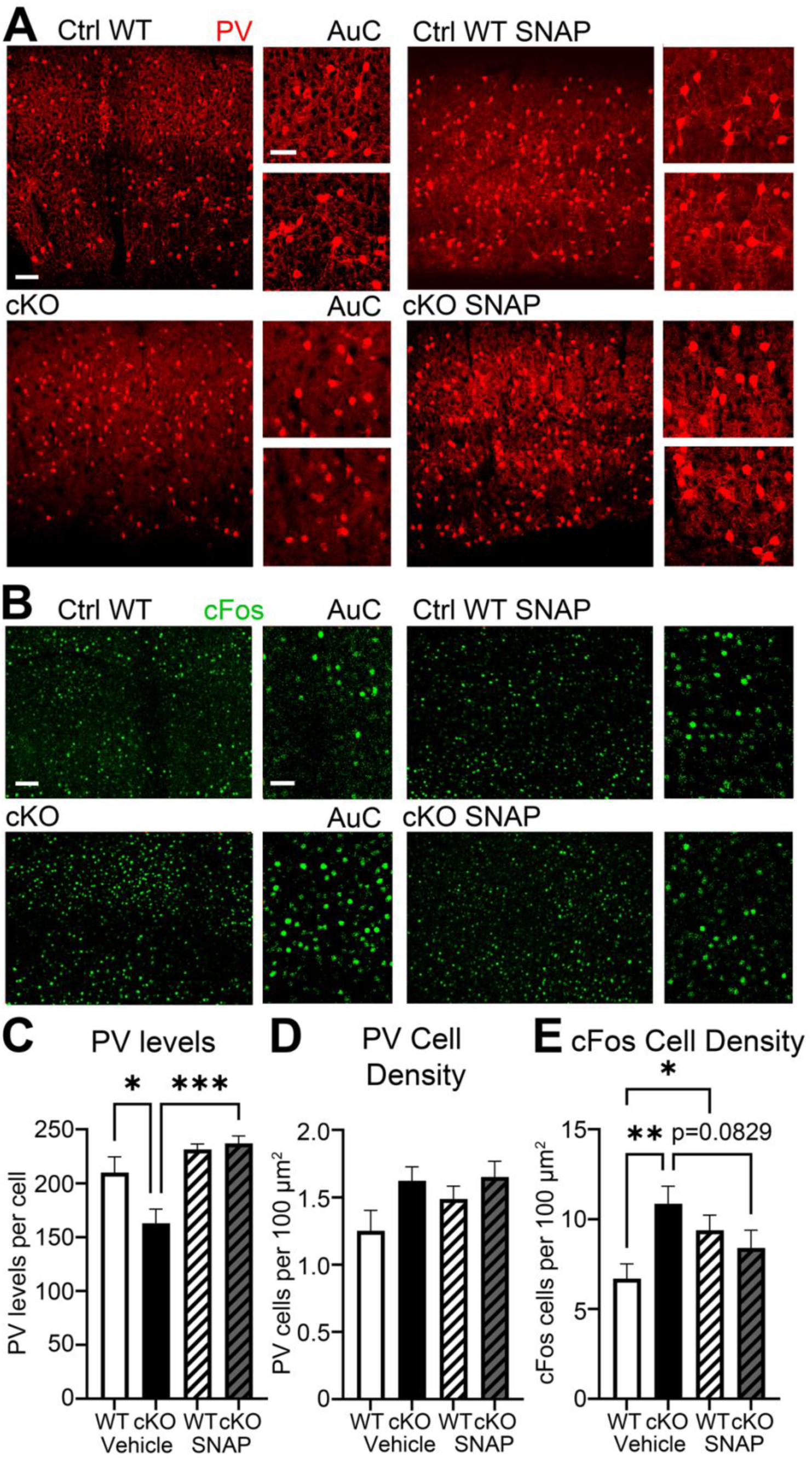
Acute blockade of reverse GABA transport in astrocytes restores PV levels and overall cortical activity in astrocyte-specific cKO mice. A, Confocal images showing immunolabeled parvalbumin cells in L1-6 AuC of P28 Ctrl WT (upper row) and cKO (lower row) mice with either vehicle (left) or SNAP treatment (right), including high magnification images from L2/3 and L5/6. Scale bar 80 µm in full size image and 40 µm in high magnification image. B, Confocal images showing immunolabeled cFos positive cells in L1-4 AuC of P28 Ctrl WT (upper row) and cKO (lower row) mice with either vehicle (left) or SNAP treatment (right), including high magnification images of L2/3. Scale bar 80 µm in full size image and 40 µm in high magnification image. C, PV levels are assessed by measuring mean intensity or PV immunoreactivity per cell. Graph shows mean ± SEM (n= 3-5 mice/group, n = 4 images/mouse, *p<0.05, ***p<0.001; two-way ANOVA, Tukey’s post hoc test). PV levels are significantly reduced in cKO mice and restored to WT levels after SNAP treatment. D, PV cell density per 100 µm^2^. Graph shows mean ± SEM (n= 3-5 mice/group, n = 4 images/mouse, p=0.05; two-way ANOVA, Tukey’s post hoc test). No significant differences were found in PV cell density. E, cFos positive cell density per 100 µm^2^. Graph shows mean ± SEM (n= 3-5 mice/group, n = 4 images/mouse, *p<0.05, **p<0.01; two-way ANOVA, Fisher’s LSD post hoc test). cFos positive cell density was significantly increased in cKO mice compared to Ctrl WT and SNAP treatment restored it to WT levels.

We next investigated whether reduced PV levels and their rescue with SNAP would influence overall cell activity in the cortex contributing to cortical hyperexcitability. We detected active cells by immunolabeling for the immediate early gene, cFos (**Figure 5B, Table S5**) and found a significantly higher density of cFos-positive cells in all layers of AuC in cKO mice that was restored to WT levels with SNAP treatment (**Figure 5E, Table S5**).

These observations indicate that reverse GABA transport in astrocytes contributes to cortical hyperexcitability in AuC of cKO mice, most likely, via inhibition of PV cell activity.

### Acute blockade of GAT3-mediated astrocytic GABA transport enhances baseline EEG power of high frequency gamma oscillations and restores low/high frequency power coupling

Enhanced cortical excitability has been previously observed in individuals with FXS and models of FXS, as well as in ASD patients and models using EEG recordings^15,66–69^. FXS-associated EEG phenotypes include enhanced baseline gamma power, abnormal timing of responses to sound (synchronization) in gamma frequency, and impaired habituation to repeated sound trains. These EEG phenotypes were also linked to loss or hypoactivity of PV cells. Therefore, we next tested cortical activity in cKO mice compared to WT and how it was affected by SNAP treatment using EEG recordings. Power spectral density of baseline (no sound stimulation) EEG responses was analyzed for AuC and FC of freely-moving Ctrl WT and cKO, before and after SNAP treatment. Although we have not observed genotype differences between WT and cKO mice that are reported in global *Fmr1* KO mice (**Figure 6G-6H, Table S6**), enhanced high gamma oscillations are apparent in cKO following SNAP treatment in AuC (**Figure 6A, 6C, Table S6**) and FC (**Figure 6B, 6D, Table S6**). SNAP treatment also enhanced high gamma power in global *Fmr1* KO mice compared to vehicle treated control with similar effects in WT mice (**Figure 6G-6H, Table S6**). In addition, we observed reduced delta power following SNAP treatment in FC of WT and cKO mice (**Figure 6D, 6G-6H, Table S6**).

**Figure 6.**
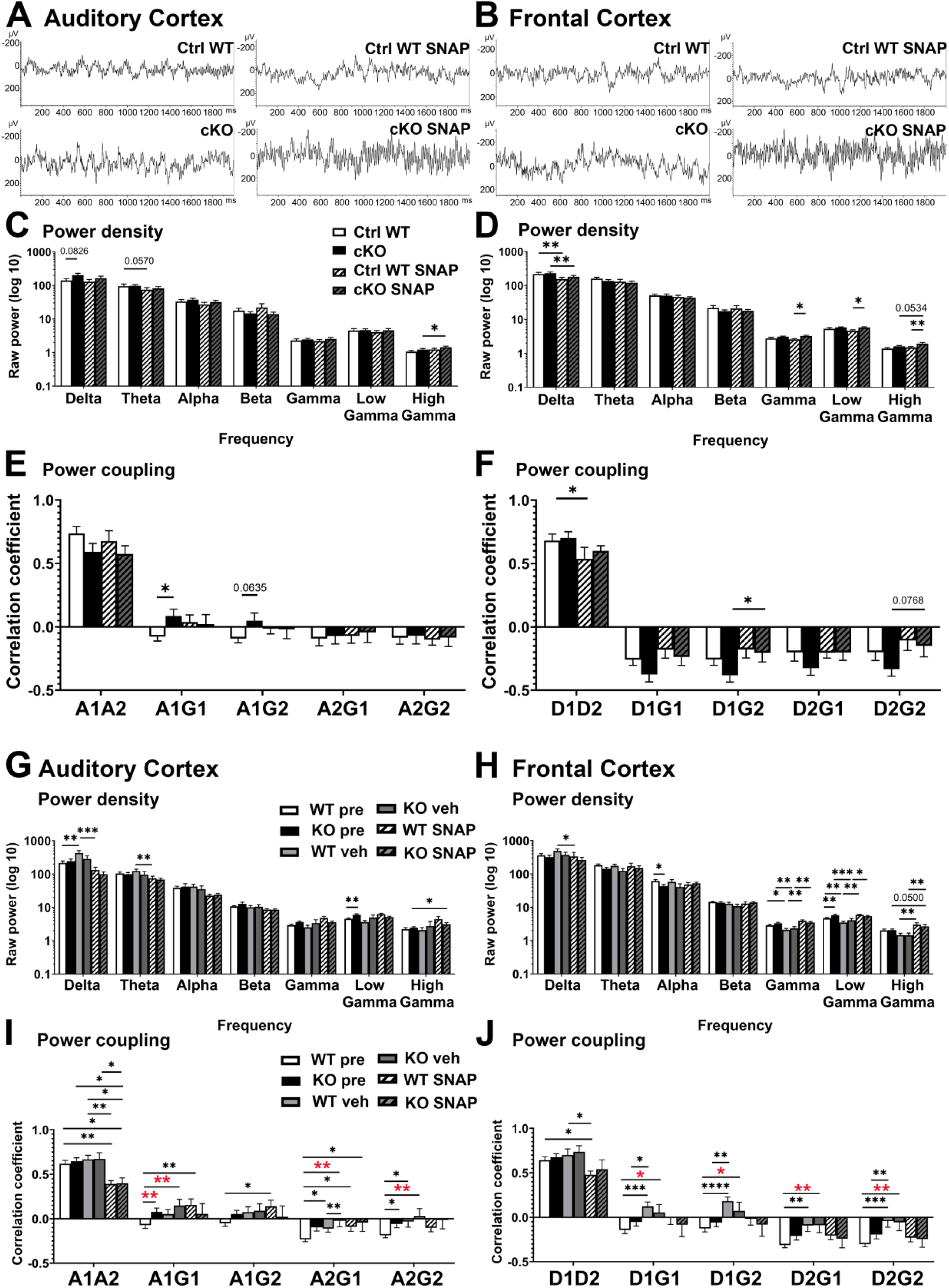
Acute blockade of GAT3-mediated astrocytic GABA transport enhances baseline EEG power of high frequency gamma oscillations and restores low/high frequency power coupling. A-B, Examples of representative EEG traces (in the absence of auditory stimulation) from electrodes implanted in AuC (A) and FC (B) in Ctrl WT (n=14) and cKO (n=13) mice before and after SNAP treatment. C-D, Quantification of spectral power differences in AuC (C) and FC (D) of Ctrl WT and cKO mice before and after SNAP treatment. Mice were first recorded at P27-P29 and then one or two days later following SNAP treatment. The enhanced high frequency gamma oscillations can be visually observed in both AuC and FC of cKO mice following SNAP treatment. Mixed-effects analysis controlling for the effect of movement revealed differences in the gamma range in FC of cKO SNAP mice after Fisher’s LSD post hoc test (*p<0.05, **p<0.01). The gamma band was further subdivided into low and high gamma revealing treatment differences in high gamma bands. E-F, Power coupling of different oscillation frequencies. Graphs show Pearson’s correlation (r) for Alpha frequency power coupling (E): AuC Alpha/FC Alpha, AuC Alpha/AuC Gamma, AuC Alpha/FC Gamma, FC Alpha/AuC Gamma, FC Alpha/FC Gamma; and Delta frequency power coupling (F): AuC Delta/FC Delta, AuC Delta/AuC Gamma, AuC Delta/FC Gamma, FC Delta/AuC Gamma, FC Delta/FC Gamma (*p<0.05; **p<0.01; Mixed-effects analysis, Fisher’s LSD post hoc test). G-H, Quantification of spectral power differences in AuC (G) and FC (H) of WT pre (n=43), *Fmr1* KO pre (n=33), WT vehicle (n=16), *Fmr1* KO vehicle (n=12), WT SNAP (n=11)and *Fmr1* KO SNAP (n=7). Mice were first recorded at P22-P28 and then one day later following vehicle or SNAP treatment. The gamma band was further subdivided into low and high gamma revealing genotype differences in low gamma bands. Enhanced gamma power was observed in SNAP treated KO compared to vehicle treated KO or KO pre groups in AuC and FC. I-J, Power coupling of different oscillation frequencies. Graphs show Pearson’s correlation (r) for Alpha frequency power coupling (I): AuC Alpha/FC Alpha, AuC Alpha/AuC Gamma, AuC Alpha/FC Gamma, FC Alpha/AuC Gamma, FC Alpha/FC Gamma; and Delta frequency power coupling (J): AuC Delta/FC Delta, AuC Delta/AuC Gamma, AuC Delta/FC Gamma, FC Delta/AuC Gamma, FC Delta/FC Gamma (*p<0.05; **p<0.01, ***p<0.001; ****p<0.0001; Mixed-effects analysis, Fisher’s LSD post hoc test). KO mice before or after vehicle treatment show dysregulated A/G and D/G power coupling compared to WT pre group within and between AuC and FC (red stars). The genotype difference is lost between SNAP-treated KO and WT pre groups. However, vehicle and SNAP treatment had negative effects in the WT group. Both WT vehicle and WT SNAP showed abnormal A/G and D/G coupling when compared to WT pre group (black stars).

As human EEG studies report impaired low/high frequency power coupling in FXS and ASD^70,71^, we next investigated cross-frequency power coupling within and between regions for Alpha/Gamma (A/G, **Figure 6E, Table S6**) and Delta/Gamma (D/G, **Figure 6F, Table S6**) frequencies in our model. Negative correlation in power coupling of AuC Alpha/AuC Gamma (A1G1) and AuC Alpha/FC Gamma (A1G2) observed in Ctrl WT and WT mice became positively correlated in cKO and global KO mice, but was normalized following SNAP treatment with no effect of vehicle treatment (**Figure 6E, 6I, Table S6**). While vehicle treatment aggravated Delta/Gamma coupling in AuC and FC of both WT and global *Fmr1* KO mice, SNAP treatment normalized it, potentially explained by the corresponding decrease in delta and increase in gamma power (**Figure 6J, Table S6**). Delta/Gamma coupling comparisons showed a similar trend in SNAP cKO mice (**Figure 6F, Table S6**).

Overall, these findings reveal that alpha/gamma and delta/gamma power coupling is likely restored to normal levels due to the elevation in high gamma power. As PV cells are known to generate and regulate gamma frequency oscillations^72–74^, the increase in high gamma power in cKO SNAP, but not Ctrl WT SNAP, may be explained by increased PV cell activity.

### Acute blockade of GAT3-mediated astrocytic GABA transport improves fidelity of temporal processing to the frequency-modulated sound chirp in high gamma range in both cKO and global *Fmr1* KO mice

Given that PV neuron activity is vital for timing fidelity of responses, we tested the hypothesis that the ability of the cortex to mount a consistent response to repeated time-varying stimuli would be impaired in astrocyte-specific cKO mice and rescued with SNAP treatment. Previous studies in rodents and humans have shown a reduction in inter-trial phase coherence (ITPC) response to up “chirp” stimulus in gamma frequency in FXS. The chirp is a 2-s broadband noise stimulus whose amplitude is modulated (100% depth) by a sinusoid of linearly increasing frequencies from 1-100 Hz. The ability of the neural generators to consistently phase lock to this time-varying signal is a measure of fidelity in temporal processing. After repeated chirp presentations, the ITPC was calculated across trials in the time X frequency domain using Morlet Wavelet analysis as previously described^22,67^. Grand average ITPC was calculated for each group, then control means were subtracted from experimental groups to show difference plots for AuC (**Figure 7A**) and FC (**Figure 7B**). For statistical comparisons, non-parametric cluster analysis was used to determine contiguous regions in the time X frequency domain that were statistically different between genotypes. We observe a significant decrease in ITPC at gamma frequencies, for low gamma in AuC and high gamma in FC of cKO mice. These data indicate that postnatal deletion of FMRP from astrocytes results in the development of gamma synchronization deficits, which are similar to the deficits observed in AuC (**Figure 7C**) and FC (**Figure 7D**) of global *Fmr1* KO mice. SNAP treatment restored responses to Ctrl WT levels for low gamma in AuC and high gamma in FC of cKO mice. In global *Fmr1* KO, SNAP treatment also significantly improves ITPC for high gamma frequencies in both AuC and FC, whereas vehicle treatment had no effect on high gamma. However, there was a vehicle effect on low gamma in AuC (**Figure 7E**) and FC (**Figure 7F**), indicating that improvements observed in low gamma may be attributed to vehicle effects.

**Figure 7.**
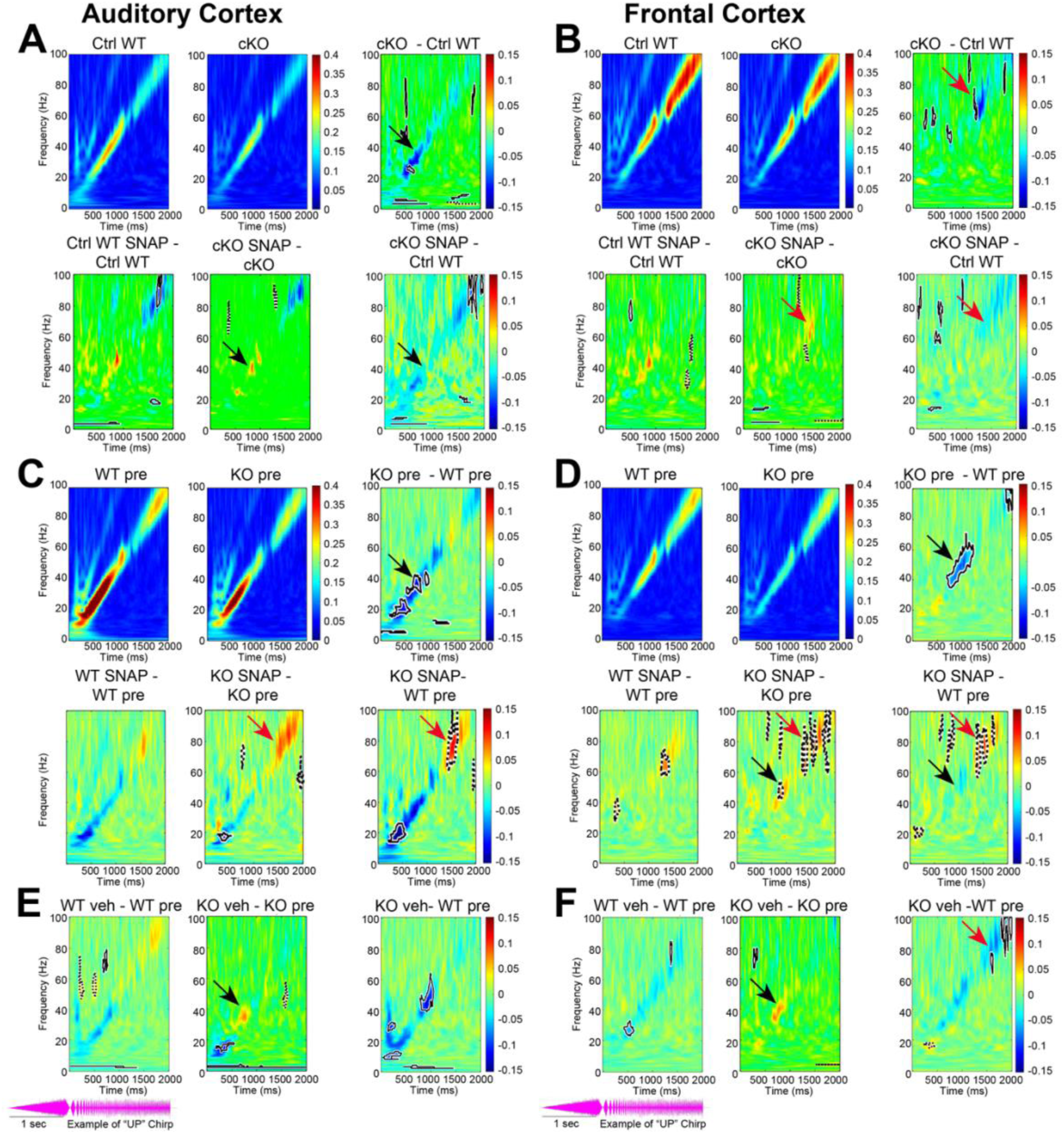
Acute blockade of GAT3-mediated astrocytic GABA transport improves fidelity of temporal processing to the frequency-modulated sound chirp in high gamma range in both cKO and global Fmr1 KO mice. The chirp stimulus (oscillogram shown at the bottom of this figure) is a 2 s broadband noise with amplitude modulated linearly by a frequency sweep with frequencies increasing from 1 to 100 Hz. To prevent the early response to stimulus onset, the chirp is preceded by a 1 s slow ramp of broadband noise. The ability of the cortical neural generators to follow this temporally dynamic stimulus is quantified by measuring the inter-trial phase coherence (ITPC, also known as phase locking factor). Trains of chirp stimuli were presented to each mouse 300 times. For each mouse, ITPC was measured to determine the degree of phase locking across trials. A-B, Grand average matrices were calculated for each genotype and treatment, and then Ctrl WT (n=12) average values were subtracted from cKO (n=13) average values (right panels) and pre-treatment values were subtracted from SNAP-treated values (bottom panels) for AuC (A) and FC (B). SNAP-treated cKO was also compared to Ctrl WT group. Blue areas indicate less phase locking, green areas no difference, and red areas more phase locking. Statistical cluster analysis reveals contiguous time x frequency regions that are significantly different between genotypes and treatments. Black solid contours (mean negative difference) and black dashed contours (mean positive difference) indicate clusters with significant differences. cKO mice express statistically significant decrease in ITPC at gamma frequencies (30-100 Hz, blue) that was restored to Ctrl WT levels following SNAP treatment. C-D, Grand average matrices of ITPC values were calculated for global KO (n=13) and WT (n=21) before (pre) and global KO (n=7) and WT (n=11) after SNAP treatment and subtracted to generate difference plots by genotype (right panels) and treatment (bottom panels) for AuC (C) and FC (D). Global KO mice express statistically significant decrease in ITPC at gamma frequencies (30-100 Hz, blue), with gamma ITPC restored to Ctrl WT levels in FC following SNAP treatment. E-F, Grand average matrices of ITPC values were calculated for global KO (n=20) and WT (n=22) before (pre) and global KO (n=12) and WT (n=16) after vehicle treatment and subtracted to generate difference plots by genotype (right panels) and treatment (bottom panels) for AuC (E) and FC (F). Vehicle treatment affected ITPC values in low gamma range without affecting high gamma.

Analysis of non-phase locked single trial power (STP) during the chirp stimulation in cKO SNAP mice showed a significant increase in gamma power in FC in the high gamma band compared to Ctrl WT mice (**Figure S4A-S4B**). The effects were different from the findings in global *Fmr1* KO mice, where background power was enhanced in KO compared to WT but the difference was lost in FC of SNAP treated KO mice (**Figure S4C-S4D**). The effects were specific in the SNAP-treated KO group and were not observed in vehicle-treated KO (**Figure S4E-S4F**). To examine effects of astrocyte-specific *Fmr1* deletion on habituation to repeated stimuli, we evaluated sound-evoked responses to trains of brief (100 ms) broadband noise stimuli (10 stimuli per train, 65-70 dB SPL, 100 repetitions of each train), delivered at a non-habituating repetition rate (0.25 Hz) and a habituating rate of sound presentation (4 Hz)^22,75^. We measured the amplitude and latency of P1, N1, and P2 waves, and N1 habituation for each repetition rate (**Figure S5**). Post-hoc analysis showed no differences between groups on amplitude, latency, or habituation, however, we observed an effect of treatment on latency of P1 waves in FC and P2 waves in AC (**Figure S5I-S5J**).

Overall these findings reveal a specific effect of GAT3-mediated GABA transport on fidelity of temporal processing to the chirp sound in high gamma range. In line with interpretation of baseline EEG data showing SNAP modulation of high gamma power, its effects on high gamma frequencies during chirp presentation can be also explained by disinhibition of PV cell activity.

### Postnatal deletion of *Fmr1* in astrocytes leads to increased locomotor activity and aberrant exploratory behavior in cKO mice

Neocortical imbalance of excitation and inhibition as a result of PV cell hypofunction and impaired inhibition is also observed in several ASD mouse models^76^ and may underlie ASD-like behaviors, such as enhanced anxiety and hyperactivity. Therefore, we tested Ctrl WT, heterozygous (Het, female) and cKO mice for hyperactivity, exploratory behavior, and anxiety-like behaviors in open field test (**Figure 8, Table S7**) and elevated plus maze (**Figure 8, Table S7**). An additional cohort of cKO males were tested 4 h following SNAP treatment. cKO mice demonstrated increased locomotor activity in the open field test with significantly higher average speed compared to Ctrl WT and Het mice throughout all 10 min of the test (**Figure 8B-8C, Table S7**). We found that during the first 5 min of the test, when mice habituated to the arena, all groups spent significantly more time in thigmotaxis compared to open field. However, while Ctrl WT mice still showed preference to the thigmotaxis during the second 5 min of the test (behaviors considered to be normal as mice usually avoid open spaces); Het and cKO mice spent comparable time in both areas, indicating greater exploratory behavior (**Figure 8D, Table S7**). Pre-treatment with SNAP rescued locomotor activity (**Figure 8B-8C, Table S7**) and significantly reduced exploratory behaviors (**Figure 8D, Table S8**). No changes in anxiety-like behaviors were observed in any groups compared with Ctrl WT mice, indicated by time spent in open arms of elevated plus maze (**Figure 8E, Table S8**). Our findings establish that postnatal deletion of FMRP from astrocytes leads to increased locomotor activity and exploratory behavior, and pharmacological blockade of astrocyte GABA transporter GAT3 rescues these behaviors.

**Figure 8.**
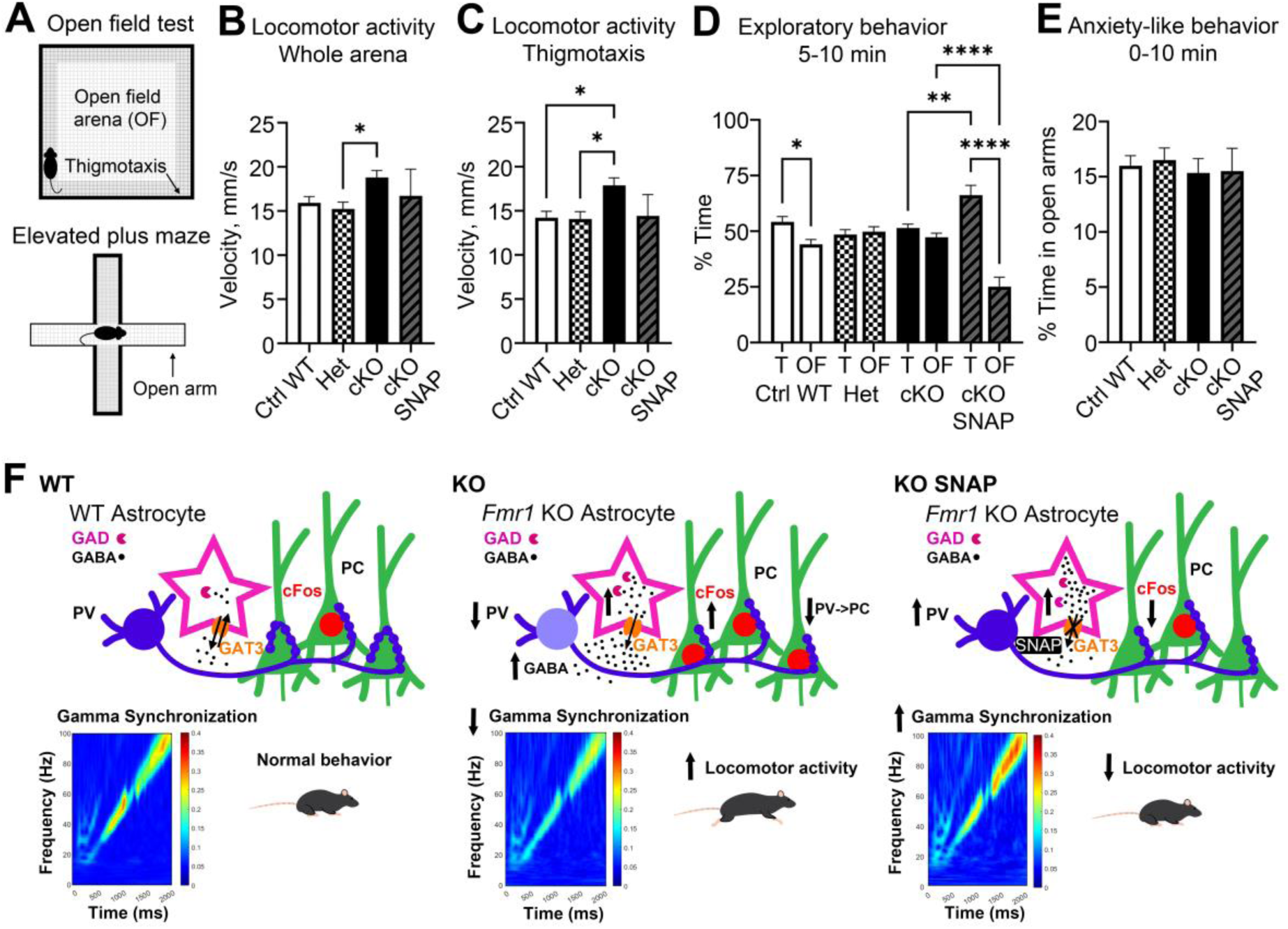
Postnatal deletion of Fmr1 in astrocytes leads to increased locomotor activity and aberrant exploratory behavior in cKO mice. A-D, Graphs demonstrate the performance of Ctrl WT (n=30), Het (n=36), cKO (n=43), and cKO SNAP (n=10) mice in the open field. Locomotor activity is measured by speed in whole arena (B) and in thigmotaxis (C). Exploratory behavior is measured by percentage of time in open field compared to thigmotaxis (D). Graphs show mean ± SEM (*p<0.05, **p<0.01, ****p<0.0001; one-way ANOVA (B,C), two-way ANOVA (D), Tukey’s post hoc test (b-d)). cKO male mice exhibit higher locomotor activity than Ctrl WT and Heterozygous (Het) female mice throughout the full 10 min of the test. cKO and Het mice exhibit more exploratory behavior than Ctrl WT and show no preference between thigmotaxis and open space in the second 5 min of the test. Pharmacological blockade of GAT3 using SNAP-5114 in cKO normalizes locomotor activity and exploratory behaviors. E, Graph demonstrates the performance of Ctrl WT (n=30), Het (n=36), cKO (n=43), and cKO SNAP (n=10) mice in the elevated plus maze. Anxiety-like behavior is measured by percentage of time in open arms. Graph shows mean ± SEM (p=0.05; one-way ANOVA, Tukey’s post hoc test). No changes in anxiety-like behaviors are observed. F, Schematic depiction of phenotypes observed following deletion of *Fmr1* from astrocytes during P14-P28 postnatal developmental period compared to Ctrl WT. cKO mice exhibit increased GABA synthesis (pink), enhanced reverse GABA transport, decreased PV-positive (blue) perisomatic innervation of PC (green), reduced PV levels (blue) and enhanced cFos activity (red). cKO mice also show reduced chirp-evoked gamma synchronization and increased locomotor activity. Acute SNAP treatment normalizes PV levels, overall cFos activity, improves chirp-evoked gamma synchronization and reduces behavioral hyperactivity.

## Discussion

Cortical hyperexcitability is a feature of ASDs that is not fully understood with potential to reveal novel targets for treatment of symptoms affecting quality of life. The main findings of this study support a novel, non-neuronal mechanism of cortical hyperexcitability in Fragile X Syndrome involving dysregulation of GABA synthesis and transport in astrocytes. We demonstrate that deletion of *Fmr1* from astrocytes during the postnatal developmental period of cortical circuit refinements results in (1) excess GABA synthesis and release from astrocytes; (2) which suppresses PV expression and inhibitory connectivity; (3) leading to excess cell activation; (4) impaired cortical responses to stimuli; and (5) behavioral hyperactivity; all of which are rescued by pharmacologically blocking astrocyte GABA transporter GAT3 in adolescent mice.

Detecting increased GABA levels and its synthesizing enzyme GAD67/67 in human astrocytes differentiated from FXS patient-derived iPSC lines allows us to draw direct comparisons with clinical features observed in the donors. For example, the cell line displaying the highest GABA and GAD65/67 levels (HEL100.2) was derived from an FXS individual diagnosed with epilepsy, supporting our interpretation that elevated astrocytic GABA may contribute to excitation/inhibition imbalance. Our finding of elevated GABA synthesizing enzyme GAD65/67 levels are also consistent with previously published LC-MS/MS proteomics data showing increased GAD65/67 protein in *Fmr1* KO astrocytes, potentially explained by FMRP regulation of *gad2 gene* translation through a miR-31-5p mechanism^77^. The possibility of reverse GABA transport from astrocytes concerns a topic of prior debate, as GATs are proposed to primarily uptake GABA. However, studies suggest GAT3 transporters can operate in reverse mode in the cortex, releasing GABA when intracellular GABA concentration is sufficiently high or extracellular GABA concentration is sufficiently low^59,61,78,79^. GATs carry a net positive charge, with 2Na^+^ and 1Cl^-^ co-transported with one GABA molecule, and can reverse when membrane potential surpasses the GAT reversal potential or with high intracellular Na^+^ concentration, observed with glutamate uptake via EAAT1/2^78,80^. It is most likely that FXS astrocytes are up-taking excess glutamate as a result of increased cortical excitability, while synthesizing more GABA from glutamate with GAD65/67 leading to reverse GABA transport from astrocytes. However, excess extracellular GABA most likely affects activity of inhibitory interneurons, with astrocytic GABA release promoting circuit hyperexcitability, resulting in more glutamate for astrocyte uptake. Interestingly, inhibiting NKCC1, highly expressed in developing glia, showed beneficial effects on cortical hyperexcitability in FXS through an unexplained mechanism^81^, potentially explained by lowered sodium concentration in astrocytes preventing GAT reversal.

Our model displays fewer GABAergic and PV-positive synapses, providing structural evidence of excitation/inhibition imbalance. A recent study suggests that GABA-receptive microglial cells can negatively regulate perisomatic PV-expressing inhibitory synapses during the same P15-P30 developmental period in the somatosensory and visual cortex^46^. It is possible that astrocytic GABA release may trigger excessive pruning of inhibitory synapses in the auditory cortex as well. Moreover, previous studies support the role of *Fmr1* KO astrocytes in synaptic changes in neurons that can be restored by developmental *Fmr1* re-expression in astrocytes^37,82,83^. PV cell hypofunction is proposed to contribute to cortical hyperexcitability in FXS and other ASDs^4,54–58^. Our observation of lower PV levels and increased proportion of low PV expressing cells in the auditory cortex is in line with previous work showing impaired PV cell function in the visual cortex of *Fmr1* KO, but does not replicate diminished PV cell density in somatosensory cortex^21,84^, suggesting an astrocytic mechanism governs PV cell hypofunction, but not PV density. This mechanism can be explained by astrocytic excess extracellular GABA suppressing PV cells via tonic inhibition. Cell type-specific effects have been observed with tonic inhibition, disproportionately inhibiting PV and VIP cells over SST or pyramidal cells in somatosensory cortex^62,63^. Lessening extracellular GABA levels by blocking GAT3 was sufficient to recover PV expression and reduce overall cortical cell excitability in our model, a mechanism that was previously described from findings in wildtype rat neocortex, where GAT3 blockade increased inhibitory postsynaptic potentials and decreased excitatory synaptic potentials in cortical neurons^59^.

To observe whether cellular changes affected cortical activity and stimulus-evoked responses, we used clinically-relevant EEG recordings. Similarly to findings in FXS human individuals^68^, we observed abnormal alpha/gamma power coupling at rest. Negative alpha/gamma coupling is proposed to control top-down regulation and gating of sensory processing^85,86^. GAT3 inhibition resulted in a significant increase in high gamma power, potentially as a result of enhanced PV cell activity. This is consistent with previous studies in mice reporting acute modulation of PV cell activity affects cortical oscillations in high gamma range^72–74^. Replicating previous observations in FXS humans and *Fmr1* KO mice, we observed impairments in temporal fidelity of temporal processing to the frequency-modulated sound chirp in high gamma range in both AuC and FC, suggesting cortical circuit deficits^67,87^. The deficits were alleviated by GAT3 inhibition in both astrocyte-specific and global *Fmr1* KO mice, consistent with the recovery in PV cell activity, as PV cell function is linked to cortical gamma oscillations. Increasing PV activity has been shown to restore visual task performance and temporal fidelity in *Fmr1* KO mice, suggesting that the effects of SNAP in our model support the role of PV cells in improved sensory processing^21^. Interestingly, altered astrocytic GABA transport induced deficits in population encoding of visual stimuli, further supporting the role of astrocytes in regulating temporal coherence of neuronal responses^88^. Our data show that the loss of FMRP from astrocytes leads to impaired gamma phase–locking to sound in both auditory and frontal cortices, which can be rescued through blockade of astrocyte GAT3-mediated GABA release, underscoring the critical role for astrocytes in pathogenesis and potential therapeutic interventions in FXS.

We observed increased locomotor activity and abnormal exploratory behavior in astrocyte-specific *Fmr1* cKO mice. Previous studies from other labs have shown astrocyte-specific *Fmr1* KO under the same GFAP promoter results in impaired motor skill learning in adulthood^89^. Astrocytes are known to modulate cognitive functions and complex animal behaviors, such as emotion, motor activity, memory formation, and sensory processing^90^. Not only have we shown a role for astrocytes in biasing excitation/inhibition balance in FXS circuit development, but correction of elevated locomotion and abnormal exploratory behavior with SNAP further demonstrates astrocytes actively modulate FXS-related behaviors, and most likely through PV cell regulation.

Together, our findings reveal a novel mechanism linking alterations in astrocyte GABA transport to the development of cortical hyperexcitability and abnormal cortical processing, demonstrating a non-neuronal mechanism for inhibitory circuit regulation in FXS, and provide a new direction for relieving symptoms of FXS and potentially other ASDs.

## Methods

### Ethics Statement

All mouse studies were done according to National Institutes of Health and Institutional Animal Care and Use Committee at the University of California Riverside (approval #20190015 and #20190029) guidelines; animal welfare assurance #A3439-01 is on file with the Office of Laboratory Animal Welfare. Mice were maintained in an Association for Assessment and Accreditation of Laboratory Animal Care-accredited facility under 12 h light/dark cycle and fed standard mouse chow.

### Human astrocytes

Human forebrain astrocytes were produced as described previously^45^ from male FXS (SO02B, SO04A, SO10c3, HEL70.3, HEL70.6, HEL100.2, and HEL69.6) and control (V36C, V44D, SC176, HEL23.3, HEL24.3, and P04) human iPSC lines reported in several previous studies^91–93^. The research using human cells was approved by the Ethics Committee of the Hospital District of Helsinki and Uusimaa, Finland and the UCR Stem Cell Research Oversight Committee (approval #SC-20190003). Briefly, iPSCs were maintained on Matrigel-coated plates in mTeSR medium (STEMCELL Technologies, Vancouver, BC, Canada). For differentiation, 80% confluent cells were passaged with ReLeSR (STEMCELL Technologies) and plated on low-attachment 6-well plate in mTeSR with 20 ng/ml human basic fibroblast growth factor (bFGF2; STEMCELL Technologies) and 20 µM Rho kinase inhibitor (Tocris Bioscience, Bristol, UK). On the following day, medium was changed to neuronal induction medium [Advanced Dulbeccós Modified Eagle Medium (DMEM)/F12, 1x N2, 2 mM L-glutamine, 1x non-essential amino acids (NEAA), and 1x penicillin-streptomycin (P-S) (all from Thermo Fisher Scientific, Waltham, MA, USA)] supplemented with 0.1 µM LDN-193189 (ABMole, Houston, TX, USA), 1 µM cyclopamine (APExBIO, Houston, TX, USA), 10 µM SB-431542 (Selleck Chemicals, Houston, TX, USA), and 0.5 µg/ml Dickkopf-related protein 1 (DKK1; RayBiotech, Peachtree Corners, GA, USA). At 12 DIV, medium was replaced by neuronal induction medium supplemented with 20 ng/ml brain-derived neurotrophic factor (BDNF; Cell Guidance Systems, Cambridge, UK), and at 30 DIV by neurosphere medium (Advanced DMEM/F12, 1x B27 -RA, 1x L-glutamate, 1x NEAA, 1x P-S [all from Thermo Fisher Scientific]) supplemented with 20 ng/ml bFGF2 and 20 ng/ml epidermal growth factor (EGF; Thermo Fisher Scientific). Growth factors were added three times a week and spheres were dissociated approximately once per week manually. At 60 DIV, spheres were dissociated and progenitors plated on poly-ornithine (Sigma-Aldrich, St. Louis, MO, USA) /laminin-coated (Thermo Fisher Scientific) culture plates in neurosphere medium supplemented with 20 ng/ml ciliary neurotrophic factor (CNTF; Peprotech). Cells were passaged with Trypsin-EDTA (0,05%) (Thermo Fisher Scientific) approximately once per week and seeded at 20 000 cells/cm2. After 75 DIV, cells had acquired astrocyte morphology and they were maintained on Matrigel-coated culture plates. As shown previously, FXS astrocytes did not express FMRP.

### Determination of GABA concentration in human astrocytes

The cell samples were homogenized in 0.2 ml of homogenization solution consisting of six parts 0.2 M HCLO4 and one part antioxidant solution containing oxalic acid in combination with acetic acid and l-cysteine99. The homogenates were centrifuged at 20,800 g for 35 min at 48°C. The supernatant was removed to 0.5 ml Vivaspin filter concentrators (10,000 MWCO PES; Vivascience AG, Hannover, Germany) and centrifuged at 8600 g at 4 °C for 35 min. The concentrations of GABA were analyzed with HPLC equipped with a fluorescence detector100. The HPLC system was consisted of a solvent delivery pump (Jasco model PU-1580 HPLC Pump, Jasco International Co, Japan) connected to an online degasser (Jasco 3-Line Degasser, DG-980-50) and a ternary gradient unit (Jasco LG-1580-02), a refrigerated autosampler (Shimadzu NexeraX2 SIL-30AC Autosampler, Shimadzu Corp), an analytical column (Kinetex C-18 5 µm, 4.60 x 50 mm, Phemomenex Inc) protected by a 0.5-mm inlet filter and thermostated by a column heater (CROCO-CIL, Cluzeau Info-Labo, France), and a fluorescence detector (Jasco Intelligent Fluorescence Detector model FP-1520). The wavelengths of the fluorescence detector were set to 330 (excitation) and 450 (emission). The mobile phase consisted of 0.1 M NaH2PO4 buffer (Merck), pH 4.9 (adjusted with Na2HPO4), 20 % (v/v) acetonitrile (Merck), and the flow-rate was 1.2 mL/min. Automated sample derivatization was carried out using the autosampler at 8 °C. The autosampler was programmed to add 6 µL of the derivatizing reagent (3 µL of mercaptoethanol ad 1 mL of o-phtaldialdehyde) to 15 µL of a microdialysis sample, to mix two times, and to inject 20 µL onto the column after a reaction time of 1 min. The chromatograms were processed by AZUR chromatography data system software (Cromatek, Essex, UK).

### Primary astrocyte culture

Primary astrocyte cultures were prepared from P1-P3 cortex of WT and KO littermates male mice (RRID: IMSR_JAX:003025) as described^94,95^ with minor modifications. Briefly, cortex from both hemispheres was dissected, cut into 1-2 mm pieces, placed into a separate tube (cortices of one mouse per tube) and treated with a papain/DNase I (0.1 M PBS/0.1% BSA/ 25 mM glucose/5% papain/1×DNase I [Sigma, catalog #D5025-15K]) solution for 20 min at 37°C. Cells were mechanically dissociated and plated on T25 flask (one brain per flask) coated with poly-D-lysine (0.05 mg/mL; Sigma, catalog #P7886) in Dulbecco’s Modified Eagle Medium (Gibco, catalog #11995-065) containing 1% penicillin-streptomycin (Gibco, catalog #15140-122) and supplemented with 10% fetal bovine serum (FBS; Gibco, catalog # 10438-018) under 10% CO2 atmosphere at 37°C. To achieve purified astrocyte cultures (>95% astrocytes) cells were shaken after 4-5 days in vitro (DIV) for 30 min at 180 rpm (to remove microglia) and then 4 h at 240 rpm (to remove oligodendrocyte precursor cells). After shaking, the medium was removed, and cells were washed twice with 0.1 M PBS (pH 7.4). Astrocytes were then treated with 0.25% trypsin/EDTA solution (Gibco, catalog #25200-056) for 2 min at 37℃ and plated on poly-D-lysine-coated T75 flask in DMEM containing 10% FBS. Once confluent, astrocytes were trypsinized and plated on coverslips in 24-well plates at a density of 35,000 cells per well and T75 flask (all remaining cells). Astrocytes were cultured for 2 days before being fixed for immunocytochemistry (in 24-well plate) or until confluent for Western blot (in T75 flask). For immunostaining, astrocytes were fixed with 2% paraformaldehyde for 30 min at room temperature, washed three times with PBS 0.1 M and stained as described below.

For Western blot, cells were collected from T75 flask and lysed in the lysis buffer (25 mM Tris-HCl, 150 mM NaCl, 5 mM EDTA, 1% Triton X-100, 1 mM sodium pervanadate, and protease inhibitor cocktail [1:100, Sigma, catalog #P8340]) at 4°C for 30 min. Cell lysates were cleared by centrifugation at 13,500 rpm for 20 min at 4°C. Samples were further concentrated using protein concentrators (Pierce, catalog #88513) to achieve 10x concentration and processed for Western blot.

### Mice

To achieve specific *Fmr1* deletion in astrocytes, two different mouse lines were generated. In group 1, ERT2-Cre*^GFAP^* (B6.Cg-Tg(*GFAP*-cre/ERT2)505Fmv/J RRID: IMSR_JAX:012849) male mice were crossed with *Fmr1*^flox/flox^ female mice (generated in and obtained from the laboratory of Dr. David Nelson (Baylor College of Medicine, Houston, Texas^96^) to obtain ERT2-Cre*^GFAP^Fmr1*^flox/y^ condition KO (cKO) male mice. ERT2-Cre*^GFAP^* mice were designated as control WT (Ctrl WT). In group 2, ERT2-Cre*^GFAP^* mice and ERT2-Cre*^GFAP^Fmr1*^flox/flox^ mice were crossed with Rosa-CAG-LSL-tdTomato reporter mice (CAG-tdTomato; RRID: IMSR_JAX:007909) to generate tdTomato-ERT2-Cre*^GFAP^* Ctrl WT and tdTomato-ERT2-Cre*^GFAP^Fmr1*^flox/y^ cKO mice, respectively, allowing for tdTomato expression in astrocytes and analysis of *Fmr1* levels. In group 3, global *Fmr1* KO mice on C57BL/6 background (Jackson Laboratories, JAX:003025) and their wild-type (WT) counterparts (Jackson Laboratories, JAX: 000664) were bred as littermates (by crossing heterozygous *Fmr1* KO females and wild-type or KO males). Real-time PCR-based analysis of genomic DNA isolated from mouse tails was used to confirm genotypes by Transnetyx.

In groups 1 and 2, Ctrl WT and cKO mice received tamoxifen at P14 intraperitoneally (0.5 mg in 5 mg/ml of 1:9 ethanol/sunflower seed oil solution) once a day for 5 consecutive days, and analysis was performed at P28 (see “Experimental Timeline” in Figure 3a). Group 1 was used for immunohistochemistry, Western blot, qRT-PCR, EEG, and behavioral analysis; group 2 was used for immunohistochemical analysis of FMRP levels and group 3 was used for EEG study.

### Methods overview

As *Fmr1* is expressed in astrocytes during postnatal development^37,97^ and we observed major changes in PV cell development during postnatal day (P)14-P21, we used ERT2-Cre*^GFAP^* line to delete *Fmr1* from astrocytes during the same P14-P21 period by administering tamoxifen daily from P14 to P18. To confirm deletion of FMRP in astrocytes, we examined expression of FMRP in the cortex of P28 mice using immunostaining. Once deletion was established, all subsequent immunohistochemistry, Western blot, qRT-PCR, and EEG experiments were performed in P28 male mice. For behavioral testing, P28 male and female mice were used. At the ages studied here, accelerated hearing loss in the C57bl/6J mice is minimal, and because all group comparisons are within-strain and age-matched, hearing loss is unlikely to factor into the phenotypes described^98^.

At P28, we analyzed the effects of astrocyte specific *Fmr1* deletion on gene expression in the cortex using qRT-PCR. Biochemical measurements of GAT3 and GABA_A_ receptors were done in the cortex of cKO and Ctrl WT mice using Western blot. Lastly, immunohistochemistry was used to analyze inhibitory and PV-positive synapses by measuring vGAT/Gephyrin and PV/vGAT co-localization in the cortex of cKO and Ctrl WT mice, and used (1) to assess GABA levels in astrocytes, (2) to determine PV levels, and (3) to examine hyperexcitability by quantifying density of cFos-positive cells in the AuC of Ctrl WT vehicle, cKO vehicle, Ctrl WT SNAP, and cKO SNAP mice. EEG recordings were performed in awake, freely moving mice to determine the effects of postnatal FMRP deletion in astrocytes on neural oscillations in the auditory and frontal cortex at baseline and in response to sound at P28, and repeated 1 or 2 days later 4 h post SNAP-5114 intraperitoneal (ip) injection^67^ or vehicle treatment. We also analyzed the effects of SNAP and vehicle in P22-P28 WT and global *Fmr1* KO mice using EEG recordings. Finally, we used established behavioral tests to examine hyperactivity and anxiety-like behaviors in Ctrl WT, Het, and cKO mice.

### Immunofluorescence

Four hours following injection of either SNAP-5114 (25 mg/mL) or vehicle (10%DMSO in saline, 10mL/kg) ip injection, P28 male Ctrl WT and cKO mice were euthanized with isoflurane and perfused transcardially first with cold phosphate-buffered saline (PBS, 0.1 M) to clear out the blood and then with 4% paraformaldehyde (PFA) in 0.1M PBS for fixation. Brains were removed and post-fixed for 2 h in 4% PFA. 100 μm brain slices were obtained using a vibratome (5100 mz Campden Instruments). Auditory cortex (AuC) and frontal cortex (FC) were identified using the brain atlas^99^. For each brain, an average of 8 brain slices were collected per region.

Immunostaining for FMRP was performed using antigen retrieval methods, as previously described^22^. Briefly, 40 μm brain slices obtained from tdTomato-ERT2-Cre*^GFAP^Fmr1*^flox/y^ cKO and tdTomato-ERT2-Cre*^GFAP^* Ctrl WT male mice were stained overnight with mouse anti-FMRP (1:100; Developmental Studies Hybridoma Bank, catalog #2F5-1-s, RRID: AB_10805421). Secondary antibody was donkey anti-mouse Alexa 488 (4 μg/mL; Molecular Probes, catalog# A-21202, RRID: AB_141607). Slices were mounted on slides with Vectashield mounting medium containing DAPI (Vector Laboratories, catalog # H-1200, RRID:AB_2336790) and Cytoseal.

Immunostaining in 100 μm brain slices containing AuC and FC was performed as previously described^22^. For slices obtained from ERT2-Cre*^GFAP^Fmr1*^flox/y^ cKO or ERT2-Cre*^GFAP^* Ctrl WT male mice, astrocyte cell bodies were identified by immunolabeling against Glutamine Synthetase using rabbit anti-Glutamine Synthetase antibody (1:500, Sigma-Aldrich Cat# G2781, RRID:AB_259853), and GABA immunoreactivity was detected by immunostaining with guinea pig anti-GABA antibody (1:500, Abcam, catalog # ab17413, RRID:AB_443865). Inhibitory presynaptic sites were detected by immunolabeling against vesicular GABA transporter (vGAT) using rabbit anti-vGAT antibody (1:100, Synaptic Systems, catalog # 131002, RRID:AB_887871). Inhibitory postsynaptic sites were detected by immunolabeling against gephyrin using mouse anti-gephyrin antibody (1:500, Synaptic Systems, catalog # 147111, RRID:AB_887719). Parvalbumin-positive sites were detected by immunolabeling against parvalbumin using mouse anti-parvalbumin antibody (Millipore Cat# MAB1572, RRID:AB_2174013). Active cells were detected by immunolabeling against cFos using rabbit anti-cFos antibody (1:500, Cell Signaling Technologies, catalog # 2250, RRID:AB_ 224721). Secondary antibodies used were as follows: AlexaFluor-488-conjugated goat anti-guinea pig IgG (4 µg/ml, Invitrogen, catalog # A-11073, RRID:AB_2534117), AlexaFluor-488-conjugated donkey anti-mouse IgG (4 µg/ml, Invitrogen, catalog # A-21202, RRID:AB_141607), FITC-conjugated goat anti-mouse IgG (4 µg/ml, Invitrogen, catalog # 31569, RRID:AB_228306), and donkey anti-rabbit Alexa Fluor 594 (4 µg/ml, Invitrogen, catalog #A-21207, RRID:AB_141637). Slices were mounted on slides with Vectashield mounting medium containing DAPI (Vector Laboratories, catalog # H-1200, RRID:AB_2336790) and Cytoseal.

Immunostaining of fixed primary astrocytes was performed as follows. Cells were blocked for 1 h at room temperature with 5% NDS in 0.1% Triton X-100 in 0.1 M PBS and then immunolabeled with guinea pig anti-GABA antibody (1:1000, Abcam, catalog # ab17413, RRID:AB_443865) and rabbit anti-GS (1:1000, Sigma-Aldrich Cat# G2781, RRID:AB_259853) antibody in blocking buffer overnight at 4°C. Coverslips then were washed 3×10 min with 1% NDS in 0.1 M PBS at room temperature followed by the incubation with goat anti-guinea pig Alexa Fluor 488 (4 µg/ml, Invitrogen, catalog #A-11073, RRID:AB_2534117) and donkey anti-rabbit Alexa Fluor 594 (4 µg/ml, Invitrogen, catalog #A-21207, RRID:AB_141637) for 2 h at room temperature. Coverslips were washed 3×10 min with 0.1 M PBS, mounted on slides with Vectashield mounting medium containing DAPI (Vector Laboratories, #H1200) and sealed with Cytoseal.

Immunostaining of fixed human astrocytes was performed as follows. Cells were blocked for 1 h at room temperature with 5% NDS in 0.1% Triton X-100 in 0.1 M PBS and then immunolabeled with mouse anti-S100b antibody (1:500, Sigma-Aldrich, Cat# S2532, RRID:AB_477499) and rabbit anti-GFAP (1:500; Cell Signaling, catalog #12389, RRID:AB_2631098) antibody in blocking buffer overnight at 4°C. Coverslips then were washed 3×10 min with 1% NDS in 0.1 M PBS at room temperature followed by the incubation with donkey anti-mouse AlexaFluor 488 (4 µg/ml, Invitrogen, catalog # A-21202, RRID:AB_141607) and donkey anti-rabbit Alexa Fluor 594 (4 µg/ml, Invitrogen, catalog #A-21207, RRID:AB_141637) for 2 h at room temperature. Coverslips were washed 3×10 min with 0.1 M PBS, mounted on slides with Vectashield mounting medium containing DAPI (Vector Laboratories, #H1200) and sealed with Cytoseal.

### Image Analysis

Confocal images of coronal brain slices containing superficial layers of AuC and FC or all layers of AuC were taken with an SP5 confocal laser-scanning microscope (Leica Microsystems) as previously described with modifications^100,101^. Briefly, high-resolution optical sections (1024 × 1024 pixels format) were captured with a 40× (1.2 NA) and 1× zoom at 1 μm step intervals to assess FMRP immunoreactivity. Confocal images of GABA in astrocytes were taken using a 40× objective (1.2 NA), and 1× zoom at high resolution (1024 × 1024 pixels format) with a 1 μm interval. Confocal images of synaptic puncta were taken using a 40× objective (1.2 NA), and 1× zoom at high resolution (1024 × 1024 pixels format) with a 0.5 μm interval. Confocal images of PV cells were taken using a 10× objective (0.45 NA), and 1× zoom at high resolution (1024 × 1024 pixels format) with a 1 μm interval. Confocal images of cFos cells were taken using a 10× objective (0.45 NA), and 1× zoom at high resolution (1024 × 1024 pixels format) with a 1 μm interval. For GABA analysis in primary astrocyte cultures, images were captured using a Nikon Eclipse TE2000-U inverted fluorescent microscope with a 20× air objective and a Hamamatsu ORCA-AG 12-bit CCD camera using Image-Pro software. For analysis, 20 images were collected per animal (10 images per coverslip, 2 coverslips/animal, 5-6 animals per group). All images were acquired under identical conditions and processed for analysis as follows: (1) For analysis of FMRP immunoreactivity in astrocytes, astrocytes were visualized with tdTomato. Using ImageJ, cell areas were outlined using selection tool, then cell area, integrated fluorescent intensity, and mean intensity were measured for FMRP in each astrocyte. Statistical analysis was performed with unpaired t-test using GraphPad Prism 10 software (RRID:SCR_002798). (2) For analysis of GABA immunoreactivity in Glutamine

Synthetase (GS) labeled astrocytes, z stacks containing high-resolution optical serial sections (1024 × 1024 pixels format) taken at 1 μm intervals were compressed into max projections on ImageJ. Then, astrocytic cell bodies in superficial layers of AuC and FC were selected and outlined using selection tool. These ROIs were saved in the ROI manager and used to measure area and perform analysis of integrated fluorescent intensity and mean intensity of GABA in each astrocyte. Statistical analysis was performed with two-way ANOVA, Fisher’s LSD post hoc test using GraphPad Prism 10 software (RRID:SCR_002798).

(3) For analysis of vGAT/Gephyrin colocalization, three-dimensional fluorescent images were created by the projection of each z stack containing 5-10 high-resolution optical serial sections (1024 × 1024 pixels format) taken at 0.5 μm intervals in the X-Y plane. Using Neurolucida, vGAT puncta over 0.15 μm^3^ in size were rendered, followed by rendering gephyrin puncta over 0.11 μm^3^ in size that were within 1 μm of vGAT puncta. (4) For analysis of vGAT/PV colocalization, three-dimensional fluorescent images were created by the projection of each z stack containing 5-10 high-resolution optical serial sections (1024 × 1024 pixels format) taken at 0.5 μm intervals in the X-Y plane. Using Neurolucida, vGAT puncta over 0.15 μm^3^ in size were rendered, followed by rendering PV puncta 0.15 μm^3^ in size that were within 0.5 μm of vGAT puncta. If the images contained background VGAT somatic staining, puncta rendered in the soma were removed prior to rendering PV boutons. Puncta rendering of vGAT/Gephyrin and vGAT/PV was conducted using Neurolucida 360 software (MicroBrightField RRID:SCR_001775). Puncta renderings were analyzed using Neurolucida Explorer software (MBF Bioscience, RRID:SCR_017348) which measured total number of puncta and the number of puncta with at least 10% colocalization. Total and colocalized puncta counts were normalized to the 3D volume.

Statistical analysis was performed with two-way ANOVA, Fisher’s LSD post hoc test using GraphPad Prism 10 software (RRID:SCR_002798). (5) For analysis of PV levels and PV cell density, z stacks containing high-resolution optical serial sections (1024 × 1024 pixels format) taken at 1 μm intervals were compressed into average projections, auto-adjusted, background subtracted, thresholded using isodata, converted to a binary image, despeckled, and watershed on ImageJ. Analyze particles was used to generate ROIs of PV cells and ROIs detecting artifacts were manually removed. Remaining ROIs were counted and normalized to area for density measurements and were applied to a max projection for measurement of mean intensity per cell. Statistical analysis was performed with two-way ANOVA, Fisher’s LSD post hoc test using GraphPad Prism 10 software (RRID:SCR_002798).

(6) For analysis of cFos cell density, z stacks containing high-resolution optical serial sections (1024 × 1024 pixels format) taken at 1 μm intervals were compressed into max projections, thresholded using isodata, converted to a binary image, and despeckled on ImageJ. ComDet plugin v0.5.6 was used to count puncta. Total puncta count were normalized to area for density measurements. Statistical analysis was performed with two-way ANOVA, Fisher’s LSD post hoc test using GraphPad Prism 10 software (RRID:SCR_002798). (7) For analysis of GABA immunoreactivity in Glutamine Synthetase (GS) labeled astrocytes, primary cultured astrocytes were selected and outlined using selection tool in ImageJ. These ROIs were saved in the ROI manager and used to measure area and perform analysis of integrated fluorescent intensity and mean intensity of GABA in each astrocyte. Statistical analysis was performed with unpaired t-test using GraphPad Prism 10 software (RRID:SCR_002798). Data represent mean ± SEM.

### RNA Extraction

Total RNA was extracted from the AuC and FC of Ctrl WT and cKO mice (n=4-5 per group) using the RNeasy kit (Qiagen, Valencia, CA, USA) method. First-strand complementary DNA (cDNA) was generated using the High-Capacity cDNA Reverse Transcription Kit (Catalog #4368814, Applied Biosystems), and reverse transcription of total RNA (500 ng) was performed for 120 min at 37°C. All surfaces for tissue collection and processing were sanitized using 70% ethanol and then treated with an RNAse inhibitor (RNaseZap, Thermo Fisher Scientific, catalog # AM9780, Waltham, MA, USA) to maintain integrity of isolated RNA. cDNA was utilized for qRT-PCR.

### Quantitative Real Time PCR Analysis

qRT-PCR was carried out using SsoAdvanced™ Universal SYBR® Green Supermix (Biorad catalog # #1725271, Hercules, CA, USA) with the following primers for mouse genes: *Slc6a11*, *Gabra1*, *Gabra3*, *Gabrg2*, *Gabra5*; the following primers were used for human genes: *GFAP*, *S100b*, *Slc1a3, and Fmr1* (see Table S1 for sequences of primers used for qRT-PCR). *GAPDH* was selected as a housekeeping gene and no changes in its expression were found across genotype. Relative quantification using the delta-delta (2−ΔΔCq) method was used to compare changes in gene expression between cKO mice and Ctrl WT mice. Reactions were run in triplicate for each animal. Statistical analysis was performed with unpaired t-test using GraphPad Prism 7 software (RRID:SCR_002798). Data represent mean ± SEM.

### Western Blot Analysis

The AuC and FC were removed from each mouse (n=4-5 mice per group), cooled in PBS, and homogenized in ice-cold lysis buffer (50 mM Tris-HCl, pH 7.4, 150 mM NaCl, 5 mM EDTA, 0.05% Triton X-100, and 1 mM PMSF) containing protease inhibitor cocktail (Sigma, catalog # P8340) and phosphatase inhibitor cocktail (Sigma, catalog # P0044). The samples were processed as previously described with modifications^22^. The samples were rotated at 4°C for at least 30 min to allow for complete cell lysis and then cleared by centrifugation at 13,200 rpm for 20 min at 4°C. Supernatants were isolated and boiled in reducing sample buffer (Laemmli 2× concentrate, catalog # S3401, Sigma), and separated on 8–16% Tris-Glycine SDS-PAGE precast gels (catalog # EC6045BOX, Life Technologies). Proteins were transferred onto Protran 0.45 μm Nitrocellulose membrane (GE Healthcare catalog # 10600007, Chicago, Illinois, USA) and blocked for 1 h at room temperature in 5% skim milk (Catalog #170-6404, Bio-Rad). Primary antibody incubations were performed overnight at 4°C with antibodies diluted in TBS/0.1% Tween-20/1% BSA. The following primary antibodies were used: rabbit anti-GABA_A_Rγ2 (1:1000; ThermoFisher Scientific, catalog #14104-1-AP, RRID:AB_10693527); rabbit anti-GABA_A_Rα5 (1:1000; ThermoFisher Scientific, catalog #PA5-31163, RRID:AB_2548637); rabbit anti-GAT3 (1:1000; Millipore Cat# AB1574, RRID:AB_90779); and mouse anti-βactin (1:1000; Santa Cruz Biotechnology Cat# sc-47778, RRID:AB_626632).

Blots were washed 3 × 10 min with TBS/0.1% Tween-20 and incubated with the appropriate HRP-conjugated secondary antibodies for 2 h at room temperature in a TBS/0.1% Tween-20/1% BSA solution. The secondary antibodies used were HRP-conjugated donkey anti-mouse IgG (Jackson ImmunoResearch, catalog #715-035-150, RRID: AB_2340770) or HRP-conjugated goat anti-rabbit IgG (Jackson ImmunoResearch, catalog #111-035-003, RRID: AB_2313567). After secondary antibody incubations, blots were washed 3 × 10 min in TBS/0.1% Tween-20, washed 1 × 10 min in TBS, incubated in Pierce ECL Western Blotting Substrate (Thermo Scientific, catalog #32106) and imaged using ChemiDoc imaging system (Bio-Rad, RRID:SCR_019037). For re-probing, membrane blots were washed in stripping buffer (2% SDS, 100 mM β-mercaptoethanol, 50 mM Tris-HCl, pH 6.8) for 30 min at 55°C, then rinsed repeatedly with TBS/0.1% Tween-20, finally blocked with 5% skim milk, and then re-probed. The first set of blots were probed for GAT3 and β-actin. The second set of blots were probed for GABRG2, GABRA5, and β-actin. Astrocyte cell culture samples were probed for FMRP using rabbit anti-FMRP (1:1000, Sigma, #F4055, RRID:AB_1840858); glutamic acid decarboxylase (GAD) 65-67 using rabbit anti-GAD 65-67 (1:1000; Abcam, catalog #183999); and aldehyde dehydrogenase (ADLH) 1A1 using rabbit anti-ALDH1A1(1:1000; Abcam Cat# ab52492, RRID:AB_867566) with the corresponding secondary HRP-conjugated antibody and re-probed for β-actin (as described above). Band density was analyzed by measuring band and background intensity using Adobe Photoshop CS5.1 software (RRID:SCR_014199). 3-4 samples per group (Ctrl WT vs. cKO, or WT vs KO for mouse astrocyte cultures, or CON vs FXS for human astrocyte cultures) were run per blot, and precision/tolerance (P/T) ratios for individual cKO samples were normalized to P/T ratios of Ctrl WT samples with the similar actin levels. Statistical analysis was performed with unpaired t-test using GraphPad Prism 7 software (RRID: SCR_002798). Data represent mean ± SEM.

### Surgery for in vivo EEG recordings

P28 male Ctrl WT (n = 14), cKO (n = 13), P22-P28 WT (n=43) and global *Fmr1* KO (n=33) mice were used for the EEG studies as previously described with modifications^22,67^. Four-five days before EEG recording, mice were anesthetized with isoflurane inhalation (0.2-0.5%) and an injection of ketamine, xylazine, and acepromazine (K/X/A) (i.p. 80/10/1 mg/kg), and then secured in a bite bar, and placed in a stereotaxic apparatus (model 930; Kopf, CA). Artificial tear gel was applied to the eyes to prevent drying. Toe pinch reflex was used to measure anesthetic state every 10 min throughout the surgery, and supplemental doses of ¼ original dose ketamine only (i.p. 80 mg/kg) were administered as needed. Once the mouse was anesthetized, a midline sagittal incision was made along the scalp to expose the skull. A Foredom dental drill was used to drill 1 mm diameter holes in the skull overlying the right auditory cortex (−1.6 mm, +4.8 mm), left frontal lobe (+3.0 mm, −1.6 mm), and left occipital (−4.2 mm, −5.1 mm) (coordinate relative to Bregma: anterior/posterior, medial/lateral). Three channel electrode posts from Protech International (8IMS3332AXXE-5MM MS333/2-A ELECT SS 3C UNTW) were attached to 1.6 mm stainless steel screws from Protech International (8L0X3905201F) and screws were advanced into drilled holes until secure. Special care was taken not to advance the screws beyond the point of contact with the Dura. Dental cement was applied around the screws, on the base of the post, and exposed skull. Triple antibiotic was applied along the edges of the dental cement followed by an injection of extended-release Buprenorphine (Ethiqa XR; s.c. 3.25 mg/kg). Mice were placed on a heating pad to aid recovery from anesthesia. Mice were monitored daily until the day of EEG recordings. The separation between the last post-surgical Buprenorphine injection and EEG recordings was between 4 and 5 days.

### Electrophysiology

EEG recordings were first performed 4-5 days after surgery. Second recording was done 1 or 2 days later after 4 h post injection with SNAP-5114 (i.p. 25 mg/kg) or vehicle. Baseline and auditory event-related potential (ERP) recordings were obtained using the BioPac system (BIOPAC Systems, Inc.) from awake and freely moving mice as published previously with modifications^67^. Mice were allowed to habituate to the recording chamber for 15 min prior to being connected to the BioPac system. A three-channel tether was connected to the electrode post (implanted during surgery) under brief isoflurane anesthesia. The mouse was then placed inside a grounded Faraday cage after recovery from isoflurane. This tether was then connected to a commutator located directly above the cage. Mice were then allowed to habituate while being connected to the tether for an additional 20 min before EEG recordings were obtained.

The BioPac MP150 acquisition system was connected to two EEG 100C amplifier units (one for each channel) to which the commutator was attached. The lead to the occipital cortex was used as reference for both frontal and auditory cortex screw electrodes. The acquisition hardware was set to high-pass (>0.5 Hz) and low-pass (<100 Hz) filters. Normal EEG output data were collected with gain maintained the same (10,000x) between all recordings. Data were sampled at a rate of either 2.5 or 5 kHz using Acqknowledge software and down sampled to 1024 Hz post hoc using Analyzer 2.1 (Brain Vision Inc.).

Sound delivery was synchronized with EEG recording using a TTL pulse to mark the onset of each sound in a train. Baseline EEGs were recorded for 5 min (no auditory stimuli were presented), followed by recordings in response to auditory stimulation.

### Acoustic Stimulation

All experiments were conducted in a sound-attenuated chamber lined with anechoic foam (Gretch-Ken Industries, OR) as previously described with modifications^22,67^. Acoustic stimuli were generated using RVPDX software and RZ6 hardware (Tucker-Davis Technologies, FL) and presented through a free-field speaker (MF1 Multi-Field Magnetic Speaker; Tucker-Davis Technologies, FL) located 12 inches directly above the cage. Sound pressure level (SPL) was modified using programmable attenuators in the RZ6 system. The speaker output was ∼65-70 dB SPL at the floor of the recording chamber with fluctuation of +/-3 dB for frequencies between 5 and 35 kHz as measured with a ¼ inch Bruel & Kjaer microphone.

We used acoustic stimulation paradigms that have been previously established in *Fmr1* KO mice^67^, which is analogous to work in humans with FXS^87^. A chirp-modulated signal (henceforth, ‘chirp’) to induce synchronized oscillations in EEG recordings was used to measure temporal response fidelity. The chirp is a 2 s broadband noise stimulus with amplitude modulated (100% modulation depth) by a sinusoid whose frequencies increase (Up-chirp) or decrease (Down-chirp) linearly in the 1-100 Hz range^102–104^. The chirp facilitates a rapid measurement of transient oscillatory response (delta to gamma frequency range) to auditory stimuli of varying frequencies and can be used to compare oscillatory responses in different groups in clinical and pre-clinical settings^104^. Inter-trial coherence analysis^105^ can then be used to determine the ability of the neural generator to synchronize oscillations to the frequencies present in the stimulus.

To avoid onset responses contaminating phase locking to the amplitude modulation of the chirp, the stimulus was ramped in sound level from 0-100% over 1 s (rise time), which then smoothly transitioned into chirp modulation of the noise. Up and Down chirp trains were presented 300 times each (for a total of 600 trains). Both directions of modulation were tested to ensure any frequency specific effects were not due to the frequency transition history within the stimulus. Up- and Down-chirp trains were presented in an alternating sequence. The interval between each train was randomly varied between 1 and 1.5 s. To study evoked response amplitudes and habituation, trains of 100 ms broadband noise were presented at two repetition rates, 0.25 Hz (a non-habituating rate) and 4 Hz (a strongly habituating rate)^75^. Each train consisted of 10 noise bursts and the inter-train interval used was 8 seconds. Each repetition rate was presented 100 times in an alternating pattern^75^. The onset of trains and individual noise bursts were tracked with separate TTL pulses that were used to quantify latency of response.

### EEG Data Analysis

Data were extracted from Acqknowledge and files saved in a file format (EDF) compatible with BrainVision Analyzer 2.1 software as previously described with modifications^22,67^. All data were notch filtered at 60 Hz to remove residual line frequency power from recordings. EEG artifacts were removed using a semi-automatic procedure in Analyzer 2.1 for all recordings. Less than 30% of data were rejected due to artifacts from any single mouse.

Baseline EEG data were divided into 2 s segments and Fast Fourier Transforms (FFT) were calculated on each segment using 0.5 Hz bins and then average power (µV2/Hz) was calculated for each mouse from 1-100 Hz. Power was then binned into standard frequency bands: Delta (1-4 Hz), Theta (4-10 Hz), Alpha (10-13 Hz), Beta (13-30 Hz), Low Gamma (30-55 Hz), and High Gamma (65-100 Hz). Responses to auditory stimuli were analyzed using Morlet wavelet analysis. Chirp trains were segmented into windows of 500 ms before chirp onset to 500 ms after the end of the chirp sound (total of 3 s because each chirp was 2 s in duration). EEG traces were processed with Morlet wavelets from 1-100 Hz using complex number output (voltage density, µV/Hz) for ITPC calculations, and power density (µV2/Hz) for non-phase locked single trial power (STP) calculations. Wavelets were run with a Morlet parameter of 10 as this gave the best frequency/power discrimination. This parameter was chosen since studies in humans found most robust difference around 40 Hz, where this parameter is centered^87^. To measure phase synchronization at each frequency across trials Inter Trial Phase Coherence (ITPC) was calculated. The equation used to calculate ITPC is: 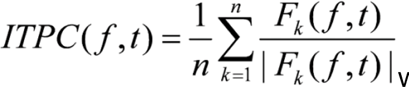 where f is the frequency, t is the time point, and k is trial number. Thus, Fk(f,t) refers to the complex wavelet coefficient at a given frequency and time for the kth trial. There were no less than 270 trials (out of 300) for any given mouse after segments containing artifacts were rejected. All raw EEG data analysis is available on: https://drive.google.com/drive/folders/1Qo0FitDb6EO6lJ6euq3x3jGo8Yw7wGdC?usp=sha ring

### EEG Statistical Analysis

Statistical group comparisons of chirp responses (ITPC and STP) were quantified by wavelet analysis. Analysis was conducted by binning time into 256 parts and frequency into 100 parts, resulting in a 100×256 matrix. Non-parametric cluster analysis was used to determine contiguous regions in the matrix that were significantly different from a distribution of 1000 randomized Monte Carlo permutations based on previously published methods with modifications^106^. Briefly, if the cluster sizes of the real group assignments (both positive and negative direction, resulting in a two-tailed alpha of p = 0.025) were larger than 97.25% of the random group assignments, those clusters were considered significantly different between groups. This method avoids statistical assumptions about the data and corrects for multiple comparisons.

### Behavioral Assessments

Behavioral tests were conducted in P28 male Ctrl WT (n=30), female Het (n=36), and male and female cKO (n=43) mice, and male cKO SNAP mice (n=10) treated with SNAP-5114 (25 mg/mL) ip injection 4 hours prior to the start of testing. Mice were first subjected to open field test, followed by elevated plus maze.

#### Open-field test

Locomotor activity and exploratory behaviors were tested in P28 Ctrl WT, Het, and cKO mice as described previously with modifications^22^. A 72 × 72-cm open-field arena with 50- cm-high walls was constructed from opaque acrylic sheets with a clear acrylic sheet for the bottom. The open field arena was placed in a brightly lit room, and one mouse at a time was placed in a corner of the open field and allowed to explore for 10 min while being recorded with digital video from above. The floor was cleaned with 2–3% acetic acid, 70% ethanol, and water between tests to eliminate odor trails. The mice were tested between the hours of 9:00 A.M. and 2:00 P.M., and this test was always performed prior to the elevated plus maze. The arena was subdivided into a 4 × 4 grid of squares with the middle of the grid defined as the center. A line 4 cm from each wall was added to measure thigmotaxis. Locomotor activity was scored by the analysis of total line crosses and speed using TopScan Lite software (Clever Sys., Inc., VA). Time in open field compared to thigmotaxis was used as an indicator of exploratory behavior. The analysis was performed in 5 min intervals for the total 10 min exploration duration. Assessments of the digital recordings were performed blind to the condition. Statistical analysis was performed with unpaired t-test using GraphPad Prism 7 software (RRID: SCR_002798). Data represent mean ± SEM.

#### Elevated plus maze

Anxiety-like behavior was tested in P28 Ctrl WT, Het, and cKO mice as described previously with modifications^22^.The elevated plus maze consisted of four arms in a plus configuration. Two opposing arms had 15-cm tall walls (closed arms), and two arms were without walls (open arms). The entire maze sat on a stand 1 m above the floor. Each arm measured 30 cm long and 10 cm wide. Mice were allowed to explore the maze for 10 min while being recorded by digital video from above. The maze was wiped with 2–3% acetic acid, 70% ethanol and water between each test to eliminate odor trails. The mice were tested between the hours of 9:00 A.M. and 2:00 P.M. This test was always done following the open-field test. TopScan Lite software was used to measure the percent of time spent in open arms and speed. The time spent in open arm was used to evaluate anxiety-like behavior while speed and total arm entries were measured to evaluate overall locomotor activity^22^. The analysis was performed in 5 min intervals for the total 10 min exploration duration. Assessments of the digital recordings were done blind to the condition using TopScan Lite software. Statistical analysis was performed with unpaired t-test using GraphPad Prism 7 software. Data represent mean ± standard error of the mean (SEM).

## Acknowledgements

This work was supported by the National Institute of Neurological Disorders and Stroke (NS129555 to I.M.E.), FRAXA research grant (to M.L.C), FRAXA Research Foundation fellowship (to A.O,N.), TRANSCEND fellowship from the California Institute of Regenerative Medicine (Award # EDUC4-12752 to V.A.W.), the National Institute of Neurological Disorders and Stroke National Research Service Award Fellowship (1F31NS117178-01 to M.R), National Institutes of Health MARC U STAR undergraduate research program (T34GM062756 to J.W.), and grant from Jane and Aatos Erkko Foundation (to M.L.C). The contents of this publication are solely the responsibility of the authors and do not necessarily represent official views of CIRM or other agencies of the State of California. We are grateful to Dr. Gary Bassell and his lab at Emory University and Dr. Karen Usdin and her lab at NIH/NIDDK for supplying iPSC lines used in this publication. We thank Ken Furuichi for technical support with mouse colony; Patricia Pirbhoy for help with EEG analysis; members of the Ethell, Binder, Razak, and Goel laboratories for helpful discussions; and Dr. David Carter for advice on confocal microscopy.

