## Supplementary data for "Astrocytic GABA transport controls fidelity of temporal processing"

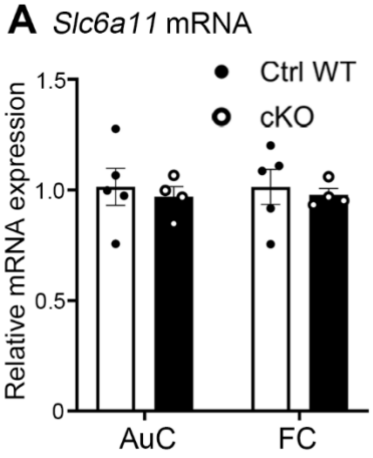

**Figure S1: Astrocyte-specific Fmr1 deletion does not affect expression of mRNA encoding GAT3, related to Figure 3**

A, Quantitative analysis of *Slc6a11* mRNA performed with qPCR. Graph shows mean ± SEM (n=4-5 mice/group; p=0.05; t-test). We found no significant differences in *Slc6a11* expression in cKO mice compared to Ctrl WT.

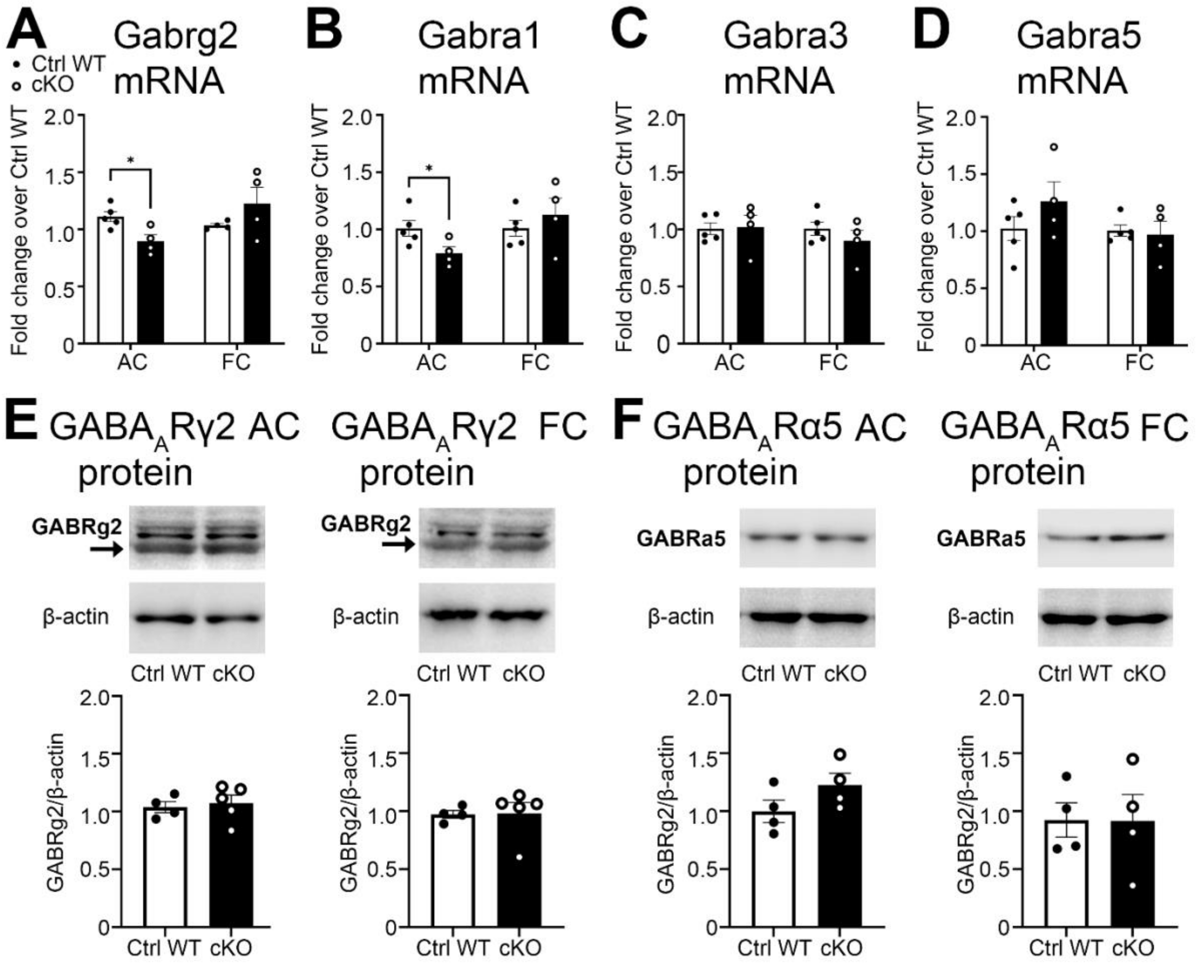

**Figure S2: Effects of astrocyte-specific Fmr1 deletion on mRNA expression and protein levels of GABAA receptor subunits, related to Figure 4**

A-D, Quantitative analysis of mRNA encoding $GABA_A$ receptor subunits performed with qPCR. Graphs show mean ± SEM (n=4-5 mice/group; *p<0.05; **p<0.01; t-test). There is a significant increase in mRNA levels of *Gabrg2* (A) and *Gabra1* (B) in the frontal cortex of cKO mice compared to Ctrl WT. No significant differences were observed in mRNA levels of *Gabra3* (C) and *Gabra5* (D). E-F, Western blots showing GABAARγ2 (E, arrows indicate the measured band), GABAARα5 (F), and beta-actin protein levels in lysates from auditory cortex (left) and frontal cortex (right) of Ctrl WT and cKO mice. Graphs show mean ± SEM (n=3-4/group, p=0.05; t-test). No differences were observed in GABAARγ2 or GABAARα5 protein levels.

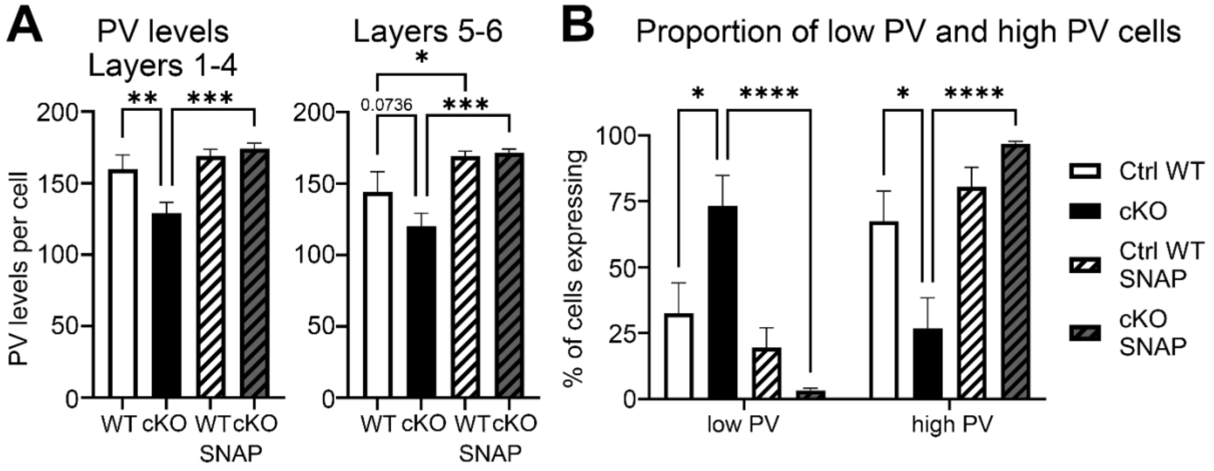

**Figure S3: Parvalbumin levels are significantly reduced in superficial layers in AuC of astrocyte-specific cKO mice and enhanced following acute blockage of GAT3-mediated astrocytic GABA transport, related to Figure 5**

A, Quantitative analysis of average PV immunoreactivity per cell in L1-4 (left) and L5/6 (right) AuC of Ctrl WT, cKO, Ctrl WT SNAP, and cKO SNAP groups. Graph shows mean ± SEM (n= 3-5 mice/group, n = 4 images/mouse, *$p<0.05$; **$p < 0.01$; ***$p < 0.001$; two-way ANOVA, Fisher's LSD post hoc test). PV levels are significantly reduced in L1-4 of cKO mice and are trending towards significance in L5/6. SNAP treatment significantly increases PV in L1-4 and L5/6 of cKO and L5/6 of Ctrl WT. B, Proportion of low expressing PV cells and high expressing PV cells in L1-6 AuC of Ctrl WT, cKO, Ctrl WT SNAP, and cKO SNAP groups. Graph shows mean ± SEM (n= 3-5 mice/group, n = 4 images/mouse, *$p<0.05$; ****$p < 0.0001$; two-way ANOVA, Šídák's post hoc test). cKO mice have a significantly higher proportion of low PV cells than Ctrl WT. SNAP treatment significantly increased proportion of high PV cells in cKO.

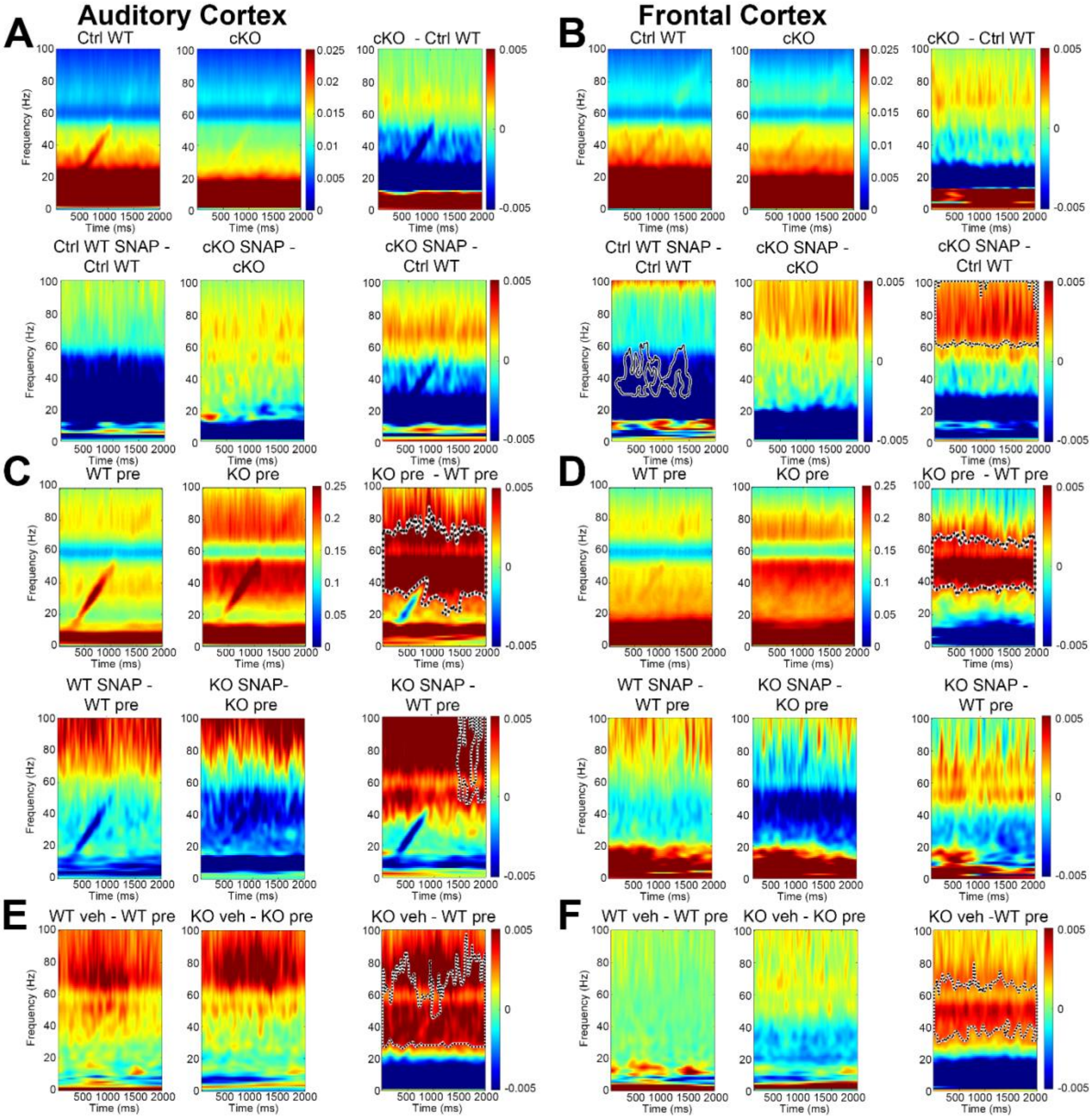

**Figure S4: Acute blockage of GAT3-mediated GABA transport in astrocytes normalized background gamma power in the auditory and frontal cortex of global KO mice during chirp presentation, related to Figure 7**

For each mouse, single-trial power (STP) was measured to determine the average total non-phase locked power during chirp train presentation. A-B, Grand average matrices were calculated for each genotype, and then Ctrl WT (n=12) STP values were subtracted from cKO (n=13) values and pre-treatment STP values were subtracted from SNAP-treated values (bottom panels) for AuC (A) and FC (B). Statistical cluster analysis reveals contiguous time x frequency regions that are significantly different between genotypes. Black dashed contour indicates these significant clusters. Consistent with the increase in gamma power changes in baseline EEGs, SNAP-treated cKO mice express statistically significant increase in non-phase gamma power range (30-100 Hz, red) throughout sound presentation in FC. C-D, Grand average of STP values were calculated for global KO (n=13) and WT (n=21) before (pre) and global KO (n=7) and WT (n=11) after SNAP treatment and subtracted to generate difference plots by genotype (right panels) and treatment (bottom panels) for AuC (C) and FC (D). Global KO mice express statistically significant increase in STP at gamma frequencies (30-100 Hz, blue), with gamma STP restored to Ctrl WT levels in AuC and FC following SNAP treatment. E-F, Grand average matrices of STP values were calculated for global KO (n=20) and WT (n=22) before (pre) and global KO (n=12) and WT (n=16) after vehicle treatment and subtracted to generate difference plots by genotype (right panels) and treatment (bottom panels) for AuC (E) and FC (F). Vehicle treatment did not significantly affect STP values in gamma range.

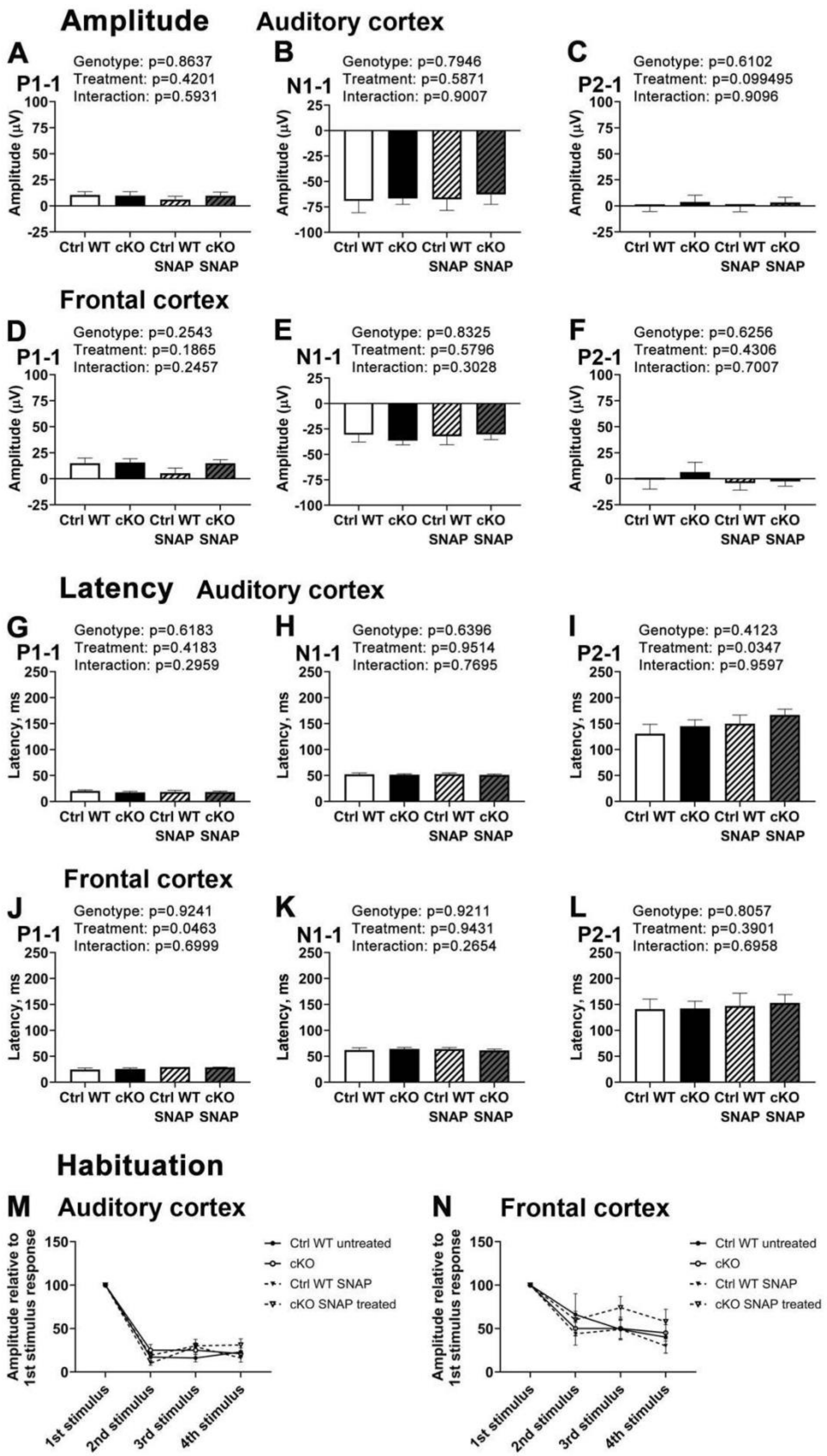

Amplitude
Auditory cortex
A
P1-1
Genotype: p=0.8637
Treatment: p=0.4201
Interaction: p=0.5931
B
N1-1
Genotype: p=0.7946
Treatment: p=0.5871
Interaction: p=0.9007
C
P2-1
Genotype: p=0.6102
Treatment: p=0.099495
Interaction: p=0.9096
Amplitude (µV)
Ctrl WT
cKO
Ctrl WT SNAP
cKO SNAP
Frontal cortex
D
P1-1
Genotype: p=0.2543
Treatment: p=0.1865
Interaction: p=0.2457
E
N1-1
Genotype: p=0.8325
Treatment: p=0.5796
Interaction: p=0.3028
F
P2-1
Genotype: p=0.6256
Treatment: p=0.4306
Interaction: p=0.7007
Latency
Auditory cortex
G
P1-1
Genotype: p=0.6183
Treatment: p=0.4183
Interaction: p=0.2959
H
N1-1
Genotype: p=0.6396
Treatment: p=0.9514
Interaction: p=0.7695
I
P2-1
Genotype: p=0.4123
Treatment: p=0.0347
Interaction: p=0.9597
Latency, ms
Frontal cortex
J
P1-1
Genotype: p=0.9241
Treatment: p=0.0463
Interaction: p=0.6999
K
N1-1
Genotype: p=0.9211
Treatment: p=0.9431
Interaction: p=0.2654
L
P2-1
Genotype: p=0.8057
Treatment: p=0.3901
Interaction: p=0.6958
Habituation
M
Auditory cortex
N
Frontal cortex
Amplitude relative to 1st stimulus response
1st stimulus
2nd stimulus
3rd stimulus
4th stimulus
Ctrl WT untreated
cKO
Ctrl WT SNAP
cKO SNAP treated

**Figure S6: Astrocyte-specific Fmr1 deletion does not affect habituation, latency or amplitude of ERP waves, related to Figure 7**

A-L, Auditory ERP amplitudes (A-F) and latencies (G-L) in AuC and FC of Ctrl WT (n=9), cKO (n=13), Ctrl WT SNAP (n=7), cKO SNAP (n=12) mice. Grand average ERPs obtained from P28 mice in response to the first 100-ms broadband noise presented at 4Hz repetition rate. P1, N1, and P2 were defined as maximum or minimum voltage deflections within 0–30 ms, 30–80 ms, or 80–150 ms, respectively. A-L, Graphs show mean ± SEM of P1 amplitude in the AuC (A) and FC (D); N1 amplitude in the AuC (B) and FC (E); P2 amplitude in the AuC (C) and FC (F); P1 latency in the AuC (G) and FC (J); N1 latency in the AuC (H) and FC (K); P2 latency in the AuC (I) and FC (L) ($p=0.05$; Mixed-effects analysis, Fisher's LSD post hoc test). No significant differences were observed with post-hoc testing; however, there is an effect of treatment on latency in P1 in FC and P2 in AuC. M-N, Habituation is indicated by N1 amplitude of subsequent stimuli relative to first stimulus amplitude in 4Hz sound train in AuC (M) and FC (N). Graphs show mean ± SEM. No differences are observed in habituation.

**Table S1**

Statistics for Figure 1B

| % GFAP/S100β+ cells | **Mean** | **SEM** | **N** |
|---|---|---|---|
| CON | 96.58 | 0.3212 | 3 |
| FXS | 97.79 | 0.4840 | 3 |
| Statistics | Two-tailed, unpaired t-test | t=2.089, df=4, p=0.1050 $\eta^2$= 0.5217 | |

List of primer sequences used for qRT-PCR in Figures 1C, 1E-1G, S1, S2A-S2D

| **Gene** | **Forward primer sequence (5'–3')** | **Reverse primer sequence (5'–3')** |
|---|---|---|
| **Human** | | |
| *Fmr1* | GGG GAA TCC CAG AAA CCT GAA | CGC AAC TGG TCT ACT TCC TTT A |
| *GFAP* | ACC TGC AGA TTC GAG AAA CCA G | GGT CCT GCC TCA CAT CAC ATC |
| *S100B* | GTG GCC CTC ATC GAC GTT TT | ACC TCC TGC TCT TTG ATT TCC TCT |
| *Slc1a3* | ATT CCA GCA GGG AGT CCG TA | TCC AAG GAT TGT ACC CAC AAT GA |
| *GAPDH* | TTG GCT ACA GCA ACA GGG TG | GGG GAG ATT CAG TGT GGT GG |
| **Mouse** | | |
| *Slc6a11* | CGG CTG GGT ATA TGG AAG CA | ACG ACT TTC CAG CAC CAC TT |
| *Gabra1* | CAGAAAAGCCAAAGAAAGTAAAGGA | TGGTTGCTGTAGGAGCATATGTG |
| *Gabra3* | GCTGCTCAGACTGGTAGATAATGG | GGGCATTCAGCGTGTATTGTT |
| *Gabrg2* | CCTGCCCCCTGGAGTTCT | ACTGCGCTTCCATTGATAAACA |
| *Gabra5* | GCAGACAGTAGGCACTGAGAACA | GGAAGTGAGCAGTCATGATCGTAT |
| *GAPDH* | ACTCCACTCACGGCAAATTC | TCTCCATGGTGGTGAAGACA |

Statistics for Figure 1C

| FMRP mRNA | **Mean** | **SEM** | **N** |
|---|---|---|---|
| CON | 0.9638 | 0.2759 | 5 |
| FXS | 0.02298 | 0.02037 | 5 |
| Statistics | Two-tailed, unpaired t-test | t=3.400, df=8, p=0.0094 $\eta^2$= 0.5910 | |

Statistics for Figure 1D

| FMRP protein | **Mean** | **SEM** | **N** |
|---|---|---|---|
| CON | 1.000 | 0.4714 | 3 |
| FXS | 0.05250 | 0.01550 | 2 |
| Statistics | Two-tailed, unpaired t-test | t=1.557, df=3, p=0.2173 $\eta^2$= 0.4469 | |

Statistics for Figure 1E

| GFAP mRNA | **Mean** | **SEM** | **N** |
|---|---|---|---|
| CON | 0.8353 | 0.1844 | 6 |
| FXS | 0.4433 | 0.2277 | 5 |
| Statistics | Two-tailed, unpaired t-test | t=1.354, df=9, p=0.2088<br>$\eta^2$= 0.1692 | |

Statistics for Figure 1F

| S100β mRNA | **Mean** | **SEM** | **N** |
|---|---|---|---|
| CON | 0.9639 | 0.1307 | 6 |
| FXS | 0.5080 | 0.1980 | 5 |
| Statistics | Two-tailed, unpaired t-test | t=1.984, df=9, p=0.0786<br>$\eta^2$= 0.3043 | |

Statistics for Figure 1G

| SLC1A3 mRNA | **Mean** | **SEM** | **N** |
|---|---|---|---|
| CON | 0.7899 | 0.2261 | 6 |
| FXS | 0.2821 | 0.1047 | 5 |
| Statistics | Two-tailed, unpaired t-test | t=1.900, df=9, p=0.0899<br>$\eta^2$= 0.2863 | |

Statistics for Figure 1H

| GABA HPLC | **Mean** | **SEM** | **N** |
|---|---|---|---|
| CON | 1.277 | 0.8225 | 3 |
| FXS | 8.768 | 1.734 | 4 |
| Statistics | Two-tailed, unpaired t-test | t=3.461, df=5, p=0.0180<br>$\eta^2$= 0.7055 | |

Statistics for Figure 1I

| GAD65/67 protein | **Mean** | **SEM** | **N** |
|---|---|---|---|
| CON | 1.000 | 0.05619 | 6 |
| FXS | 1.250 | 0.07430 | 7 |
| Statistics | Two-tailed, unpaired t-test | t=2.605, df=11, p=0.0245,<br>$\eta^2$= 0.3816 | |

**Table S2**

Statistics for Figure 2A

| GABA levels | **Mean** | **SEM** | **N** |
|---|---|---|---|
| WT | 1.000 | 0.04265 | 5 |
| KO | 1.248 | 0.07530 | 5 |
| Statistics | Two-tailed, unpaired t-test | t=2.864, df=8, p=0.0210<br>$\eta^2$=0.5062 | |

Statistics for Figure 2B

| GAD65/67 protein | **Mean** | **SEM** | **N** |
|---|---|---|---|
| WT | 1.000 | 0.01723 | 5 |
| KO | 1.525 | 0.1727 | 6 |
| Statistics | Two-tailed, unpaired t-test | t=2.742, df=9, p=0.0228 $\eta^2$=0.4551 | |

| Aldha1 protein | **Mean** | **SEM** | **N** |
|---|---|---|---|
| WT | 1.000 | 0.1396 | 5 |
| KO | 0.8600 | 0.05125 | 6 |
| Statistics | Two-tailed, unpaired t-test | t=1.013, df=9, p=0.3373 $\eta^2$=0.1024 | |

**Table S3**

Statistics for Figure 3C

| FMRP in astrocytes | **Mean** | **SEM** | **N** |
|---|---|---|---|
| Ctrl WT | 14.3 | 0.96 | 3 |
| cKO | 1.76 | 0.11 | 3 |
| Statistics | Two-tailed, unpaired t-test | t=12.93, df=4, p=0.0002 $\eta^2$=0.9768 | |

Statistics for Figure 3E

GABA levels in astrocytes in AuC

| **ANOVA table** | **SS (Type III)** | **DF** | **MS** | **F (DFn, DFd)** | **P value** | **% of total variation** |
|---|---|---|---|---|---|---|
| Genotype x Treatment | 14215 | 1 | 14215 | F (1, 118) = 19.23 | P<0.0001 | 11.10 |
| Genotype | 13383 | 1 | 13383 | F (1, 118) = 18.10 | P<0.0001 | 10.45 |
| Treatment | 15641 | 1 | 15641 | F (1, 118) = 21.16 | P<0.0001 | 12.21 |

| **Uncorrected Fisher's LSD** | **Predicted (LS) mean diff.** | **95.00% CI of diff.** | **Individual P Value** |
|---|---|---|---|
| Ctrl WT | | | |
| Vehicle vs. SNAP | -1.090 | -13.55 to 11.37 | 0.8627 |
| cKO | | | |
| Vehicle vs. SNAP | -45.65 | -61.44 to -29.85 | <0.0001 |
| Vehicle | | | |
| Ctrl WT vs. cKO | 0.6621 | -14.20 to 15.52 | 0.9299 |
| SNAP | | | |
| Ctrl WT vs. cKO | -43.89 | -57.46 to -30.33 | <0.0001 |

Statistics for Figure 3F
GABA levels in astrocytes in FC

| ANOVA table | SS (Type III) | DF | MS | F (DFn, DFd) | P value | % of total variation |
|---|---|---|---|---|---|---|
| Genotype x Treatment | 3309 | 1 | 3309 | F (1, 85) = 6.717 | P=0.0112 | 4.540 |
| Genotype | 27351 | 1 | 27351 | F (1, 85) = 55.52 | P<0.0001 | 37.53 |
| Treatment | 1844 | 1 | 1844 | F (1, 85) = 3.743 | P=0.0563 | 2.530 |

| Uncorrected Fisher's LSD | Predicted (LS) mean diff. | 95.00% CI of diff. | Individual P Value |
|---|---|---|---|
| Ctrl WT | | | |
| Vehicle vs. SNAP | 3.106 | -9.633 to 15.84 | 0.6291 |
| cKO | | | |
| Vehicle vs. SNAP | -21.40 | -35.22 to -7.575 | 0.0028 |
| Vehicle | | | |
| Ctrl WT vs. cKO | -22.97 | -36.00 to -9.944 | 0.0007 |
| SNAP | | | |
| Ctrl WT vs. cKO | -47.47 | -61.02 to -33.92 | <0.0001 |

Statistics for Figure 3G

| GAT3 protein AC | Mean | SEM | N |
|---|---|---|---|
| Ctrl WT | 1.000 | 0.1560 | 5 |
| cKO | 0.9848 | 0.1299 | 5 |
| Statistics | Two-tailed, unpaired t-test | t=0.07491, df=8, p=0.9421 $\eta^2$=0.0007010 | |

| GAT3 protein FC | Mean | SEM | N |
|---|---|---|---|
| Ctrl WT | 1.000 | 0.03907 | 5 |
| cKO | 1.052 | 0.1186 | 4 |
| Statistics | Two-tailed, unpaired t-test | t=0.4597, df=7, p=0.6597 $\eta^2$=0.02930 | |

**Table S4**
Statistics for Figure 4C
vGAT/Gephyrin co-localization

| ANOVA table | SS (Type III) | DF | MS | F (DFn, DFd) | P value | % of total variation |
|---|---|---|---|---|---|---|
| Brain region x Genotype | 9.698 | 1 | 9.698 | F (1, 129) = 0.02451 | P=0.8758 | 0.005079 |
| Brain region | 134845 | 1 | 134845 | F (1, 129) = 340.8 | P<0.0001 | 70.62 |
| Genotype | 7143 | 1 | 7143 | F (1, 129) = 18.05 | P<0.0001 | 3.741 |

| Uncorrected Fisher's LSD | Predicted (LS) mean diff. | 95.00% CI of diff. | Individual P Value |
|---|---|---|---|
| AuC | | | |
| Ctrl WT vs. cKO | 15.21 | 5.663 to 24.76 | 0.0020 |
| FC | | | |
| Ctrl WT vs. cKO | 14.13 | 4.358 to 23.90 | 0.0049 |
| Ctrl WT | | | |
| AC vs. FC | -63.21 | -72.91 to -53.50 | <0.0001 |
| cKO | | | |
| AC vs. FC | -64.29 | -73.90 to -54.67 | <0.0001 |

Statistics for Figure 4D
vGAT puncta density

| ANOVA table | SS (Type III) | DF | MS | F (DFn, DFd) | P value | % of total variation |
|---|---|---|---|---|---|---|
| Brain region x Genotype | 28.84 | 1 | 28.84 | F (1, 129) = 0.04784 | P=0.8272 | 0.009404 |
| Brain region | 224452 | 1 | 224452 | F (1, 129) = 372.3 | P<0.0001 | 73.19 |
| Genotype | 8242 | 1 | 8242 | F (1, 129) = 13.67 | P=0.0003 | 2.688 |

| Uncorrected Fisher's LSD | Predicted (LS) mean diff. | 95.00% CI of diff. | Individual P Value |
|---|---|---|---|
| AuC | | | |
| Ctrl WT vs. cKO | 14.83 | 2.854 to 26.82 | 0.0156 |
| FC | | | |
| Ctrl WT vs. cKO | 16.70 | 4.818 to 28.58 | 0.0062 |
| Ctrl WT | | | |
| AC vs. FC | -83.21 | -95.09 to -71.33 | <0.0001 |
| cKO | | | |
| AC vs. FC | -81.35 | -93.33 to -69.37 | <0.0001 |

Statistics for Figure 4E
PV/vGAT co-localization

| ANOVA table | SS (Type III) | DF | MS | F (DFn, DFd) | P value | % of total variation |
|---|---|---|---|---|---|---|
| Brain region x Genotype | 133.9 | 1 | 133.9 | F (1, 64) = 1.319 | P=0.2550 | 0.7827 |
| Brain region | 10064 | 1 | 10064 | F (1, 64) = 99.14 | P<0.0001 | 58.83 |
| Genotype | 452.7 | 1 | 452.7 | F (1, 64) = 4.459 | P=0.0386 | 2.646 |

| Uncorrected Fisher's LSD | Predicted (LS) mean diff. | 95.00% CI of diff. | Individual P Value |
|---|---|---|---|
| AuC | | | |
| Ctrl WT vs. cKO | 2.356 | -4.560 to 9.271 | 0.4987 |
| FC | | | |
| Ctrl WT vs. cKO | 7.974 | 1.070 to 14.88 | 0.0243 |
| Ctrl WT | | | |
| AC vs. FC | -27.16 | -33.97 to -20.35 | <0.0001 |
| cKO | | | |
| AC vs. FC | -21.54 | -28.55 to -14.53 | <0.0001 |

Statistics for Figure 4F
PV puncta density

| ANOVA table | SS (Type III) | DF | MS | F (DFn, DFd) | P value | % of total variation |
|---|---|---|---|---|---|---|
| Brain region x Genotype | 259.3 | 1 | 259.3 | F (1, 63) = 2.469 | P=0.1211 | 1.504 |
| Brain region | 9745 | 1 | 9745 | F (1, 63) = 92.81 | P<0.0001 | 56.51 |
| Genotype | 537.8 | 1 | 537.8 | F (1, 63) = 5.122 | P=0.0271 | 3.119 |

| Uncorrected Fisher's LSD | Predicted (LS) mean diff. | 95.00% CI of diff. | Individual P Value |
|---|---|---|---|
| AuC | | | |
| Ctrl WT vs. cKO | 1.734 | -5.301 to 8.770 | 0.6240 |
| FC | | | |
| Ctrl WT vs. cKO | 9.612 | 2.480 to 16.74 | 0.0091 |
| Ctrl WT | | | |
| AC vs. FC | -28.09 | -35.01 to -21.16 | <0.0001 |
| cKO | | | |
| AC vs. FC | -20.21 | -27.45 to -12.97 | <0.0001 |

**Table S5**
Statistics for Figure 5C
PV levels

| ANOVA table | SS (Type III) | DF | MS | F (DFn, DFd) | P value | % of total variation |
|---|---|---|---|---|---|---|
| Genotype x Treatment | 9575 | 1 | 9575 | F (1, 52) = 5.794 | P=0.0197 | 7.451 |
| Genotype | 5878 | 1 | 5878 | F (1, 52) = 3.557 | P=0.0649 | 4.574 |
| Treatment | 31319 | 1 | 31319 | F (1, 52) = 18.95 | P<0.0001 | 24.37 |

| Tukey's multiple comparisons test | Predicted (LS) mean diff. | 95.00% CI of diff. | Adjusted P Value |
|---|---|---|---|
| Ctrl WT :Vehicle vs. Ctrl WT :SNAP | -21.36 | -59.51 to 16.78 | 0.4528 |
| Ctrl WT :Vehicle vs. cKO:Vehicle | 47.13 | 5.924 to 88.33 | 0.0190 |
| Ctrl WT :Vehicle vs. cKO:SNAP | -27.09 | -68.29 to 14.12 | 0.3115 |
| Ctrl WT :SNAP vs. cKO:Vehicle | 68.49 | 27.29 to 109.7 | 0.0003 |
| Ctrl WT :SNAP vs. cKO:SNAP | -5.721 | -46.92 to 35.48 | 0.9827 |
| cKO:Vehicle vs. cKO:SNAP | -74.21 | -118.3 to -30.16 | 0.0002 |

Statistics for Figure 5D
PV cell density

| ANOVA table | SS (Type III) | DF | MS | F (DFn, DFd) | P value | % of total variation |
|---|---|---|---|---|---|---|
| Genotype x Treatment | 0.1490 | 1 | 0.1490 | F (1, 52) = 0.7172 | P=0.4009 | 1.218 |
| Genotype | 0.9763 | 1 | 0.9763 | F (1, 52) = 4.700 | P=0.0348 | 7.983 |
| Treatment | 0.2392 | 1 | 0.2392 | F (1, 52) = 1.152 | P=0.2882 | 1.956 |

| Tukey's multiple comparisons test | Predicted (LS) mean diff. | 95.00% CI of diff. | Adjusted P Value |
|---|---|---|---|
| Ctrl WT :Vehicle vs. Ctrl WT :SNAP | -0.2363 | -0.6640 to 0.1914 | 0.4648 |
| Ctrl WT :Vehicle vs. cKO:Vehicle | -0.3710 | -0.8330 to 0.09091 | 0.1566 |
| Ctrl WT :Vehicle vs. cKO:SNAP | -0.3989 | -0.8608 to 0.06307 | 0.1131 |
| Ctrl WT :SNAP vs. cKO:Vehicle | -0.1347 | -0.5967 to 0.3272 | 0.8658 |
| Ctrl WT :SNAP vs. cKO:SNAP | -0.1626 | -0.6245 to 0.2994 | 0.7867 |
| cKO:Vehicle vs. cKO:SNAP | -0.02784 | -0.5217 to 0.4660 | 0.9988 |

Statistics for Figure 5E
cFos cell density

| ANOVA table | SS (Type III) | DF | MS | F (DFn, DFd) | P value | % of total variation |
|---|---|---|---|---|---|---|
| Genotype x Treatment | 89.11 | 1 | 89.11 | F (1, 51) = 8.039 | P=0.0065 | 12.83 |
| Genotype | 33.83 | 1 | 33.83 | F (1, 51) = 3.052 | P=0.0866 | 4.870 |
| Treatment | 0.2051 | 1 | 0.2051 | F (1, 51) = 0.01850 | P=0.8923 | 0.02952 |

| Tukey's multiple comparisons test | Predicted (LS) mean diff. | 95.00% CI of diff. | Adjusted P Value |
|---|---|---|---|
| Ctrl WT :Vehicle vs. Ctrl WT :SNAP | -2.706 | -5.069 to -0.3427 | 0.0257 |
| Ctrl WT :Vehicle vs. cKO:Vehicle | -4.173 | -6.725 to -1.620 | 0.0019 |
| Ctrl WT :Vehicle vs. cKO:SNAP | -1.715 | -4.333 to 0.9031 | 0.1944 |
| Ctrl WT :SNAP vs. cKO:Vehicle | -1.467 | -4.020 to 1.085 | 0.2539 |
| Ctrl WT :SNAP vs. cKO:SNAP | 0.9909 | -1.627 to 3.609 | 0.4508 |
| cKO:Vehicle vs. cKO:SNAP | 2.458 | -0.3320 to 5.248 | 0.0829 |

**Table S6**
Statistics for Figure 6C
Resting EEG AuC Delta

| | | |
|---|---|---|
| Mixed-<br>effects model (REML) | Matching: Across ro<br>w | |
| Assume sphericity? | Yes | |
| Alpha | 0.05 | |
| Fixed effects (type III) | P value | F (DFn, DFd) |
| Genotype | 0.0843 | F (1, 25) = 3.231 |
| Treatment | 0.1207 | F (1, 20) = 2.628 |
| Genotype x Treatment | 0.6578 | F (1, 20) = 0.2021 |
| Random effects | SD | Variance |
| Genotype | 55.24 | 3051 |
| Residual | 58.23 | 3391 |

| Uncorrected Fisher's LSD | Predicted (LS) mean diff. | 95.00% CI of diff. | Individual P Value |
|---|---|---|---|
| Ctrl WT | | | |
| Untreated vs. SNAP | 19.99 | -30.07 to 70.04 | 0.4148 |
| cKO | | | |
| Untreated vs. SNAP | 35.33 | -15.27 to 85.92 | 0.1608 |
| Untreated | | | |
| Ctrl WT vs. cKO | -56.58 | -120.8 to 7.600 | 0.0826 |
| SNAP | | | |
| Ctrl WT vs. cKO | -41.24 | -106.4 to 23.95 | 0.2092 |

Resting EEG AuC Theta

| Mixed-effects model (REML) | Matching: Across row | |
|---|---|---|
| Assume sphericity? | Yes | |
| Alpha | 0.05 | |
| Fixed effects (type III) | P value | F (DFn, DFd) |
| Genotype | 0.7406 | "F (1, 25) = 0.1121" |
| Treatment | 0.0378 | "F (1, 20) = 4.945" |
| Genotype x Treatment | 0.5456 | "F (1, 20) = 0.3780" |
| Random effects | SD | Variance |
| Genotype | 25.43 | 646.7 |
| Residual | 33.45 | 1119 |

| **Uncorrected Fisher's LSD** | **Predicted (LS) mean diff.** | **95.00% CI of diff.** | **Individual P Value** |
|---|---|---|---|
| Ctrl WT | | | |
| Untreated vs. SNAP | 27.66 | -0.9019 to 56.23 | 0.0570 |
| cKO | | | |
| Untreated vs. SNAP | 15.68 | -13.25 to 44.61 | 0.2717 |
| Untreated | | | |
| Ctrl WT vs. cKO | 1.376 | -32.34 to 35.09 | 0.9348 |
| SNAP | | | |
| Ctrl WT vs. cKO | -10.61 | -44.91 to 23.69 | 0.5365 |

Resting EEG AuC Alpha

| Mixed-effects model (REML) | Matching: Across row | |
|---|---|---|
| Assume sphericity? | Yes | |
| Alpha | 0.05 | |
| Fixed effects (type III) | P value | F (DFn, DFd) |
| Genotype | 0.4570 | "F (1, 25) = 0.5709" |
| Treatment | 0.0393 | "F (1, 19) = 4.899" |
| Genotype x Treatment | 0.7191 | "F (1, 19) = 0.1332" |
| Random effects | SD | Variance |
| Genotype | 11.63 | 135.2 |
| Residual | 9.952 | 99.04 |

Resting EEG AuC Beta

| | | |
|---|---|---|
| Mixed-effects model (REML) | Matching: Across ro w | |
| Assume sphericity? | Yes | |
| Alpha | 0.05 | |
| Fixed effects (type III) | P value | F (DFn, DFd) |
| Genotype | 0.1904 | "F (1, 25) = 1.812" |
| Treatment | 0.4304 | "F (1, 21) = 0.6464" |
| Genotype x Treatment | 0.2858 | "F (1, 21) = 1.200" |
| Random effects | SD | Variance |
| Genotype | 9.525 | 90.72 |
| Residual | 8.872 | 78.72 |

Resting EEG AuC Gamma

| | | |
|---|---|---|
| Mixed-effects model (REML) | Matching: Across ro w | |
| Assume sphericity? | Yes | |
| Alpha | 0.05 | |
| Fixed effects (type III) | P value | F (DFn, DFd) |
| Genotype | 0.5307 | "F (1, 25) = 0.4042" |
| Treatment | 0.7923 | "F (1, 22) = 0.07102" |
| Genotype x Treatment | 0.6802 | "F (1, 22) = 0.1744" |
| Random effects | SD | Variance |
| Genotype | 0.8000 | 0.6400 |
| Residual | 0.4786 | 0.2290 |

Resting EEG AuC Low Gamma

| | | |
|---|---|---|
| Mixed-effects model (REML) | Matching: Across ro w | |
| Assume sphericity? | Yes | |
| Alpha | 0.05 | |
| Fixed effects (type III) | P value | F (DFn, DFd) |
| Genotype | 0.7608 | "F (1, 25) = 0.09476" |
| Treatment | 0.5586 | "F (1, 22) = 0.3528" |

| | | |
|---|---|---|
| Genotype x Treatment | 0.7120 | "F (1, 22) = 0.1399" |
| Random effects | SD | Variance |
| Genotype | 1.773 | 3.144 |
| Residual | 0.9417 | 0.8868 |

Resting EEG AuC High Gamma

| | | |
|---|---|---|
| Mixed-effects model (REML) | Matching: Across ro w | |
| Assume sphericity? | Yes | |
| Alpha | 0.05 | |
| Fixed effects (type III) | P value | F (DFn, DFd) |
| Genotype | 0.1646 | "F (1, 25) = 2.050" |
| Treatment | 0.0137 | "F (1, 23) = 7.127" |
| Genotype x Treatment | 0.7191 | "F (1, 23) = 0.1325" |
| Random effects | SD | Variance |
| Genotype | 0.2998 | 0.08988 |
| Residual | 0.2411 | 0.05813 |

| **Uncorrected Fisher's LSD** | **Predicted (LS) mean diff.** | **95.00% CI of diff.** | **Individual P Value** |
|---|---|---|---|
| Ctrl WT | | | |
| Untreated vs. SNAP | -0.1562 | -0.3570 to 0.04454 | 0.1211 |
| cKO | | | |
| Untreated vs. SNAP | -0.2055 | -0.4012 to -0.009906 | 0.0403 |
| Untreated | | | |
| Ctrl WT vs. cKO | -0.1670 | -0.4650 to 0.1309 | 0.2652 |
| SNAP | | | |
| Ctrl WT vs. cKO | -0.2164 | -0.5218 to 0.08903 | 0.1608 |

Statistics for Figure 6D

Resting EEG FC Delta

| | | |
|---|---|---|
| Mixed-effects model (REML) | Matching: Across ro w | |
| Assume sphericity? | Yes | |
| Alpha | 0.05 | |
| Fixed effects (type III) | P value | F (DFn, DFd) |
| Genotype | 0.5610 | "F (1, 25) = 0.3472" |
| Treatment | 0.0002 | "F (1, 20) = 19.78" |

| | | |
|---|---|---|
| Genotype x Treatment | 0.7514 | "F (1, 20) = 0.1032" |
| Random effects | SD | Variance |
| Genotype | 69.86 | 4880 |
| Residual | 37.41 | 1399 |

| Uncorrected Fisher's LSD | Predicted (LS) mean diff. | 95.00% CI of diff. | Individual P Value |
|---|---|---|---|
| Ctrl WT | | | |
| Untreated vs. SNAP | 53.19 | 20.32 to 86.07 | 0.0030 |
| cKO | | | |
| Untreated vs. SNAP | 46.03 | 13.09 to 78.97 | 0.0086 |
| Untreated | | | |
| Ctrl WT vs. cKO | -13.57 | -75.57 to 48.43 | 0.6615 |
| SNAP | | | |
| Ctrl WT vs. cKO | -20.74 | -84.30 to 42.83 | 0.5145 |

Resting EEG FC Theta

| Mixed-effects model (REML) | Matching: Across ro w | |
|---|---|---|
| Assume sphericity? | Yes | |
| Alpha | 0.05 | |
| Fixed effects (type III) | P value | F (DFn, DFd) |
| Genotype | 0.4256 | "F (1, 25) = 0.6562" |
| Treatment | 0.0977 | "F (1, 21) = 3.003" |
| Genotype x Treatment | 0.5050 | "F (1, 21) = 0.4600" |
| Random effects | SD | Variance |
| Genotype | 33.71 | 1136 |
| Residual | 45.90 | 2106 |

Resting EEG FC Alpha

| Mixed-effects model (REML) | Matching: Across ro w | |
|---|---|---|
| Assume sphericity? | Yes | |
| Alpha | 0.05 | |
| Fixed effects (type III) | P value | F (DFn, DFd) |
| Genotype | 0.8962 | "F (1, 25) = 0.01736" |

| | | |
|---|---|---|
| Treatment | 0.1632 | "F (1, 21) = 2.089" |
| Genotype x Treatment | 0.9586 | "F (1, 21) = 0.002764" |
| Random effects | SD | Variance |
| Genotype | 10.91 | 119.0 |
| Residual | 13.60 | 184.9 |

Resting EEG FC Beta

| Mixed-effects model (REML) | Matching: Across row | |
|---|---|---|
| Assume sphericity? | Yes | |
| Alpha | 0.05 | |
| Fixed effects (type III) | P value | F (DFn, DFd) |
| Genotype | 0.2470 | "F (1, 25) = 1.405" |
| Treatment | 0.8721 | "F (1, 20) = 0.02658" |
| Genotype x Treatment | 0.7111 | "F (1, 20) = 0.1412" |
| Random effects | SD | Variance |
| Genotype | 8.561 | 73.29 |
| Residual | 6.863 | 47.10 |

Resting EEG FC Gamma

| Mixed-effects model (REML) | Matching: Across row | |
|---|---|---|
| Assume sphericity? | Yes | |
| Alpha | 0.05 | |
| Fixed effects (type III) | P value | F (DFn, DFd) |
| Genotype | 0.0133 | "F (1, 25) = 7.092" |
| Treatment | 0.9497 | "F (1, 22) = 0.004066" |
| Genotype x Treatment | 0.3389 | "F (1, 22) = 0.9557" |
| Random effects | SD | Variance |
| Genotype | 0.1806 | 0.03263 |
| Residual | 0.6077 | 0.3693 |

| **Uncorrected Fisher's LSD** | **Predicted (LS) mean diff.** | **95.00% CI of diff.** | **Individual P Value** |
|---|---|---|---|

| | | | |
|---|---|---|---|
| Ctrl WT | | | |
| Untreated vs. SNAP | 0.1784 | -0.3319 to 0.6886 | 0.4761 |
| cKO | | | |
| Untreated vs. SNAP | -0.1565 | -0.6508 to 0.3378 | 0.5182 |
| Untreated | | | |
| Ctrl WT vs. cKO | -0.3249 | -0.8161 to 0.1663 | 0.1898 |
| SNAP | | | |
| Ctrl WT vs. cKO | -0.6597 | -1.182 to -0.1375 | 0.0144 |

Resting EEG FC Low Gamma

| Mixed-effects model (REML) | Matching: Across row | |
|---|---|---|
| Assume sphericity? | Yes | |
| Alpha | 0.05 | |
| Fixed effects (type III) | P value | F (DFn, DFd) |
| Genotype | 0.0669 | "F (1, 25) = 3.671" |
| Treatment | 0.2863 | "F (1, 22) = 1.194" |
| Genotype x Treatment | 0.3482 | "F (1, 22) = 0.9187" |
| Random effects | SD | Variance |
| Genotype | 0.7075 | 0.5006 |
| Residual | 1.056 | 1.116 |

| **Uncorrected Fisher's LSD** | **Predicted (LS) mean diff.** | **95.00% CI of diff.** | **Individual P Value** |
|---|---|---|---|
| Ctrl WT | | | |
| Untreated vs. SNAP | 0.6151 | -0.2839 to 1.514 | 0.1699 |
| cKO | | | |
| Untreated vs. SNAP | 0.04030 | -0.8190 to 0.8996 | 0.9234 |
| Untreated | | | |
| Ctrl WT vs. cKO | -0.4889 | -1.474 to 0.4963 | 0.3232 |
| SNAP | | | |
| Ctrl WT vs. cKO | -1.064 | -2.106 to -0.02161 | 0.0456 |

Resting EEG FC High Gamma

| Mixed-effects model (REML) | Matching: Across row |
|---|---|
| Assume sphericity? | Yes |

| | | |
|---|---|---|
| Alpha | 0.05 | |
| Fixed effects (type III) | P value | F (DFn, DFd) |
| Genotype | 0.0075 | "F (1, 25) = 8.451" |
| Treatment | 0.0666 | "F (1, 23) = 3.709" |
| Genotype x Treatment | 0.3527 | "F (1, 23) = 0.9000" |
| Random effects | SD | Variance |
| Genotype | 0.05590 | 0.003125 |
| Residual | 0.4032 | 0.1625 |

| **Uncorrected Fisher's LSD** | **Predicted (LS) mean diff.** | **95.00% CI of diff.** | **Individual P Value** |
|---|---|---|---|
| Ctrl WT | | | |
| Untreated vs. SNAP | -0.1095 | -0.4378 to 0.2189 | 0.4973 |
| cKO | | | |
| Untreated vs. SNAP | -0.3220 | -0.6492 to 0.005102 | 0.0534 |
| Untreated | | | |
| Ctrl WT vs. cKO | -0.2254 | -0.5406 to 0.08986 | 0.1571 |
| SNAP | | | |
| Ctrl WT vs. cKO | -0.4379 | -0.7655 to -0.1103 | 0.0099 |

Statistics for Figure 6E
Resting EEG Power Coupling A1A2

| | | |
|---|---|---|
| Mixed-effects model (REML) | Matching: Across ro w | |
| Assume sphericity? | Yes | |
| Alpha | 0.05 | |
| Fixed effects (type III) | P value | F (DFn, DFd) |
| Genotype | 0.1087 | "F (1, 24) = 2.775" |
| Treatment | 0.5037 | "F (1, 23) = 0.4614" |
| Genotype x Treatment | 0.7014 | "F (1, 23) = 0.1507" |
| Random effects | SD | Variance |
| Genotype | 0.1160 | 0.01347 |
| Residual | 0.2059 | 0.04241 |

Resting EEG Power Coupling A1G1

| Mixed-effects model (REML) | Matching: Across ro w | |
|---|---|---|
| Assume sphericity? | Yes | |
| Alpha | 0.05 | |
| Fixed effects (type III) | P value | F (DFn, DFd) |
| Genotype | 0.1943 | "F (1, 48) = 1.733" |
| Treatment | 0.6402 | "F (1, 48) = 0.2212" |
| Genotype x Treatment | 0.1113 | "F (1, 48) = 2.632" |
| Random effects | SD | Variance |
| Genotype | 0.000 | 0.000 |
| Residual | 0.2004 | 0.04017 |

| **Uncorrected Fisher's LSD** | **Predicted (LS) mean diff.** | **95.00% CI of diff.** | **Individual P Value** |
|---|---|---|---|
| Ctrl WT | | | |
| Untreated vs. SNAP | -0.1165 | -0.2750 to 0.04203 | 0.1460 |
| cKO | | | |
| Untreated vs. SNAP | 0.06414 | -0.09393 to 0.2222 | 0.4186 |
| Untreated | | | |
| Ctrl WT vs. cKO | -0.1636 | -0.3188 to -0.008390 | 0.0393 |
| SNAP | | | |
| Ctrl WT vs. cKO | 0.01703 | -0.1443 to 0.1784 | 0.8328 |

Resting EEG Power Coupling A1G2

| Mixed-effects model (REML) | Matching: Across ro w | |
|---|---|---|
| Assume sphericity? | Yes | |
| Alpha | 0.05 | |
| Fixed effects (type III) | P value | F (DFn, DFd) |
| Genotype | 0.2120 | "F (1, 47) = 1.601" |
| Treatment | 0.9224 | "F (1, 47) = 0.009601" |
| Genotype x Treatment | 0.1856 | "F (1, 47) = 1.805" |
| Random effects | SD | Variance |
| Genotype | 0.000 | 0.000 |
| Residual | 0.1923 | 0.03699 |

| Uncorrected Fisher's LSD | Predicted (LS) mean diff. | 95.00% CI of diff. | Individual P Value |
|---|---|---|---|
| Ctrl WT | | | |
| Untreated vs. SNAP | -0.07780 | -0.2300 to 0.07442 | 0.3091 |
| cKO | | | |
| Untreated vs. SNAP | 0.06722 | -0.08767 to 0.2221 | 0.3871 |
| Untreated | | | |
| Ctrl WT vs. cKO | -0.1408 | -0.2898 to 0.008228 | 0.0635 |
| SNAP | | | |
| Ctrl WT vs. cKO | 0.004219 | -0.1537 to 0.1622 | 0.9574 |

Resting EEG Power Coupling A2G1

| Mixed-effects model (REML) | Matching: Across ro w | |
|---|---|---|
| Assume sphericity? | Yes | |
| Alpha | 0.05 | |
| Fixed effects (type III) | P value | F (DFn, DFd) |
| Genotype | 0.7139 | "F (1, 48) = 0.1360" |
| Treatment | 0.6980 | "F (1, 48) = 0.1523" |
| Genotype x Treatment | 0.9433 | "F (1, 48) = 0.005108" |
| Random effects | SD | Variance |
| Genotype | 0.000 | 0.000 |
| Residual | 0.2304 | 0.05311 |

Resting EEG Power Coupling A2G2

| Mixed-effects model (REML) | Matching: Across ro w | |
|---|---|---|
| Assume sphericity? | Yes | |
| Alpha | 0.05 | |
| Fixed effects (type III) | P value | F (DFn, DFd) |
| Genotype | 0.7856 | "F (1, 47) = 0.07482" |
| Treatment | 0.7915 | "F (1, 47) = 0.07068" |
| Genotype x Treatment | 0.9768 | "F (1, 47) = 0.0008511" |

| Random effects | SD | Variance |
|---|---|---|
| Genotype | 0.000 | 0.000 |
| Residual | 0.2056 | 0.04229 |

Statistics for Figure 6F

Resting EEG Power Coupling D1D2

| Mixed-effects model (REML) | Matching: Across ro w | |
|---|---|---|
| Assume sphericity? | Yes | |
| Alpha | 0.05 | |
| Fixed effects (type III) | P value | F (DFn, DFd) |
| Genotype | 0.5810 | "F (1, 24) = 0.3130" |
| Treatment | 0.0136 | "F (1, 23) = 7.149" |
| Genotype x Treatment | 0.6674 | "F (1, 23) = 0.1894" |
| Random effects | SD | Variance |
| Genotype | 0.1370 | 0.01876 |
| Residual | 0.1616 | 0.02612 |

| **Uncorrected Fisher's LSD** | **Predicted (LS) mean diff.** | **95.00% CI of diff.** | **Individual P Value** |
|---|---|---|---|
| Ctrl WT | | | |
| Untreated vs. SNAP | 0.1414 | 0.006446 to 0.2763 | 0.0408 |
| cKO | | | |
| Untreated vs. SNAP | 0.1018 | -0.02933 to 0.2329 | 0.1219 |
| Untreated | | | |
| Ctrl WT vs. cKO | -0.01958 | -0.1867 to 0.1476 | 0.8147 |
| SNAP | | | |
| Ctrl WT vs. cKO | -0.05917 | -0.2292 to 0.1108 | 0.4872 |

Resting EEG Power Coupling D1G1

| Mixed-effects model (REML) | Matching: Across ro w | |
|---|---|---|
| Assume sphericity? | Yes | |
| Alpha | 0.05 | |
| Fixed effects (type III) | P value | F (DFn, DFd) |
| Genotype | 0.1751 | "F (1, 25) = 1.948" |
| Treatment | 0.0729 | "F (1, 22) = 3.548" |
| Genotype x Treatment | 0.5842 | "F (1, 22) = 0.3085" |
| Random effects | SD | Variance |

| | | |
|---|---|---|
| Genotype | 0.06846 | 0.004686 |
| Residual | 0.2034 | 0.04138 |

Resting EEG Power Coupling D1G2

| Mixed-effects model (REML) | Matching: Across row | |
|---|---|---|
| Assume sphericity? | Yes | |
| Alpha | 0.05 | |
| Fixed effects (type III) | P value | F (DFn, DFd) |
| Genotype | 0.2647 | "F (1, 25) = 1.302" |
| Treatment | 0.0349 | "F (1, 22) = 5.053" |
| Genotype x Treatment | 0.3451 | "F (1, 22) = 0.9309" |
| Random effects | SD | Variance |
| Genotype | 0.09433 | 0.008898 |
| Residual | 0.1970 | 0.03880 |

| **Uncorrected Fisher's LSD** | **Predicted (LS) mean diff.** | **95.00% CI of diff.** | **Individual P Value** |
|---|---|---|---|
| Ctrl WT | | | |
| Untreated vs. SNAP | -0.07124 | -0.2366 to 0.09416 | 0.3814 |
| cKO | | | |
| Untreated vs. SNAP | -0.1784 | -0.3386 to -0.01814 | 0.0307 |
| Untreated | | | |
| Ctrl WT vs. cKO | 0.1292 | -0.04303 to 0.3014 | 0.1380 |
| SNAP | | | |
| Ctrl WT vs. cKO | 0.02206 | -0.1536 to 0.1977 | 0.8016 |

Resting EEG Power Coupling D2G1

| Mixed-effects model (REML) | Matching: Across row | |
|---|---|---|
| Assume sphericity? | Yes | |
| Alpha | 0.05 | |
| Fixed effects (type III) | P value | F (DFn, DFd) |
| Genotype | 0.3058 | "F (1, 48) = 1.071" |
| Treatment | 0.3165 | "F (1, 48) = 1.024" |
| Genotype x Treatment | 0.2989 | "F (1, 48) = 1.103" |

| Random effects | SD | Variance |
|---|---|---|
| Genotype | 0.000 | 0.000 |
| Residual | 0.2166 | 0.04692 |

Resting EEG Power Coupling D2G2

| Mixed-effects model (REML) | Matching: Across ro w | |
|---|---|---|
| Assume sphericity? | Yes | |
| Alpha | 0.05 | |
| Fixed effects (type III) | P value | F (DFn, DFd) |
| Genotype | 0.2274 | "F (1, 48) = 1.495" |
| Treatment | 0.0618 | "F (1, 48) = 3.659" |
| Genotype x Treatment | 0.5245 | "F (1, 48) = 0.4109" |
| Random effects | SD | Variance |
| Genotype | 0.000 | 0.000 |
| Residual | 0.2611 | 0.06818 |

| **Uncorrected Fisher's LSD** | **Predicted (LS) mean diff.** | **95.00% CI of diff.** | **Individual P Value** |
|---|---|---|---|
| Ctrl WT | | | |
| Untreated vs. SNAP | -0.09224 | -0.2988 to 0.1143 | 0.3737 |
| cKO | | | |
| Untreated vs. SNAP | -0.1852 | -0.3912 to 0.02070 | 0.0768 |
| Untreated | | | |
| Ctrl WT vs. cKO | 0.1352 | -0.06704 to 0.3374 | 0.1853 |
| SNAP | | | |
| Ctrl WT vs. cKO | 0.04219 | -0.1680 to 0.2524 | 0.6883 |

Statistics for Figure 6G

Resting EEG AuC Delta

| Mixed-effects model (REML) | Matching: Across ro w | |
|---|---|---|
| Assume sphericity? | Yes | |
| Alpha | 0.05 | |
| Fixed effects (type III) | P value | F (DFn, DFd) |
| Genotype | 0.6162 | F (1, 74) =0.2533 |

| | | |
|---|---|---|
| Treatment | 0.0020 | F (2, 45) = 7.166 |
| Genotype x Treatment | 0.1827 | F (2, 45) = 1.766 |
| Random effects | SD | Variance |
| Genotype | 45.29 | 2051 |
| Residual | 218.6 | 47785 |

| **Uncorrected Fisher's LSD** | **Predicted (LS) mean diff.** | **95.00% CI of diff.** | **Individual P Value** |
|---|---|---|---|
| WT | | | |
| Pre vs vehicle | -217.4 | -342.1 to -92.80 | 0.0010 |
| Pre vs SNAP | 116.4 | -34.12 to 266.9 | 0.1263 |
| Vehicle vs SNAP | 333.8 | 164.5 to 503.1 | 0.0003 |
| KO | | | |
| Pre vs vehicle | -49.50 | -191.1 to 92.09 | 0.4850 |
| Pre vs SNAP | 100.6 | -97.53 to 298.6 | 0.3121 |
| Vehicle vs SNAP | 150.1 | -66.47 to 366.6 | 0.1696 |
| Pre | | | |
| WT vs. KO | -25.39 | -127.7 to 76.91 | 0.6240 |
| Vehicle | | | |
| WT vs. KO | 142.5 | -14.90 to 300.0 | 0.0756 |
| SNAP | | | |
| WT vs. KO | -41.24 | -265.3 to 182.9 | 0.7162 |

Resting EEG AuC Theta

| | | |
|---|---|---|
| Mixed-effects model (REML) | Matching: Across row | |
| Assume sphericity? | Yes | |
| Alpha | 0.05 | |
| Fixed effects (type III) | P value | F (DFn, DFd) |
| Genotype | 0.7190 | F (1, 75) = 0.1305 |
| Treatment | 0.1290 | F (2, 44) = 2.147 |
| Genotype x Treatment | 0.4100 | F (2, 44) = 0.9099 |
| Random effects | SD | Variance |
| Genotype | 32.44 | 1052 |
| Residual | 55.69 | 3102 |

| **Uncorrected Fisher's LSD** | **Predicted (LS) mean diff.** | **95.00% CI of diff.** | **Individual P Value** |
|---|---|---|---|
| WT | | | |
| Pre vs vehicle | -26.11 | -59.22 to 6.993 | 0.1191 |

| | | | |
|---|---|---|---|
| Pre vs SNAP | 33.53 | -6.861 to 73.92 | 0.1014 |
| Vehicle vs SNAP | 59.64 | 15.65 to 103.6 | 0.0090 |
| KO | | | |
| Pre vs vehicle | -2.736 | -40.56 to 35.09 | 0.8848 |
| Pre vs SNAP | 10.98 | -42.47 to 64.44 | 0.6808 |
| Vehicle vs SNAP | 13.72 | -43.25 to 70.69 | 0.6298 |
| Pre | | | |
| WT vs. KO | 5.335 | -24.19 to 34.86 | 0.7211 |
| Vehicle | | | |
| WT vs. KO | 28.71 | -15.92 to 73.35 | 0.2053 |
| SNAP | | | |
| WT vs. KO | -17.21 | -79.66 to 45.24 | 0.5863 |

Resting EEG AuC Alpha

| Mixed-effects model (REML) | Matching: Across row | |
|---|---|---|
| Assume sphericity? | Yes | |
| Alpha | 0.05 | |
| Fixed effects (type III) | P value | F (DFn, DFd) |
| Genotype | 0.8209 | F (1, 119) = 0.05146 |
| Treatment | 0.2396 | F (2, 119) = 1.446 |
| Genotype x Treatment | 0.7925 | F (2, 119) = 0.2330 |
| Random effects | SD | Variance |
| Genotype | 0.000 | 0.000 |
| Residual | 36.15 | 1307 |

| Uncorrected Fisher's LSD | Predicted (LS) mean diff. | 95.00% CI of diff. | Individual P Value |
|---|---|---|---|
| WT | | | |

| | | | |
|---|---|---|---|
| Pre vs vehicle | -3.608 | -23.70 to 16.49 | 0.7229 |
| Pre vs SNAP | 14.55 | -9.637 to 38.73 | 0.2360 |
| Vehicle vs SNAP | 18.16 | -9.237 to 45.55 | 0.1919 |
| KO | | | |
| Pre vs vehicle | 6.769 | -16.06 to 29.60 | 0.5583 |
| Pre vs SNAP | 19.55 | -12.22 to 51.32 | 0.2254 |
| Vehicle vs SNAP | 12.78 | -22.15 to 47.71 | 0.4701 |
| Pre | | | |
| WT vs. KO | -3.316 | -19.88 to 13.25 | 0.6925 |
| Vehicle | | | |
| WT vs. KO | 7.060 | -18.45 to 32.57 | 0.5847 |
| SNAP | | | |
| WT vs. KO | 1.684 | -34.64 to 38.01 | 0.9270 |

Resting EEG AuC Beta

| Mixed-effects model (REML) | Matching: Across ro w | |
|---|---|---|
| Assume sphericity? | Yes | |
| Alpha | 0.05 | |
| Fixed effects (type III) | P value | F (DFn, DFd) |
| Genotype | 0.5375 | F (1, 119) = 0.3824 |
| Treatment | 0.1182 | F (2, 119) = 2.174 |
| Genotype x Treatment | 0.7026 | F (2, 119) = 0.3540 |
| Random effects | SD | Variance |
| Genotype | 0.000 | 0.000 |
| Residual | 5.919 | 35.03 |

| **Uncorrected Fisher's LSD** | **Predicted (LS) mean diff.** | **95.00% CI of diff.** | **Individual P Value** |
|---|---|---|---|

| | | | |
|---|---|---|---|
| WT | | | |
| Pre vs vehicle | 0.5528 | -2.737 to 3.843 | 0.7399 |
| Pre vs SNAP | 2.235 | -1.725 to 6.195 | 0.2660 |
| Vehicle vs SNAP | 1.682 | -2.803 to 6.167 | 0.4592 |
| KO | | | |
| Pre vs vehicle | 2.337 | -1.401 to 6.075 | 0.2182 |
| Pre vs SNAP | 4.219 | -0.9822 to 9.421 | 0.1109 |
| Vehicle vs SNAP | 1.882 | -3.836 to 7.601 | 0.5158 |
| Pre | | | |
| WT vs. KO | -2.064 | -4.776 to 0.6486 | 0.1346 |
| Vehicle | | | |
| WT vs. KO | -0.2796 | -4.456 to 3.897 | 0.8947 |
| SNAP | | | |
| WT vs. KO | -0.07937 | -6.027 to 5.869 | 0.9790 |

Resting EEG AuC Gamma

| Mixed-effects model (REML) | Matching: Across ro w | |
|---|---|---|
| Assume sphericity? | Yes | |
| Alpha | 0.05 | |
| Fixed effects (type III) | P value | F (DFn, DFd) |
| Genotype | 0.0129 | F (1, 119) = 6.375 |
| Treatment | 0.0560 | F (2, 119) = 2.954 |
| Genotype x Treatment | 0.8593 | F (2, 119) = 0.1518 |
| Random effects | SD | Variance |
| Genotype | 0.000 | 0.000 |
| Residual | 1.641 | 2.692 |

| Uncorrected Fisher's LSD | Predicted (LS) mean diff. | 95.00% CI of diff. | Individual P Value |
|---|---|---|---|
| WT | | | |
| Pre vs vehicle | 0.4022 | -0.5099 to 1.314 | 0.3844 |
| Pre vs SNAP | -0.6624 | -1.760 to 0.4353 | 0.2345 |
| Vehicle vs SNAP | -1.065 | -2.308 to 0.1788 | 0.0926 |
| KO | | | |
| Pre vs vehicle | 0.2369 | -0.7993 to 1.273 | 0.6516 |
| Pre vs SNAP | -1.155 | -2.597 to 0.2870 | 0.1154 |
| Vehicle vs SNAP | -1.392 | -2.977 to 0.1935 | 0.0847 |
| Pre | | | |
| WT vs. KO | -0.6948 | -1.447 to 0.0570 | 0.0698 |
| Vehicle | | | |
| WT vs. KO | -0.8600 | -2.018 to 0.2977 | 0.1439 |
| SNAP | | | |
| WT vs. KO | -1.187 | -2.836 to 0.4616 | 0.1566 |

Resting EEG AuC Low Gamma

| Mixed-effects model (REML) | Matching: Across row | |
|---|---|---|
| Assume sphericity? | Yes | |
| Alpha | 0.05 | |
| Fixed effects (type III) | P value | F (DFn, DFd) |
| Genotype | 0.0038 | F (1, 119) = 8.738 |
| Treatment | 0.0354 | F (2, 119) = 3.435 |
| Genotype x Treatment | 0.9445 | F (2, 119) = 0.05708 |
| Random effects | SD | Variance |
| Genotype | 0.000 | 0.000 |
| Residual | 2.011 | 4.043 |

| Uncorrected Fisher's LSD | Predicted (LS) mean diff. | 95.00% CI of diff. | Individual P Value |
|---|---|---|---|
| WT | | | |
| Pre vs vehicle | 0.8951 | -0.2226 to 2.013 | 0.1155 |
| Pre vs SNAP | -0.5597 | -1.905 to 0.7857 | 0.4117 |
| Vehicle vs SNAP | -1.455 | -2.979 to 0.0689 | 0.0611 |
| KO | | | |
| Pre vs vehicle | 1.075 | -0.1949 to 2.345 | 0.0963 |
| Pre vs SNAP | -0.2176 | -1.985 to 1.550 | 0.8078 |
| Vehicle vs SNAP | -1.293 | -3.235 to 0.6502 | 0.1902 |
| Pre | | | |
| WT vs. KO | -1.485 | -2.407 to -0.5640 | 0.0018 |
| Vehicle | | | |
| WT vs. KO | -1.306 | -2.724 to 0.1133 | 0.0710 |
| SNAP | | | |
| WT vs. KO | -1.143 | -3.164 to 0.8774 | 0.2648 |

Resting EEG AuC High Gamma

| Mixed-effects model (REML) | Matching: Across ro w | |
|---|---|---|
| Assume sphericity? | Yes | |
| Alpha | 0.05 | |
| Fixed effects (type III) | P value | F (DFn, DFd) |
| Genotype | 0.0686 | F (1, 119) = 3.378 |
| Treatment | 0.0194 | F (2, 119) = 4.078 |
| Genotype x Treatment | 0.5022 | F (2, 119) = 0.6928 |
| Random effects | SD | Variance |
| Genotype | 0.000 | 0.000 |

| | | |
|---|---|---|
| Residual | 1.832 | 3.355 |

| Uncorrected Fisher's LSD | Predicted (LS) mean diff. | 95.00% CI of diff. | Individual P Value |
|---|---|---|---|
| WT | | | |
| Pre vs vehicle | 0.1387 | -0.8796 to 1.157 | 0.7879 |
| Pre vs SNAP | -0.8663 | -2.092 to 0.3593 | 0.1642 |
| Vehicle vs SNAP | -1.005 | -2.393 to 0.3832 | 0.1543 |
| KO | | | |
| Pre vs vehicle | -0.3137 | -1.471 to 0.8431 | 0.5923 |
| Pre vs SNAP | -2.019 | -3.629 to -0.4096 | 0.0144 |
| Vehicle vs SNAP | -1.706 | -3.475 to 0.0642 | 0.0588 |
| Pre | | | |
| WT vs. KO | -0.2077 | -1.047 to 0.6317 | 0.6251 |
| Vehicle | | | |
| WT vs. KO | -0.6601 | -1.953 to 0.6324 | 0.3140 |
| SNAP | | | |
| WT vs. KO | -1.361 | -3.202 to 0.4801 | 0.1459 |

Statistics for Figure 6H
Resting EEG FC Delta

| Mixed-effects model (REML) | Matching: Across row | |
|---|---|---|
| Assume sphericity? | Yes | |
| Alpha | 0.05 | |
| Fixed effects (type III) | P value | F (DFn, DFd) |
| Genotype | 0.6193 | F (1, 75) = 0.2488 |
| Treatment | 0.1799 | F (2, 44) = 1.784 |
| Genotype x Treatment | 0.5014 | F (2, 44) = 0.7012 |

| Random effects | SD | Variance |
| --- | --- | --- |
| Genotype | 13.11 | 171.8 |
| Residual | 264.1 | 69771 |

| Uncorrected Fisher's LSD | Predicted (LS) mean diff. | 95.00% CI of diff. | Individual P Value |
| --- | --- | --- | --- |
| WT | | | |
| Pre vs vehicle | -125.4 | -274.9 to 24.17 | 0.0982 |
| Pre vs SNAP | 106.4 | -73.56 to 286.4 | 0.2397 |
| Vehicle vs SNAP | 231.8 | 28.00 to 435.6 | 0.0267 |
| KO | | | |
| Pre vs vehicle | -54.80 | -224.7 to 115.1 | 0.5190 |
| Pre vs SNAP | -16.75 | -253.2 to 219.7 | 0.8871 |
| Vehicle vs SNAP | 38.05 | -221.8 to 297.9 | 0.7694 |
| Pre | | | |
| WT vs. KO | 46.69 | -74.50 to 167.9 | 0.4470 |
| Vehicle | | | |
| WT vs. KO | 117.2 | -69.36 to 303.9 | 0.2159 |
| SNAP | | | |
| WT vs. KO | -76.50 | -342.3 to 189.3 | 0.5698 |

Resting EEG FC Theta

| Mixed-effects model (REML) | Matching: Across row | |
| --- | --- | --- |
| Assume sphericity? | Yes | |
| Alpha | 0.05 | |
| Fixed effects (type III) | P value | F (DFn, DFd) |
| Genotype | 0.2332 | F (1, 75) = 1.445 |
| Treatment | 0.7909 | F (2, 44) = 0.2358 |
| Genotype x Treatment | 0.4437 | F (2, 44) = 0.8278 |

| Random effects | SD | Variance |
|---|---|---|
| Genotype | 8.831 | 77.99 |
| Residual | 92.77 | 8606 |

| **Uncorrected Fisher's LSD** | **Predicted (LS) mean diff.** | **95.00% CI of diff.** | **Individual P Value** |
|---|---|---|---|
| WT | | | |
| Pre vs vehicle | 8.580 | -44.00 to 61.16 | 0.7438 |
| Pre vs SNAP | 31.80 | -31.54 to 95.14 | 0.3171 |
| Vehicle vs SNAP | 23.22 | -48.41 to 94.85 | 0.5170 |
| KO | | | |
| Pre vs vehicle | 18.22 | -41.54 to 77.97 | 0.5421 |
| Pre vs SNAP | -29.16 | -112.4 to 54.07 | 0.4839 |
| Vehicle vs SNAP | -47.37 | -138.8 to 44.02 | 0.3019 |
| Pre | | | |
| WT vs. KO | 41.94 | -0.7673 to 84.64 | 0.0542 |
| Vehicle | | | |
| WT vs. KO | 51.57 | -14.18 to 117.3 | 0.1231 |
| SNAP | | | |
| WT vs. KO | -19.02 | -112.7 to 74.62 | 0.6882 |

Resting EEG FC Alpha

| Mixed-effects model (REML) | Matching: Across row | |
|---|---|---|
| Assume sphericity? | Yes | |
| Alpha | 0.05 | |
| Fixed effects (type III) | P value | F (DFn, DFd) |
| Genotype | 0.0859 | F (1, 119) = 2.999 |
| Treatment | 0.8998 | F (2, 119) = 0.1056 |

| | | |
|---|---|---|
| Genotype x Treatment | 0.7683 | F (2, 119) = 0.2642 |
| Random effects | SD | Variance |
| Genotype | 0.000 | 0.000 |
| Residual | 34.61 | 1198 |

| **Uncorrected Fisher's LSD** | **Predicted (LS) mean diff.** | **95.00% CI of diff.** | **Individual P Value** |
|---|---|---|---|
| WT | | | |
| Pre vs vehicle | 3.196 | -16.04 to 22.43 | 0.7428 |
| Pre vs SNAP | 9.316 | -13.84 to 32.47 | 0.4272 |
| Vehicle vs SNAP | 6.120 | -20.11 to 32.35 | 0.6449 |
| KO | | | |
| Pre vs vehicle | 3.124 | -18.73 to 24.98 | 0.7777 |
| Pre vs SNAP | -4.375 | -34.79 to 26.04 | 0.7763 |
| Vehicle vs SNAP | -7.498 | -40.94 to 25.94 | 0.6578 |
| Pre | | | |
| WT vs. KO | 17.81 | 1.952 to 33.67 | 0.0281 |
| Vehicle | | | |
| WT vs. KO | 17.74 | -6.681 to 42.16 | 0.1530 |
| SNAP | | | |
| WT vs. KO | 4.121 | -30.66 to 38.90 | 0.8149 |

Resting EEG FC Beta

| | | |
|---|---|---|
| Mixed-effects model (REML) | Matching: Across ro w | |
| Assume sphericity? | Yes | |
| Alpha | 0.05 | |
| Fixed effects (type III) | P value | F (DFn, DFd) |
| Genotype | 0.2759 | F (1, 119) = 1.198 |

| | | |
|---|---|---|
| Treatment | 0.1068 | F (2, 119) = 2.279 |
| Genotype x Treatment | 0.8699 | F (2, 119) = 0.1395 |
| Random effects | SD | Variance |
| Genotype | 0.000 | 0.000 |
| Residual | 5.071 | 25.71 |

| **Uncorrected Fisher's LSD** | **Predicted (LS) mean diff.** | **95.00% CI of diff.** | **Individual P Value** |
|---|---|---|---|
| WT | | | |
| Pre vs vehicle | 1.744 | -1.075 to 4.562 | 0.2230 |
| Pre vs SNAP | 0.6937 | -2.699 to 4.086 | 0.6863 |
| Vehicle vs SNAP | -1.050 | -4.892 to 2.793 | 0.5895 |
| KO | | | |
| Pre vs vehicle | 2.856 | -0.3462 to 6.059 | 0.0800 |
| Pre vs SNAP | 0.7175 | -3.738 to 5.174 | 0.7504 |
| Vehicle vs SNAP | -2.139 | -7.038 to 2.760 | 0.3891 |
| Pre | | | |
| WT vs. KO | 0.8458 | -1.478 to 3.169 | 0.4725 |
| Vehicle | | | |
| WT vs. KO | 1.958 | -1.619 to 5.536 | 0.2806 |
| SNAP | | | |
| WT vs. KO | 0.8696 | -4.226 to 5.965 | 0.7360 |

Resting EEG FC Gamma

| | | |
|---|---|---|
| Mixed-effects model (REML) | Matching: Across ro w | |
| Assume sphericity? | Yes | |
| Alpha | 0.05 | |
| Fixed effects (type III) | P value | F (DFn, DFd) |

| | | |
|---|---|---|
| Genotype | 0.1236 | F (1, 119) = 2.405 |
| Treatment | <0.0001 | F (2, 119) = 11.69 |
| Genotype x Treatment | 0.8804 | F (2, 119) = 0.1275 |
| Random effects | SD | Variance |
| Genotype | 0.000 | 0.000 |
| Residual | 1.087 | 1.182 |

| Uncorrected Fisher's LSD | Predicted (LS) mean diff. | 95.00% CI of diff. | Individual P Value |
|---|---|---|---|
| WT | | | |
| Pre vs vehicle | 0.7656 | 0.1612 to 1.370 | 0.0135 |
| Pre vs SNAP | -0.6325 | -1.360 to 0.0948 | 0.0877 |
| Vehicle vs SNAP | -1.398 | -2.222 to -0.5742 | 0.0010 |
| KO | | | |
| Pre vs vehicle | 0.9971 | 0.3105 to 1.684 | 0.0048 |
| Pre vs SNAP | -0.5997 | -1.555 to 0.3556 | 0.2163 |
| Vehicle vs SNAP | -1.597 | -2.647 to -0.5465 | 0.0032 |
| Pre | | | |
| WT vs. KO | -0.4601 | -0.9582 to 0.038 | 0.0700 |
| Vehicle | | | |
| WT vs. KO | -0.2285 | -0.9956 to 0.538 | 0.5564 |
| SNAP | | | |
| WT vs. KO | -0.4273 | -1.520 to 0.6652 | 0.4402 |

Resting EEG FC Low Gamma

| | |
|---|---|
| Mixed-effects model (REML) | Matching: Across ro w |
| Assume sphericity? | Yes |
| Alpha | 0.05 |

| Fixed effects (type III) | P value | F (DFn, DFd) |
|---|---|---|
| Genotype | 0.0383 | F (1, 119) = 4.386 |
| Treatment | <0.0001 | F (2, 119) = 11.53 |
| Genotype x Treatment | 0.6738 | F (2, 119) = 0.3961 |
| Random effects | SD | Variance |
| Genotype | 0.000 | 0.000 |
| Residual | 1.578 | 2.489 |

| **Uncorrected Fisher's LSD** | **Predicted (LS) mean diff.** | **95.00% CI of diff.** | **Individual P Value** |
|---|---|---|---|
| WT | | | |
| Pre vs vehicle | 1.204 | 0.3271 to 2.081 | 0.0075 |
| Pre vs SNAP | -0.7085 | -1.764 to 0.3471 | 0.1864 |
| Vehicle vs SNAP | -1.913 | -3.108 to -0.7170 | 0.0020 |
| KO | | | |
| Pre vs vehicle | 1.715 | 0.7182 to 2.711 | 0.0009 |
| Pre vs SNAP | -0.1619 | -1.548 to 1.225 | 0.8175 |
| Vehicle vs SNAP | -1.877 | -3.401 to -0.3521 | 0.0163 |
| Pre | | | |
| WT vs. KO | -1.081 | -1.804 to -0.3584 | 0.0037 |
| Vehicle | | | |
| WT vs. KO | -0.5710 | -1.684 to 0.5423 | 0.3119 |
| SNAP | | | |
| WT vs. KO | -0.5348 | -2.120 to 1.051 | 0.5055 |

Resting EEG FC High Gamma

| Mixed-effects model (REML) | Matching: Across ro w |
|---|---|
| Assume sphericity? | Yes |

| | | |
|---|---|---|
| Alpha | 0.05 | |
| Fixed effects (type III) | P value | F (DFn, DFd) |
| Genotype | 0.6068 | F (1, 119) = 0.2662 |
| Treatment | 0.0002 | F (2, 119) = 9.123 |
| Genotype x Treatment | 0.8825 | F (2, 119) = 0.1252 |
| Random effects | SD | Variance |
| Genotype | 0.000 | 0.000 |
| Residual | 1.072 | 1.148 |

| Uncorrected Fisher's LSD | Predicted (LS) mean diff. | 95.00% CI of diff. | Individual P Value |
|---|---|---|---|
| WT | | | |
| Pre vs vehicle | 0.5657 | -0.0299 to 1.161 | 0.0625 |
| Pre vs SNAP | -0.6758 | -1.393 to 0.0411 | 0.0644 |
| Vehicle vs SNAP | -1.241 | -2.053 to -0.4295 | 0.0030 |
| KO | | | |
| Pre vs vehicle | 0.6217 | -0.0550 to 1.298 | 0.0714 |
| Pre vs SNAP | -0.9415 | -1.883 to 0.0001 | 0.0500 |
| Vehicle vs SNAP | -1.563 | -2.599 to -0.5280 | 0.0034 |
| Pre | | | |
| WT vs. KO | -0.05209 | -0.5431 to 0.438 | 0.8340 |
| Vehicle | | | |
| WT vs. KO | 0.003962 | -0.7521 to 0.760 | 0.9917 |
| SNAP | | | |
| WT vs. KO | -0.3178 | -1.395 to 0.7590 | 0.5600 |

Statistics for Figure 6I
Resting EEG Power Coupling A1A2

| ANOVA results | P value | F (DFn, DFd) |
|---|---|---|
| Genotype | 0.7863 | F (1, 119) = 0.07386 |
| Treatment | 0.0003 | F (2, 119) = 8.793 |
| Genotype x Treatment | 0.9636 | F (2, 119) = 0.03704 |

| Uncorrected Fisher's LSD | Predicted (LS) mean diff. | 95.00% CI of diff. | Individual P Value |
|---|---|---|---|
| WT:Pre vs. WT:Veh | -0.05028 | -0.1740 to 0.0734 | 0.4227 |

| | | | |
|---|---|---|---|
| WT:Pre vs. WT:SNAP | 0.2271 | 0.07811 to 0.3760 | 0.0031 |
| WT:Pre vs. KO:Pre | -0.02803 | -0.1300 to 0.0739 | 0.5875 |
| WT:Pre vs. KO:Veh | -0.05489 | -0.1905 to 0.0807 | 0.4246 |
| WT:Pre vs. KO:SNAP | 0.2196 | 0.02753 to 0.411 | 0.0254 |
| WT:Veh vs. WT:SNAP | 0.2773 | 0.1086 to 0.4460 | 0.0015 |
| WT:Veh vs. KO:Pre | 0.02226 | -0.1069 to 0.151 | 0.7336 |
| WT:Veh vs. KO:Veh | -0.004602 | -0.1617 to 0.1525 | 0.9538 |
| WT:Veh vs. KO:SNAP | 0.2699 | 0.06212 to 0.477 | 0.0113 |
| WT:SNAP vs. KO:Pre | -0.2551 | -0.408 to -0.1016 | 0.0013 |
| WT:SNAP vs. KO:Veh | -0.2819 | -0.4595 to -0.104 | 0.0021 |
| WT:SNAP vs. KO:SNAP | -0.007418 | -0.2311 to 0.2163 | 0.9478 |
| KO:Pre vs. KO:Veh | -0.02686 | -0.1675 to 0.113 | 0.7059 |
| KO:Pre vs. KO:SNAP | 0.2477 | 0.0520 to 0.4433 | 0.0135 |
| KO:Veh vs. KO:SNAP | 0.2745 | 0.05942 to 0.489 | 0.0128 |

Resting EEG Power Coupling A1G1

| ANOVA results | P value | F (DFn, DFd) |
|---|---|---|
| Genotype | 0.3554 | F (2, 119) = 1.801 |
| Treatment | 0.0880 | F (2, 119) = 2.481 |
| Genotype x Treatment | 0.1696 | F (1, 119) = 0.8606 |

| **Uncorrected Fisher's LSD** | **Predicted (LS) mean diff.** | **95.00% CI of diff.** | **Individual P Value** |
|---|---|---|---|
| WT:Pre vs. WT:Veh | -0.1254 | -0.2579 to 0.0070 | 0.0633 |
| WT:Pre vs. WT:SNAP | -0.2274 | -0.386 to -0.0680 | 0.0056 |
| WT:Pre vs. KO:Pre | -0.1513 | -0.260 to -0.0421 | 0.0070 |
| WT:Pre vs. KO:Veh | -0.2207 | -0.3659 to -0.075 | 0.0032 |
| WT:Pre vs. KO:SNAP | -0.1271 | -0.3327 to 0.078 | 0.2234 |
| WT:Veh vs. WT:SNAP | -0.1020 | -0.2826 to 0.078 | 0.2655 |
| WT:Veh vs. KO:Pre | -0.02594 | -0.1642 to 0.112 | 0.7109 |
| WT:Veh vs. KO:Veh | -0.09532 | -0.263 to 0.0728 | 0.2639 |
| WT:Veh vs. KO:SNAP | -0.001684 | -0.2241 to 0.220 | 0.9881 |
| WT:SNAP vs. KO:Pre | 0.07609 | -0.0881 to 0.240 | 0.3609 |
| WT:SNAP vs. KO:Veh | 0.006707 | -0.1834 to 0.196 | 0.9444 |
| WT:SNAP vs. KO:SNAP | 0.1003 | -0.1391 to 0.339 | 0.4084 |
| KO:Pre vs. KO:Veh | -0.06938 | -0.219 to 0.0811 | 0.3632 |
| KO:Pre vs. KO:SNAP | 0.02426 | -0.1852 to 0.233 | 0.8190 |
| KO:Veh vs. KO:SNAP | 0.09364 | -0.136 to 0.3239 | 0.4222 |

Resting EEG Power Coupling A1G2

| ANOVA results | P value | F (DFn, DFd) |
|---|---|---|
| Genotype | 0.9942 | F (1, 119) = 5.240e-005 |
| Treatment | 0.2164 | F (2, 119) = 1.551 |
| Genotype x Treatmen t | 0.2406 | F (2, 119) = 1.442 |

| **Uncorrected Fisher's LSD** | **Predicted (LS) mean diff.** | **95.00% CI of diff.** | **Individual P Value** |
|---|---|---|---|
| WT:Pre vs. WT:Veh | -0.1246 | -0.2603 to 0.011 | 0.0716 |
| WT:Pre vs. WT:SNAP | -0.1897 | -0.3530 to -0.026 | 0.0232 |
| WT:Pre vs. KO:Pre | -0.1035 | -0.2154 to 0.008 | 0.0694 |
| WT:Pre vs. KO:Veh | -0.1398 | -0.2886 to 0.008 | 0.0651 |
| WT:Pre vs. KO:SNAP | -0.07211 | -0.2828 to 0.138 | 0.4992 |
| WT:Veh vs. WT:SNAP | -0.06513 | -0.2501 to 0.119 | 0.4871 |
| WT:Veh vs. KO:Pre | 0.02107 | -0.1206 to 0.162 | 0.7688 |
| WT:Veh vs. KO:Veh | -0.01526 | -0.1875 to 0.157 | 0.8611 |
| WT:Veh vs. KO:SNAP | 0.05247 | -0.1754 to 0.280 | 0.6492 |
| WT:SNAP vs. KO:Pre | 0.08620 | -0.0821 to 0.254 | 0.3126 |
| WT:SNAP vs. KO:Veh | 0.04987 | -0.1449 to 0.244 | 0.6131 |
| WT:SNAP vs. KO:SNAP | 0.1176 | -0.1277 to 0.362 | 0.3445 |
| KO:Pre vs. KO:Veh | -0.03633 | -0.1905 to 0.117 | 0.6417 |
| KO:Pre vs. KO:SNAP | 0.03140 | -0.1831 to 0.245 | 0.7725 |
| KO:Veh vs. KO:SNAP | 0.06773 | -0.1681 to 0.303 | 0.5707 |

Resting EEG Power Coupling A2G1

| ANOVA results | P value | F (DFn, DFd) |
|---|---|---|
| Genotype | 0.0459 | F (1, 119) = 4.070 |
| Treatment | 0.0377 | F (2, 119) = 3.370 |
| Genotype x Treatmen t | 0.6731 | F (2, 119) = 0.3972 |

| Uncorrected Fisher's LSD | Predicted (LS) mean diff. | 95.00% CI of diff. | Individual P Value |
|---|---|---|---|
| WT:Pre vs. WT:Veh | -0.1223 | -0.2349 to -0.009 | 0.0336 |
| WT:Pre vs. WT:SNAP | -0.1437 | -0.2793 to -0.008 | 0.0380 |
| WT:Pre vs. KO:Pre | -0.1366 | -0.2295 to -0.043 | 0.0043 |
| WT:Pre vs. KO:Veh | -0.2092 | -0.3327 to -0.085 | 0.0011 |
| WT:Pre vs. KO:SNAP | -0.1908 | -0.3656 to -0.015 | 0.0328 |
| WT:Veh vs. WT:SNAP | -0.02142 | -0.1750 to 0.132 | 0.7829 |
| WT:Veh vs. KO:Pre | -0.01433 | -0.1319 to 0.103 | 0.8097 |
| WT:Veh vs. KO:Veh | -0.08693 | -0.2299 to 0.056 | 0.2310 |
| WT:Veh vs. KO:SNAP | -0.06848 | -0.2576 to 0.120 | 0.4748 |
| WT:SNAP vs. KO:Pre | 0.007086 | -0.1326 to 0.146 | 0.9202 |
| WT:SNAP vs. KO:Veh | -0.06551 | -0.2272 to 0.096 | 0.4239 |
| WT:SNAP vs. KO:SNAP | -0.04707 | -0.2507 to 0.156 | 0.6480 |
| KO:Pre vs. KO:Veh | -0.07259 | -0.2006 to 0.055 | 0.2636 |
| KO:Pre vs. KO:SNAP | -0.05415 | -0.2322 to 0.123 | 0.5482 |
| KO:Veh vs. KO:SNAP | 0.01844 | -0.1773 to 0.214 | 0.8524 |

Resting EEG Power Coupling A2G2

| ANOVA results | P value | F (DFn, DFd) |
|---|---|---|
| Genotype | 0.0619 | F (1, 119) = 3.553 |
| Treatment | 0.0357 | F (2, 119) = 3.428 |
| Genotype x Treatment | 0.8017 | F (2, 119) = 0.2215 |

| Uncorrected Fisher's LSD | Predicted (LS) mean diff. | 95.00% CI of diff. | Individual P Value |
|---|---|---|---|
| WT:Pre vs. WT:Veh | -0.1547 | -0.2805 to -0.028 | 0.0163 |
| WT:Pre vs. WT:SNAP | -0.08529 | -0.2367 to 0.066 | 0.2669 |
| WT:Pre vs. KO:Pre | -0.1280 | -0.2317 to -0.024 | 0.0160 |
| WT:Pre vs. KO:Veh | -0.2210 | -0.3589 to -0.083 | 0.0019 |
| WT:Pre vs. KO:SNAP | -0.1734 | -0.3686 to 0.021 | 0.0813 |
| WT:Veh vs. WT:SNAP | 0.06944 | -0.1020 to 0.240 | 0.4242 |
| WT:Veh vs. KO:Pre | 0.02674 | -0.1045 to 0.158 | 0.6874 |
| WT:Veh vs. KO:Veh | -0.06627 | -0.2259 to 0.093 | 0.4128 |
| WT:Veh vs. KO:SNAP | -0.01864 | -0.2299 to 0.192 | 0.8616 |
| WT:SNAP vs. KO:Pre | -0.04270 | -0.1987 to 0.113 | 0.5888 |
| WT:SNAP vs. KO:Veh | -0.1357 | -0.3162 to 0.044 | 0.1393 |
| WT:SNAP vs. KO:SNAP | -0.08808 | -0.3155 to 0.139 | 0.4446 |
| KO:Pre vs. KO:Veh | -0.09301 | -0.2359 to 0.049 | 0.2000 |
| KO:Pre vs. KO:SNAP | -0.04538 | -0.2442 to 0.153 | 0.6522 |

| | | | |
|---|---|---|---|
| KO:Veh vs. KO:SNAP | 0.04763 | -0.1710 to 0.266 | 0.6670 |

Statistics for Figure 6J

Resting EEG Power Coupling D1D2

| ANOVA results | P value | F (DFn, DFd) |
|---|---|---|
| Genotype | 0.4164 | F (1, 119) = 0.6651 |
| Treatment | 0.0218 | F (2, 119) = 3.954 |
| Genotype x Treatment | 0.9727 | F (2, 119) = 0.02772 |

| **Uncorrected Fisher's LSD** | **Predicted (LS) mean diff.** | **95.00% CI of diff.** | **Individual P Value** |
|---|---|---|---|
| WT:Pre vs. WT:Veh | -0.05762 | -0.1916 to 0.076 | 0.3961 |
| WT:Pre vs. WT:SNAP | 0.1644 | 0.00315 to 0.325 | 0.0457 |
| WT:Pre vs. KO:Pre | -0.03052 | -0.141 to 0.0799 | 0.5852 |
| WT:Pre vs. KO:Veh | -0.09507 | -0.2419 to 0.051 | 0.2023 |
| WT:Pre vs. KO:SNAP | 0.1023 | -0.1057 to 0.310 | 0.3321 |
| WT:Veh vs. WT:SNAP | 0.2220 | 0.0393 to 0.4046 | 0.0176 |
| WT:Veh vs. KO:Pre | 0.02710 | -0.1127 to 0.166 | 0.7019 |
| WT:Veh vs. KO:Veh | -0.03745 | -0.2075 to 0.132 | 0.6635 |
| WT:Veh vs. KO:SNAP | 0.1599 | -0.0650 to 0.384 | 0.1619 |
| WT:SNAP vs. KO:Pre | -0.1949 | -0.3611 to -0.028 | 0.0219 |
| WT:SNAP vs. KO:Veh | -0.2595 | -0.4517 to -0.067 | 0.0086 |
| WT:SNAP vs. KO:SNAP | -0.06211 | -0.304 to 0.1801 | 0.6125 |
| KO:Pre vs. KO:Veh | -0.06455 | -0.2168 to 0.087 | 0.4027 |
| KO:Pre vs. KO:SNAP | 0.1328 | -0.0789 to 0.344 | 0.2168 |
| KO:Veh vs. KO:SNAP | 0.1974 | -0.0354 to 0.430 | 0.0959 |

Resting EEG Power Coupling D1G1

| ANOVA results | P value | F (DFn, DFd) |
|---|---|---|
| Genotype | 0.7599 | F (1, 119) = 0.09383 |
| Treatment | 0.0054 | F (2, 119) = 5.455 |
| Genotype x Treatment | 0.2659 | F (2, 119) = 1.339 |

| Uncorrected Fisher's LSD | Predicted (LS) mean diff. | 95.00% CI of diff. | Individual P Value |
|---|---|---|---|
| WT:Pre vs. WT:Veh | -0.2650 | -0.4123 to -0.117 | 0.0005 |
| WT:Pre vs. WT:SNAP | -0.1357 | -0.313 to 0.0417 | 0.1326 |
| WT:Pre vs. KO:Pre | -0.09092 | -0.212 to 0.0305 | 0.1410 |
| WT:Pre vs. KO:Veh | -0.1980 | -0.3596 to -0.036 | 0.0167 |
| WT:Pre vs. KO:SNAP | -0.05791 | -0.2867 to 0.170 | 0.6171 |
| WT:Veh vs. WT:SNAP | 0.1293 | -0.0716 to 0.330 | 0.2050 |
| WT:Veh vs. KO:Pre | 0.1740 | 0.0202 to 0.3279 | 0.0269 |
| WT:Veh vs. KO:Veh | 0.06693 | -0.1201 to 0.254 | 0.4801 |
| WT:Veh vs. KO:SNAP | 0.2070 | -0.0404 to 0.454 | 0.1002 |
| WT:SNAP vs. KO:Pre | 0.04474 | -0.138 to 0.2275 | 0.6288 |
| WT:SNAP vs. KO:Veh | -0.06237 | -0.273 to 0.1492 | 0.5604 |
| WT:SNAP vs. KO:SNAP | 0.07775 | -0.1887 to 0.344 | 0.5645 |
| KO:Pre vs. KO:Veh | -0.1071 | -0.2746 to 0.060 | 0.2078 |
| KO:Pre vs. KO:SNAP | 0.03301 | -0.200 to 0.2660 | 0.7795 |
| KO:Veh vs. KO:SNAP | 0.1401 | -0.116 to 0.3963 | 0.2810 |

Resting EEG Power Coupling D1G2

| ANOVA results | P value | F (DFn, DFd) |
|---|---|---|
| Genotype | 0.5529 | F (1, 119) = 0.3542 |
| Treatment | 0.0011 | F (2, 119) = 7.229 |
| Genotype x Treatmen t | 0.2648 | F (2, 119) = 1.344 |

| Uncorrected Fisher's LSD | Predicted (LS) mean diff. | 95.00% CI of diff. | Individual P Value |
|---|---|---|---|
| WT:Pre vs. WT:Veh | -0.3078 | -0.4589 to -0.156 | <0.0001 |
| WT:Pre vs. WT:SNAP | -0.1106 | -0.292 to 0.0712 | 0.2309 |
| WT:Pre vs. KO:Pre | -0.06886 | -0.193 to 0.0556 | 0.2758 |
| WT:Pre vs. KO:Veh | -0.1988 | -0.3644 to -0.033 | 0.0190 |
| WT:Pre vs. KO:SNAP | -0.04363 | -0.278 to 0.1909 | 0.7132 |
| WT:Veh vs. WT:SNAP | 0.1972 | -0.008 to 0.4032 | 0.0604 |
| WT:Veh vs. KO:Pre | 0.2389 | 0.08123 to 0.396 | 0.0033 |
| WT:Veh vs. KO:Veh | 0.1090 | -0.0828 to 0.300 | 0.2628 |
| WT:Veh vs. KO:SNAP | 0.2641 | 0.0104 to 0.5178 | 0.0414 |
| WT:SNAP vs. KO:Pre | 0.04173 | -0.1456 to 0.229 | 0.6600 |
| WT:SNAP vs. KO:Veh | -0.08821 | -0.305 to 0.1286 | 0.4221 |
| WT:SNAP vs. KO:SNAP | 0.06696 | -0.206 to 0.3401 | 0.6283 |
| KO:Pre vs. KO:Veh | -0.1299 | -0.3016 to 0.042 | 0.1365 |

| | | | |
|---|---|---|---|
| KO:Pre vs. KO:SNAP | 0.02523 | -0.213 to 0.2641 | 0.8347 |
| KO:Veh vs. KO:SNAP | 0.1552 | -0.107 to 0.4178 | 0.2443 |

Resting EEG Power Coupling D2G1

| ANOVA results | P value | F (DFn, DFd) |
|---|---|---|
| Genotype | 0.6608 | F (1, 119) = 0.1936 |
| Treatment | 0.0041 | F (2, 119) = 5.746 |
| Genotype x Treatment | 0.4438 | F (2, 119) = 0.8180 |

| **Uncorrected Fisher's LSD** | **Predicted (LS) mean diff.** | **95.00% CI of diff.** | **Individual P Value** |
|---|---|---|---|
| WT:Pre vs. WT:Veh | -0.2209 | -0.353 to -0.0884 | 0.0013 |
| WT:Pre vs. WT:SNAP | -0.1066 | -0.266 to 0.0527 | 0.1877 |
| WT:Pre vs. KO:Pre | -0.1015 | -0.2106 to 0.007 | 0.0682 |
| WT:Pre vs. KO:Veh | -0.2232 | -0.368 to -0.0780 | 0.0029 |
| WT:Pre vs. KO:SNAP | -0.07224 | -0.2778 to 0.133 | 0.4878 |
| WT:Veh vs. WT:SNAP | 0.1142 | -0.066 to 0.2947 | 0.2127 |
| WT:Veh vs. KO:Pre | 0.1194 | -0.0188 to 0.257 | 0.0897 |
| WT:Veh vs. KO:Veh | -0.002306 | -0.170 to 0.1658 | 0.9784 |
| WT:Veh vs. KO:SNAP | 0.1486 | -0.0737 to 0.371 | 0.1882 |
| WT:SNAP vs. KO:Pre | 0.005180 | -0.159 to 0.1694 | 0.9503 |
| WT:SNAP vs. KO:Veh | -0.1165 | -0.3066 to 0.073 | 0.2271 |
| WT:SNAP vs. KO:SNAP | 0.03440 | -0.205 to 0.2738 | 0.7765 |
| KO:Pre vs. KO:Veh | -0.1217 | -0.272 to 0.0287 | 0.1118 |
| KO:Pre vs. KO:SNAP | 0.02922 | -0.180 to 0.2385 | 0.7828 |
| KO:Veh vs. KO:SNAP | 0.1509 | -0.0792 to 0.381 | 0.1966 |

Resting EEG Power Coupling D2G2

| ANOVA results | P value | F (DFn, DFd) |
|---|---|---|
| Genotype | 0.6384 | F (1, 119) = 0.2220 |
| Treatment | 0.0013 | F (2, 119) = 7.005 |
| Genotype x Treatment | 0.4294 | F (2, 119) = 0.8515 |

| Uncorrected Fisher's LSD | Predicted (LS) mean diff. | 95.00% CI of diff. | Individual P Value |
|---|---|---|---|
| WT:Pre vs. WT:Veh | -0.2602 | -0.4018 to -0.118 | 0.0004 |
| WT:Pre vs. WT:SNAP | -0.07270 | -0.243 to 0.0976 | 0.3998 |
| WT:Pre vs. KO:Pre | -0.1093 | -0.226 to 0.0074 | 0.0662 |
| WT:Pre vs. KO:Veh | -0.2467 | -0.402 to -0.092 | 0.0021 |
| WT:Pre vs. KO:SNAP | -0.05637 | -0.276 to 0.163 | 0.6125 |
| WT:Veh vs. WT:SNAP | 0.1875 | -0.005 to 0.3805 | 0.0567 |
| WT:Veh vs. KO:Pre | 0.1510 | 0.0032 to 0.2987 | 0.0453 |
| WT:Veh vs. KO:Veh | 0.01352 | -0.166 to 0.1932 | 0.8818 |
| WT:Veh vs. KO:SNAP | 0.2039 | -0.0338 to 0.442 | 0.0921 |
| WT:SNAP vs. KO:Pre | -0.03657 | -0.212 to 0.1390 | 0.6808 |
| WT:SNAP vs. KO:Veh | -0.1740 | -0.377 to 0.0292 | 0.0925 |
| WT:SNAP vs. KO:SNAP | 0.01634 | -0.2396 to 0.272 | 0.8996 |
| KO:Pre vs. KO:Veh | -0.1375 | -0.298 to 0.0234 | 0.0932 |
| KO:Pre vs. KO:SNAP | 0.05290 | -0.171 to 0.2767 | 0.6406 |
| KO:Veh vs. KO:SNAP | 0.1904 | -0.0557 to 0.436 | 0.1282 |

**Table S7**

Statistics for Figure 8B

Locomotor activity in whole arena

| ANOVA table | SS (Type III) | DF | MS | F (DFn, DFd) | P value | R squared |
|---|---|---|---|---|---|---|
| Group | 176.7 | 3 | 58.90 | F (3, 93) = 2.428 | P=0.0703 | 0.07262 |

| Tukey's multiple comparisons test | Mean Diff. | 95.00% CI of diff. | Adjusted P Value |
|---|---|---|---|
| Ctrl WT vs. Het | 0.6681 | -2.517 to 3.854 | 0.9467 |
| Ctrl WT vs. cKO | -2.883 | -6.550 to 0.7830 | 0.1749 |
| Ctrl WT vs. cKO SNAP | -0.8083 | -5.514 to 3.897 | 0.9696 |
| Het vs. cKO | -3.551 | -7.090 to -0.01315 | 0.0488 |
| Het vs. cKO SNAP | -1.476 | -6.083 to 3.130 | 0.8360 |
| cKO vs. cKO SNAP | 2.075 | -2.876 to 7.026 | 0.6926 |

Statistics for Figure 8C

Locomotor activity in thigmotaxis

| ANOVA table | SS (Type III) | DF | MS | F (DFn, DFd) | P value | R squared |
|---|---|---|---|---|---|---|
| Genotype | 228.0 | 3 | 76.00 | F (3, 93) = 3.293 | P=0.0240 | 0.09602 |

| Tukey's multiple comparisons test | Mean Diff. | 95.00% CI of diff. | Adjusted P Value |
|---|---|---|---|
| Ctrl WT vs. Het | 0.1497 | -2.957 to 3.257 | 0.9993 |
| Ctrl WT vs. cKO | -3.670 | -7.245 to -0.09356 | 0.0420 |
| Ctrl WT vs. cKO SNAP | -0.2017 | -4.791 to 4.388 | 0.9995 |
| Het vs. cKO | -3.819 | -7.270 to -0.3682 | 0.0240 |
| Het vs. cKO SNAP | -0.3514 | -4.844 to 4.141 | 0.9969 |
| cKO vs. cKO SNAP | 3.468 | -1.361 to 8.297 | 0.2442 |

Statistics for Figure 8D
Exploratory behavior

| ANOVA table | SS (Type III) | DF | MS | F (DFn, DFd) | P value | % of total variation |
|---|---|---|---|---|---|---|
| Preference x Genotype | 7030 | 3 | 2343 | F (3, 106) = 7.917 | P<0.0001 | 16.66 |
| Preference | 7573 | 1 | 7573 | F (1, 106) = 25.58 | P<0.0001 | 17.94 |
| Genotype | 237.9 | 3 | 79.30 | F (3, 106) = 43.80 | P<0.0001 | 0.5638 |

| Tukey's multiple comparisons test | Predicted (LS) mean diff. | 95.00% CI of diff. | Adjusted P Value |
|---|---|---|---|
| Ctrl WT | | | |
| Open field vs. Thigmotaxis | -10.16 | -18.96 to -1.350 | 0.0242 |
| Het | | | |
| Open field vs. Thigmotaxis | 1.300 | -7.983 to 10.58 | 0.7818 |
| cKO | | | |
| Open field vs. Thigmotaxis | -4.186 | -11.54 to 3.170 | 0.2618 |
| cKO SNAP | | | |
| Open field vs. Thigmotaxis | -41.11 | -56.36 to -25.86 | <0.0001 |
| Thigmotaxis | | | |
| Het vs. Ctrl WT | -5.740 | -14.12 to 2.642 | 0.2891 |
| cKO vs. Ctrl WT | -2.782 | -10.30 to 4.735 | 0.7732 |
| cKO SNAP vs. Ctrl WT | 11.96 | 0.4220 to 23.50 | 0.0390 |
| cKO vs. Het | 2.958 | -4.801 to 10.72 | 0.7568 |
| cKO SNAP vs. Het | 17.70 | 6.003 to 29.40 | 0.0007 |
| cKO SNAP vs. cKO | 14.74 | 3.648 to 25.84 | 0.0039 |
| Open field | | | |
| Het vs. Ctrl WT | 5.717 | -2.666 to 14.10 | 0.2926 |
| cKO vs. Ctrl WT | 3.189 | -4.328 to 10.71 | 0.6908 |
| cKO SNAP vs. Ctrl WT | -18.99 | -30.53 to -7.455 | 0.0002 |

| | | | |
|---|---|---|---|
| cKO vs. Het | -2.528 | -10.29 to 5.231 | 0.8335 |
| cKO SNAP vs. Het | -24.71 | -36.41 to -13.01 | <0.0001 |
| cKO SNAP vs. cKO | -22.18 | -33.28 to -11.09 | <0.0001 |

Statistics for Figure 8E
Anxiety-like behavior

| ANOVA table | SS (Type III) | DF | MS | F (DFn, DFd) | P value | R squared |
|---|---|---|---|---|---|---|
| Group | 24.93 | 3 | 8.309 | F (3, 102) = 0.1931 | P=0.9009 | 0.005649 |

| Tukey's multiple comparisons test | Mean Diff. | 95.00% CI of diff. | Adjusted P Value |
|---|---|---|---|
| Ctrl WT vs. Het | -0.5178 | -4.963 to 3.928 | 0.9902 |
| Ctrl WT vs. cKO | 0.6449 | -3.716 to 5.006 | 0.9803 |
| Ctrl WT vs. cKO SNAP | 0.4702 | -5.872 to 6.812 | 0.9974 |
| Het vs. cKO | 1.163 | -2.966 to 5.291 | 0.8825 |
| Het vs. cKO SNAP | 0.9880 | -5.196 to 7.172 | 0.9754 |
| cKO vs. cKO SNAP | -0.1747 | -6.298 to 5.949 | 0.9999 |

**Table S8**
Statistics for Figure S1

| GAT3 mRNA in AuC | Mean | SEM | N |
|---|---|---|---|
| Ctrl WT | 1.014 | 0.08360 | 5 |
| cKO | 0.9697 | 0.04553 | 4 |
| Statistics | Two-tailed, unpaired t-test | t=0.4309, df=7, p=0.6795 $\eta^2$=0.02584 | |

| GAT3 mRNA in FC | Mean | SEM | N |
|---|---|---|---|
| Ctrl WT | 1.013 | 0.07932 | 5 |
| cKO | 0.9782 | 0.02823 | 4 |
| Statistics | Two-tailed, unpaired t-test | t=0.3762, df=7, p=0.7179 $\eta^2$=0.01982 | |

**Table S9**
Statistics for Figure S2A

| Gabrg2 mRNA in AuC | Mean | SEM | N |
|---|---|---|---|
| Ctrl WT | 1.110 | 0.04341 | 5 |
| cKO | 0.8950 | 0.05685 | 4 |
| Statistics | Two-tailed, unpaired t-test | t=3.062, df=7, p= 0.0183 $\eta^2$= 0.5726 | |

| Gabrg2 mRNA in FC | Mean | SEM | N |
|---|---|---|---|
| Ctrl WT | 1.034 | 0.01727 | 4 |
| cKO | 1.225 | 0.1413 | 4 |

| | | |
|---|---|---|
| Statistics | Two-tailed, unpaired t-test | $t=1.344$, $df=6$, $p= 0.2275$ $\eta^2= 0.2315$ |

Statistics for Figure S2B

| Gabra1 mRNA in AuC | **Mean** | **SEM** | **N** |
|---|---|---|---|
| Ctrl WT | 1.009 | 0.06766 | 5 |
| cKO | 0.7900 | 0.05683 | 4 |
| Statistics | Two-tailed, unpaired t-test | $t=2.389$, $df=7$, $p= 0.0483$ $\eta^2= 0.4491$ | |

| Gabra1 mRNA in FC | **Mean** | **SEM** | **N** |
|---|---|---|---|
| Ctrl WT | 1.009 | 0.06929 | 5 |
| cKO | 1.128 | 0.1446 | 4 |
| Statistics | Two-tailed, unpaired t-test | $t=0.7962$, $df=7$, $p= 0.4521$ $\eta^2= 0.08305$ | |

Statistics for Figure S2C

| Gabra3 mRNA in AuC | **Mean** | **SEM** | **N** |
|---|---|---|---|
| Ctrl WT | 1.005 | 0.05176 | 5 |
| cKO | 1.020 | 0.1034 | 4 |
| Statistics | Two-tailed, unpaired t-test | $t=0.1403$, $df=7$, $p=0.8923$ $\eta^2=0.002806$ | |

| Gabra3 mRNA in FC | **Mean** | **SEM** | **N** |
|---|---|---|---|
| Ctrl WT | 1.006 | 0.05811 | 5 |
| cKO | 0.9004 | 0.09001 | 4 |
| Statistics | Two-tailed, unpaired t-test | $t=1.030$, $df=7$, $p= 0.3371$ $\eta^2= 0.1317$ | |

Statistics for Figure S2D

| Gabra5 mRNA in AuC | **Mean** | **SEM** | **N** |
|---|---|---|---|
| Ctrl WT | 1.024 | 0.1040 | 5 |
| cKO | 1.262 | 0.1712 | 4 |
| Statistics | Two-tailed, unpaired t-test | $t=1.248$, $df=7$, $p= 0.2523$ $\eta^2= 0.1819$ | |

| Gabra5 mRNA in FC | **Mean** | **SEM** | **N** |
|---|---|---|---|
| Ctrl WT | 1.004 | 0.04863 | 5 |
| cKO | 0.9706 | 0.1177 | 4 |
| Statistics | Two-tailed, unpaired t-test | $t=0.2885$, $df=7$, $p= 0.7813$ $\eta^2= 0.01175$ | |

Statistics for Figure S2E

| GABAARγ2 mRNA in AuC | Mean | SEM | N |
|---|---|---|---|
| Ctrl WT | 1.039 | 0.04794 | 4 |
| cKO | 1.073 | 0.06912 | 5 |
| Statistics | Two-tailed, unpaired t-test | t=0.3914, df=7, p= 0.7071 $\eta^2$= 0.02142 | |

| GABAARγ2 mRNA in FC | Mean | SEM | N |
|---|---|---|---|
| Ctrl WT | 0.9726 | 0.03286 | 5 |
| cKO | 0.9810 | 0.09556 | 4 |
| Statistics | Two-tailed, unpaired t-test | t=0.07529, df=7, p= 0.9421 $\eta^2$= 0.0008092 | |

Statistics for Figure S2F

| GABAARα5 mRNA in AuC | Mean | SEM | N |
|---|---|---|---|
| Ctrl WT | 0.9998 | 0.09649 | 4 |
| cKO | 1.226 | 0.1014 | 4 |
| Statistics | Two-tailed, unpaired t-test | t=1.616, df=6, p= 0.1571 $\eta^2$= 0.3033 | |

| GABAARα5 mRNA in FC | Mean | SEM | N |
|---|---|---|---|
| Ctrl WT | 0.9245 | 0.1485 | 4 |
| cKO | 0.9165 | 0.2269 | 4 |
| Statistics | Two-tailed, unpaired t-test | t=0.02950, df=6, p= 0.9774 $\eta^2$= 0.0001450 | |

**Table S10**

Statistics for Figure S3A

PV levels in L1-4

| ANOVA table | SS (Type III) | DF | MS | F (DFn, DFd) | P value | % of total variation |
|---|---|---|---|---|---|---|
| Genotype x Treatment | 4446 | 1 | 4446 | F (1, 52) = 6.160 | P=0.0163 | 8.417 |
| Genotype | 2317 | 1 | 2317 | F (1, 52) = 3.211 | P=0.0790 | 4.388 |
| Treatment | 10180 | 1 | 10180 | F (1, 52) = 14.11 | P=0.0004 | 19.28 |

| Uncorrected Fisher's LSD | Predicted (LS) mean diff. | 95.00% CI of diff. | Individual P Value |
|---|---|---|---|
| Ctrl WT | | | |
| Vehicle vs. SNAP | -9.241 | -28.30 to 9.819 | 0.3351 |
| cKO | | | |
| Vehicle vs. SNAP | -45.25 | -67.26 to -23.24 | 0.0001 |
| Vehicle | | | |
| Ctrl WT vs. cKO | 31.00 | 10.42 to 51.59 | 0.0039 |
| SNAP | | | |
| Ctrl WT vs. cKO | -5.006 | -25.59 to 15.58 | 0.6277 |

PV levels in L5-6

| ANOVA table | SS (Type III) | DF | MS | F (DFn, DFd) | P value | % of total variation |
|---|---|---|---|---|---|---|
| Genotype x Treatment | 2386 | 1 | 2386 | F (1, 52) = 1.988 | P=0.1645 | 2.808 |
| Genotype | 1649 | 1 | 1649 | F (1, 52) = 1.375 | P=0.2464 | 1.942 |
| Treatment | 20073 | 1 | 20073 | F (1, 52) = 16.73 | P=0.0002 | 23.63 |

| Uncorrected Fisher's LSD | Predicted (LS) mean diff. | 95.00% CI of diff. | Individual P Value |
|---|---|---|---|
| Ctrl WT | | | |
| Vehicle vs. SNAP | -25.07 | -49.64 to -0.4948 | 0.0457 |
| cKO | | | |
| Vehicle vs. SNAP | -51.45 | -79.82 to -23.07 | 0.0006 |
| Vehicle | | | |
| Ctrl WT vs. cKO | 24.16 | -2.388 to 50.70 | 0.0736 |
| SNAP | | | |
| Ctrl WT vs. cKO | -2.222 | -28.77 to 24.32 | 0.8672 |

Statistics for Figure S3B

Proportion of low PV and high PV cells

| ANOVA table | SS (Type III) | DF | MS | F (DFn, DFd) | P value | % of total variation |
|---|---|---|---|---|---|---|
| PV expression x Genotype/ Treatment | 62023 | 3 | 20674 | F (3, 100) = 18.45 | P<0.0001 | 28.81 |
| PV expression | 33632 | 1 | 33632 | F (1, 100) = 30.01 | P<0.0001 | 15.62 |

| | | | | | | |
|---|---|---|---|---|---|---|
| Genotype/ Treatment | 5.831e-005 | 3 | 1.944e-005 | F (3, 100) = 1.735e-008 | P>0.9999 | 2.709e-008 |

| Šídák's multiple comparisons test | Predicted (LS) mean diff. | 95.00% CI of diff. | Adjusted P Value |
|---|---|---|---|
| low PV | | | |
| Ctrl WT vs. cKO | -40.67 | -76.34 to -5.004 | 0.0169 |
| Ctrl WT vs. Ctrl WT SNAP | 12.99 | -19.30 to 45.28 | 0.8640 |
| Ctrl WT vs. cKO SNAP | 29.37 | -5.424 to 64.17 | 0.1442 |
| cKO vs. Ctrl WT SNAP | 53.66 | 18.47 to 88.85 | 0.0005 |
| cKO vs. cKO SNAP | 70.05 | 32.54 to 107.6 | <0.0001 |
| Ctrl WT SNAP vs. cKO SNAP | 16.39 | -17.93 to 50.70 | 0.7434 |
| high PV | | | |
| Ctrl WT vs. cKO | 40.67 | 5.005 to 76.34 | 0.0169 |
| Ctrl WT vs. Ctrl WT SNAP | -12.99 | -45.28 to 19.30 | 0.8641 |
| Ctrl WT vs. cKO SNAP | -29.37 | -64.17 to 5.428 | 0.1443 |
| cKO vs. Ctrl WT SNAP | -53.66 | -88.85 to -18.47 | 0.0005 |
| cKO vs. cKO SNAP | -70.04 | -107.5 to -32.54 | <0.0001 |
| Ctrl WT SNAP vs. cKO SNAP | -16.38 | -50.70 to 17.93 | 0.7436 |
| Ctrl WT | | | |
| low PV vs. high PV | -34.87 | -59.12 to -10.62 | 0.0053 |
| cKO | | | |
| low PV vs. high PV | 46.47 | 18.15 to 74.79 | 0.0015 |
| Ctrl WT SNAP | | | |
| low PV vs. high PV | -60.85 | -84.33 to -37.37 | <0.0001 |
| cKO SNAP | | | |
| low PV vs. high PV | -93.62 | -120.7 to -66.50 | <0.0001 |

**Table S11**

Statistics for Figure S5A-S5L

| Frequency | Factor | ANOVA results | p-value |
|---|---|---|---|
| **AuC** | | | |
| P1 amplitude | Genotype | "F (1, 20) = 0.03025" | 0.8637 |
| | Treatment | "F (1, 17) = 0.6828" | 0.4201 |
| | Genotype x Treatment | "F (1, 17) = 0.2966" | 0.5931 |
| P1 latency | Genotype | "F (1, 20) = 0.2562" | 0.6183 |
| | Treatment | "F (1, 17) = 0.6882" | 0.4183 |
| | Genotype x Treatment | "F (1, 17) = 1.163" | 0.2959 |

| | | | |
|---|---|---|---|
| N1 amplitude | Genotype | "F (1, 20) = 0.06962" | 0.7946 |
| | Treatment | "F (1, 17) = 0.3064" | 0.5871 |
| | Genotype x Treatment | "F (1, 17) = 0.01605" | 0.9007 |
| N1 latency | Genotype | "F (1, 20) = 0.2261" | 0.6396 |
| | Treatment | "F (1, 17) = 0.003827" | 0.9514 |
| | Genotype x Treatment | "F (1, 17) = 0.08865" | 0.7695 |
| P2 amplitude | Genotype | "F (1, 20) = 0.2682" | 0.6102 |
| | Treatment | "F (1, 17) = 0.004136" | 0.9495 |
| | Genotype x Treatment | "F (1, 17) = 0.01481" | 0.9046 |
| P2 latency | Genotype | "F (1, 20) = 0.7012" | 0.4123 |
| | Treatment | "F (1, 17) = 5.268" | 0.0347 |
| | Genotype x Treatment | "F (1, 17) = 0.002633" | 0.9597 |
| **FC** | | | |
| P1 amplitude | Genotype | "F (1, 20) = 1.378" | 0.2543 |
| | Treatment | "F (1, 17) = 1.895" | 0.1865 |
| | Genotype x Treatment | "F (1, 17) = 1.445" | 0.2457 |
| P1 latency | Genotype | "F (1, 36) = 0.009201" | 0.9241 |
| | Treatment | "F (1, 36) = 4.259" | 0.0463 |
| | Genotype x Treatment | "F (1, 36) = 0.1509" | 0.6999 |
| N1 amplitude | Genotype | "F (1, 20) = 0.04591" | 0.8325 |
| | Treatment | "F (1, 17) = 0.3190" | 0.5796 |
| | Genotype x Treatment | "F (1, 17) = 1.129" | 0.3028 |
| N1 latency | Genotype | "F (1, 20) = 0.01005" | 0.9211 |
| | Treatment | "F (1, 17) = 0.005249" | 0.9431 |
| | Genotype x Treatment | "F (1, 17) = 1.326" | 0.2654 |
| P2 amplitude | Genotype | "F (1, 20) = 0.2456" | 0.6256 |
| | Treatment | "F (1, 17) = 0.6518" | 0.4306 |
| | Genotype x Treatment | "F (1, 17) = 0.1529" | 0.7007 |
| P2 latency | Genotype | "F (1, 20) = 0.06216" | 0.8057 |
| | Treatment | "F (1, 17) = 0.7778" | 0.3901 |
| | Genotype x Treatment | "F (1, 17) = 0.1582" | 0.6958 |

Western Blots

FMRP blot for Figure 1D

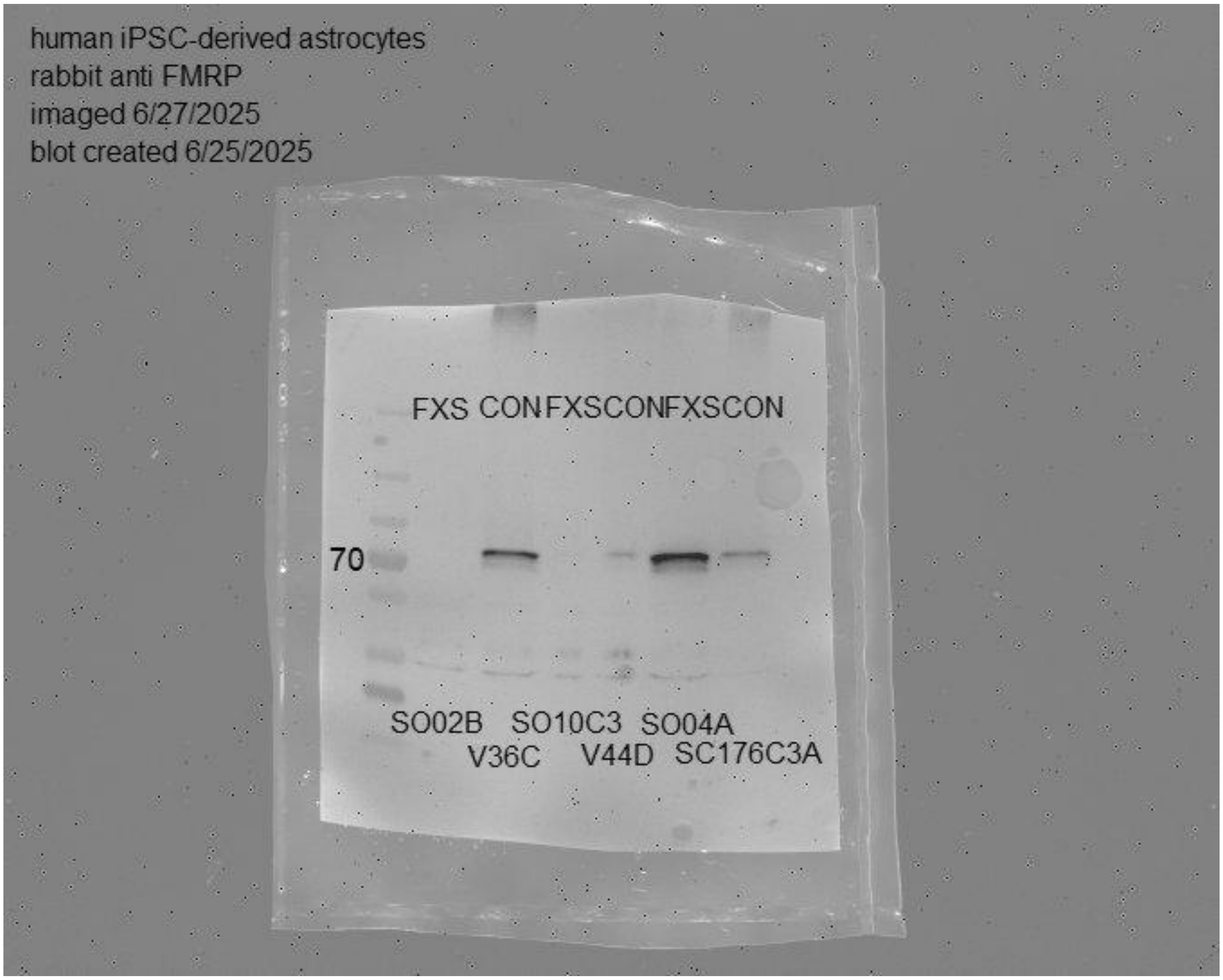

Beta-actin blot for Figure 1D, 1I

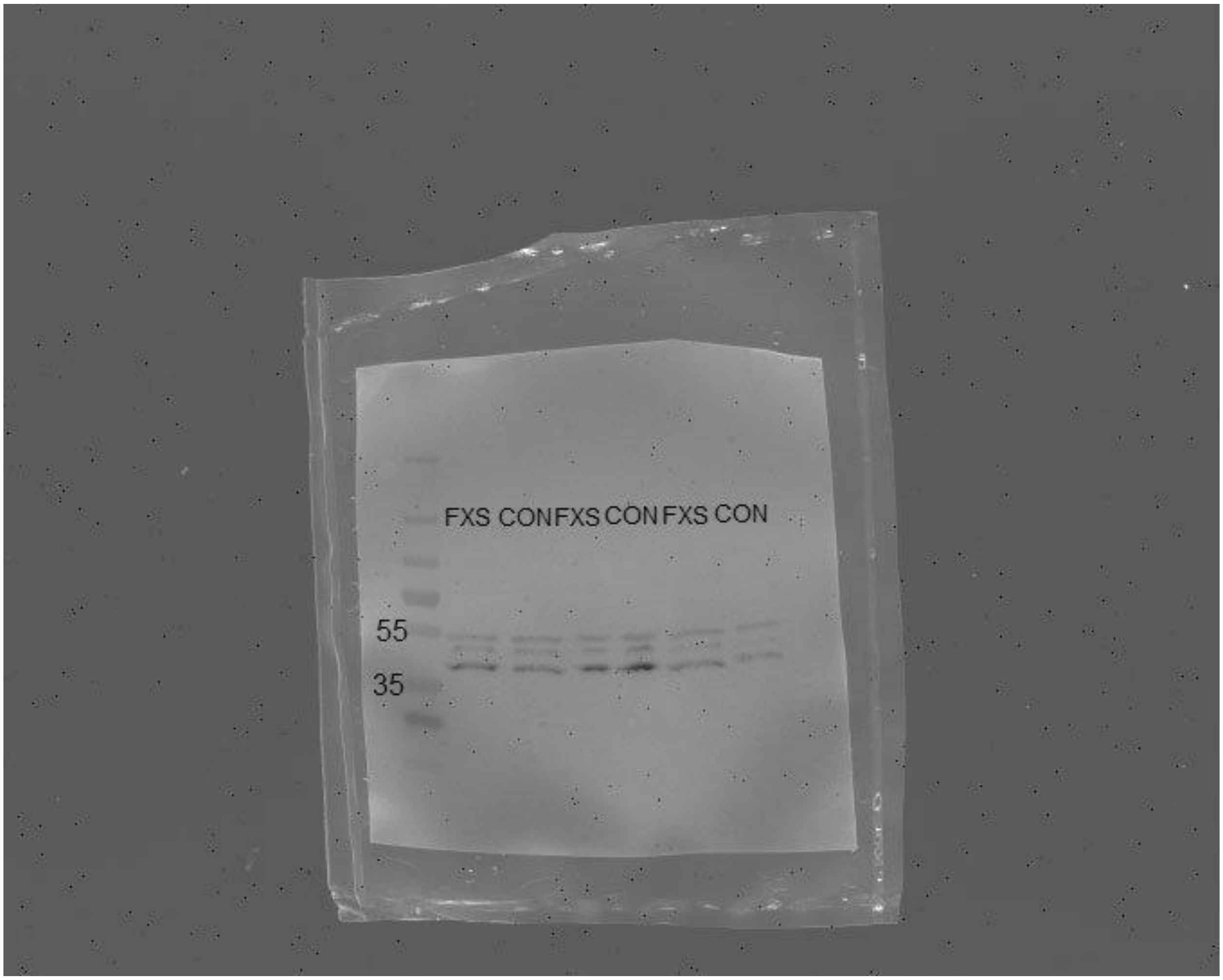

GAD65/67 blots for Figure 1I

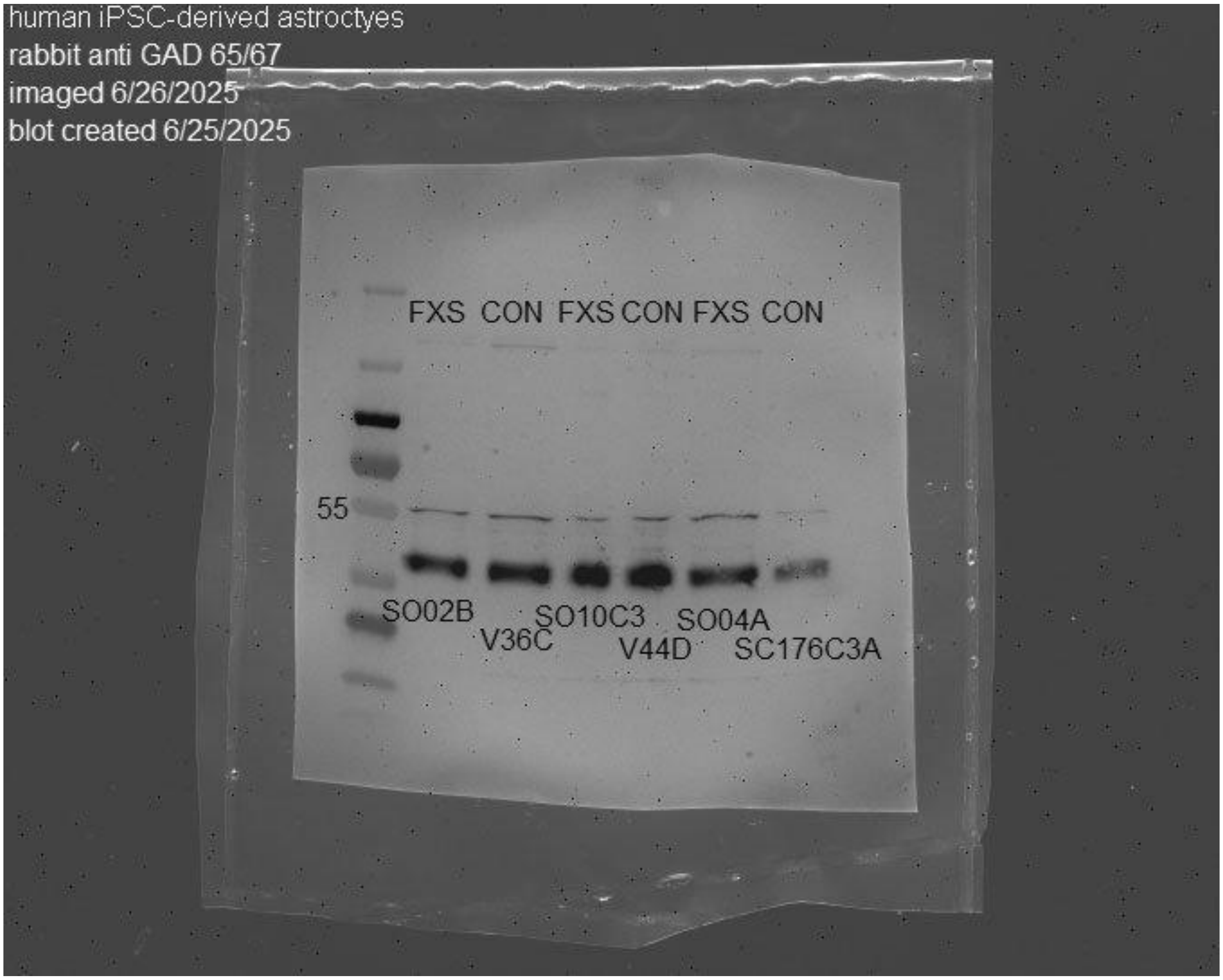

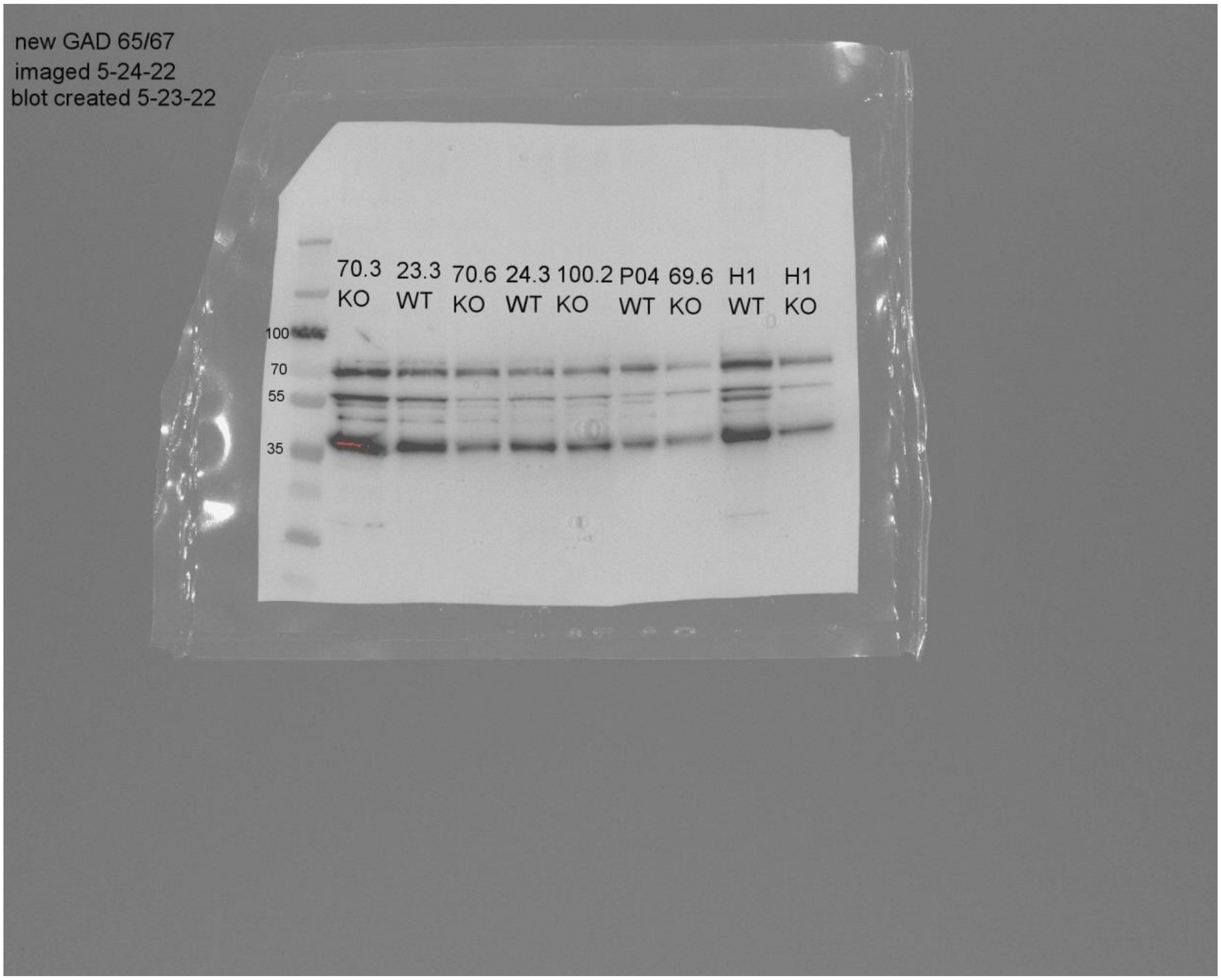
new GAD 65/67
imaged 5-24-22
blot created 5-23-22
70.3 KO
23.3 WT
70.6 KO
24.3 WT
100.2 KO
P04 WT
69.6 KO
H1 WT
H1 KO
100
70
55
35

Beta-actin blot for Figure 1I

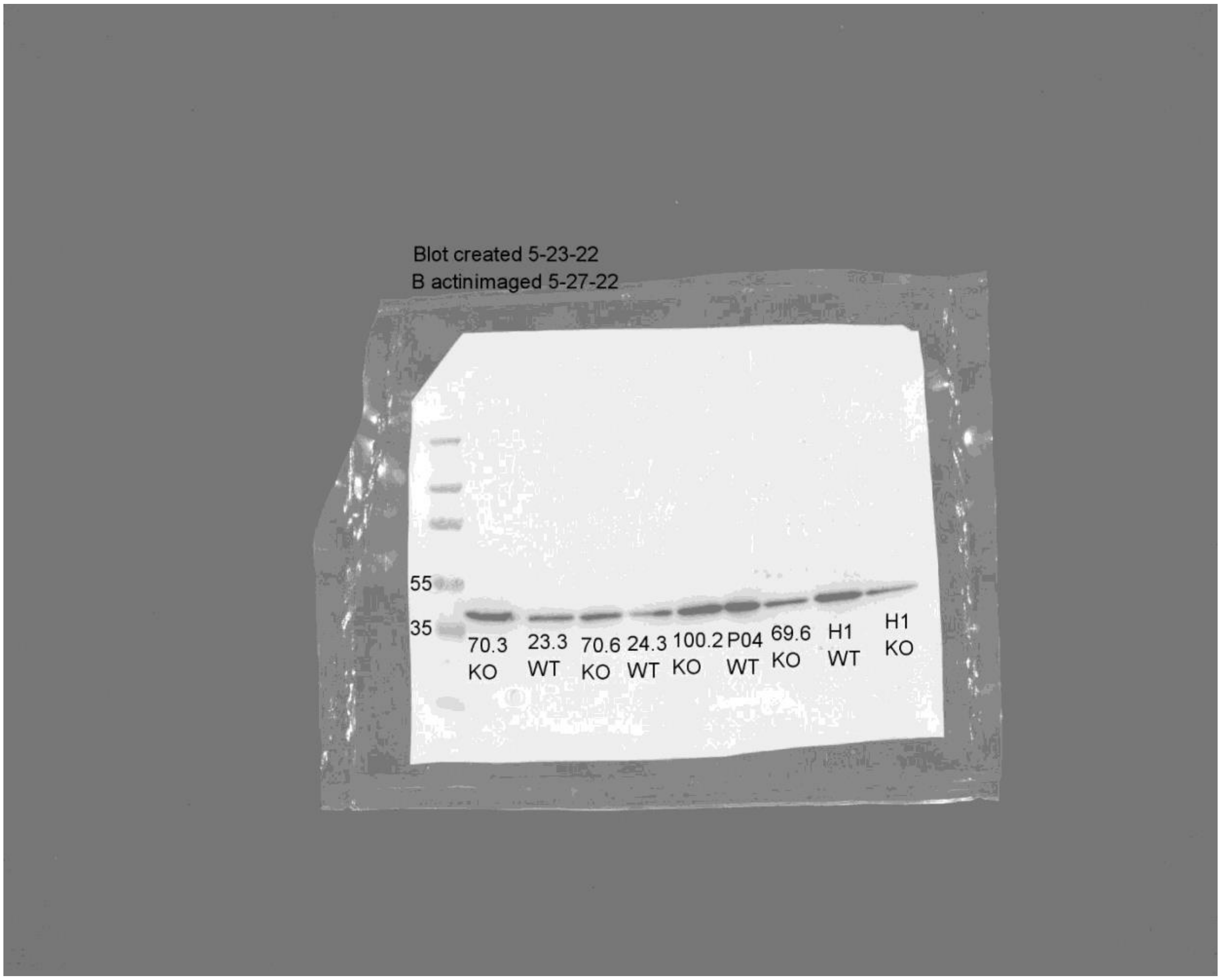

GAD 65/67 blots for Figure 2B

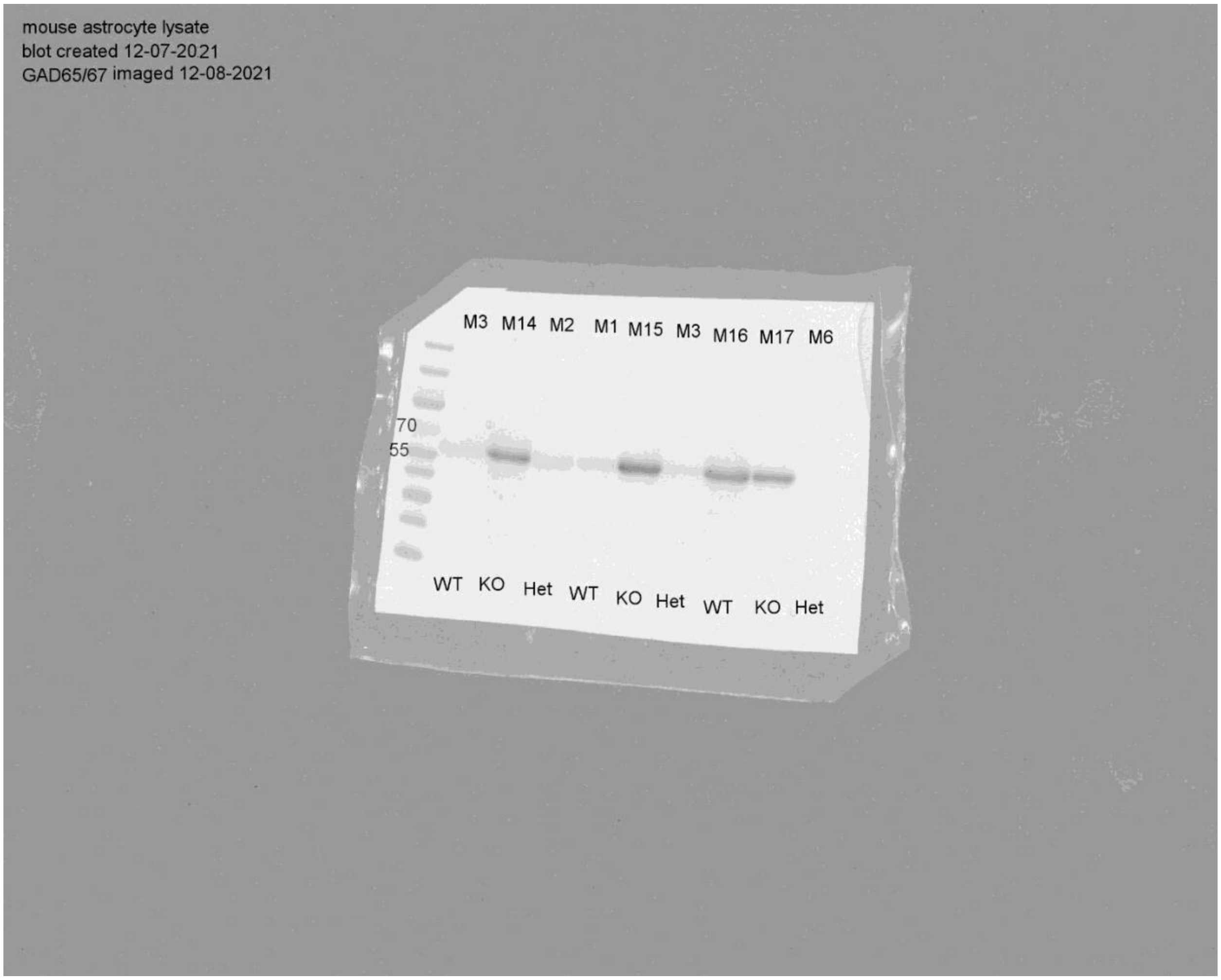

Beta-actin blots for Figure 2B

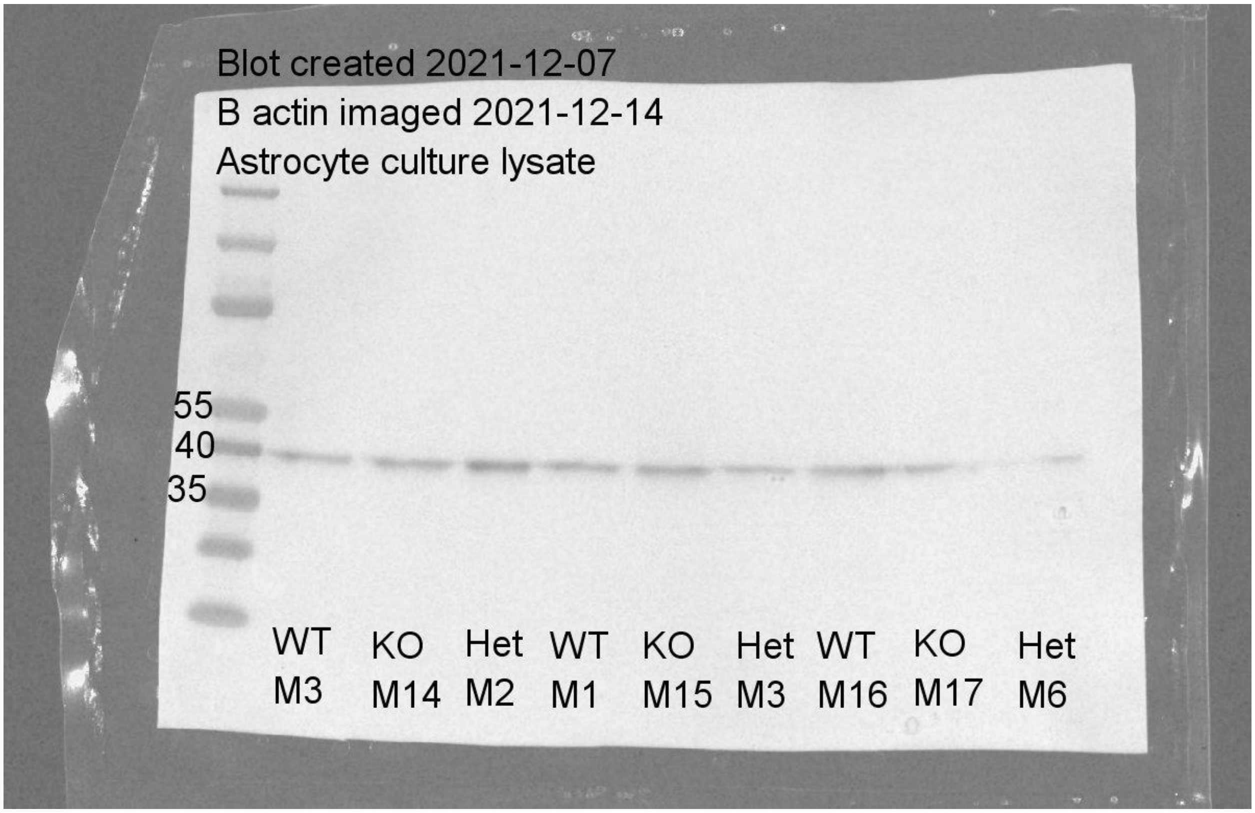

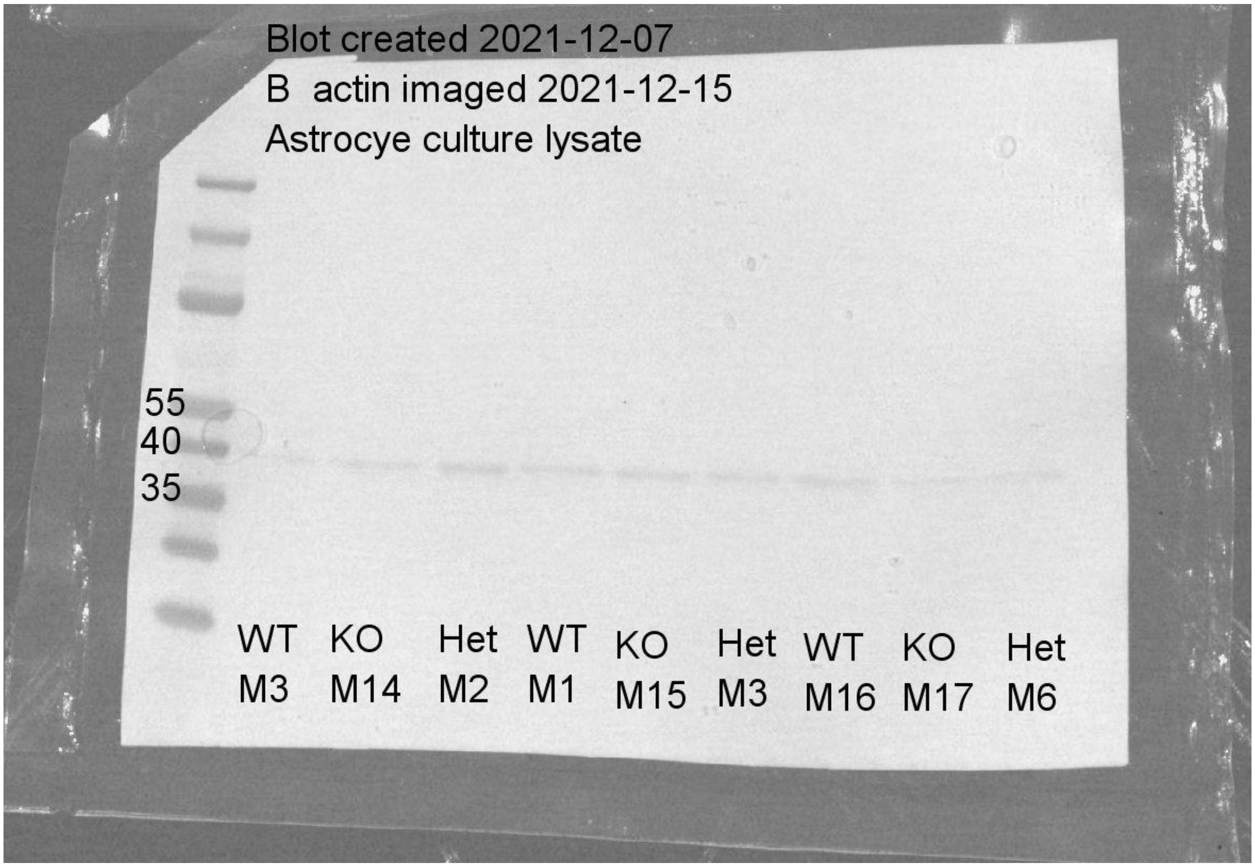
Blot created 2021-12-07
B actin imaged 2021-12-15
Astrocye culture lysate
55
40
35
WT M3
KO M14
Het M2
WT M1
KO M15
Het M3
WT M16
KO M17
Het M6

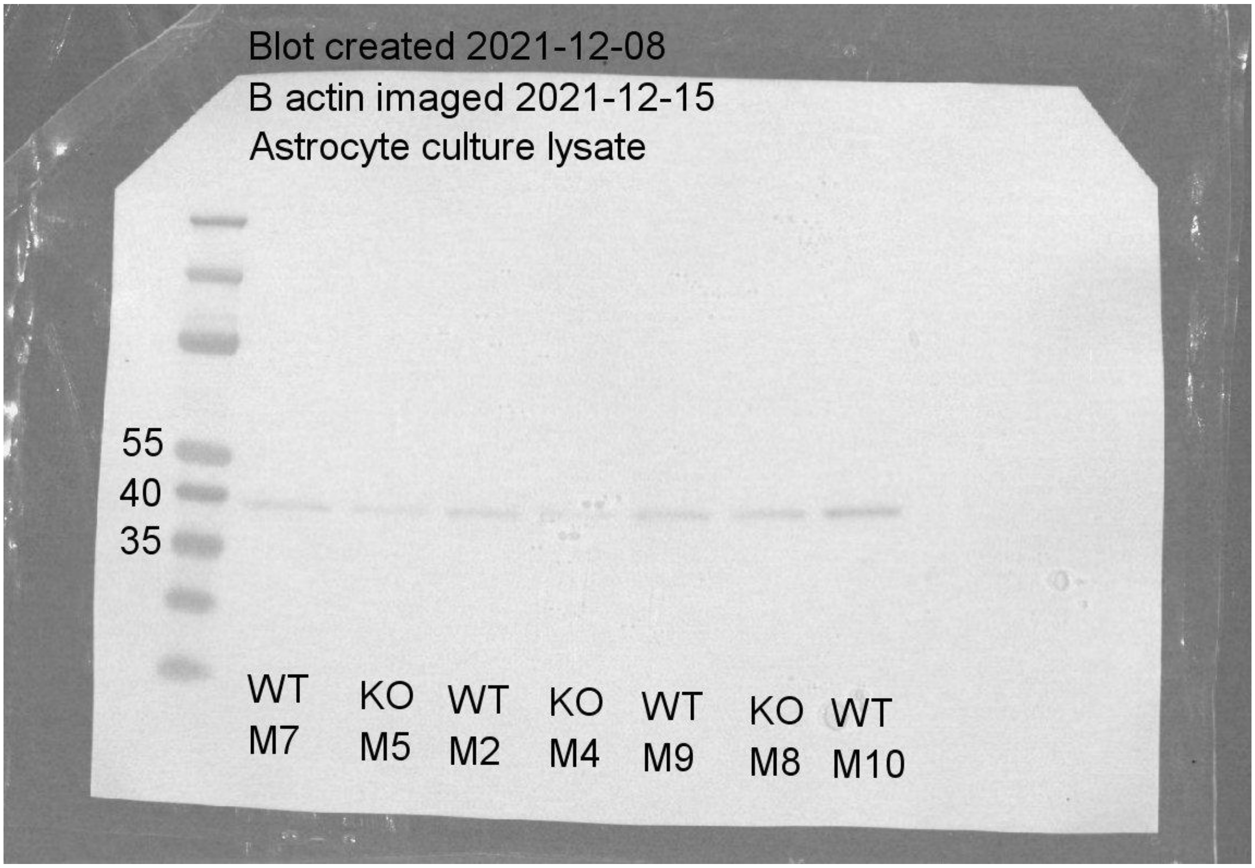
Blot created 2021-12-08
B actin imaged 2021-12-15
Astrocyte culture lysate
55
40
35
WT M7
KO M5
WT M2
KO M4
WT M9
KO M8
WT M10

Aldha1 blots for Figure 2B

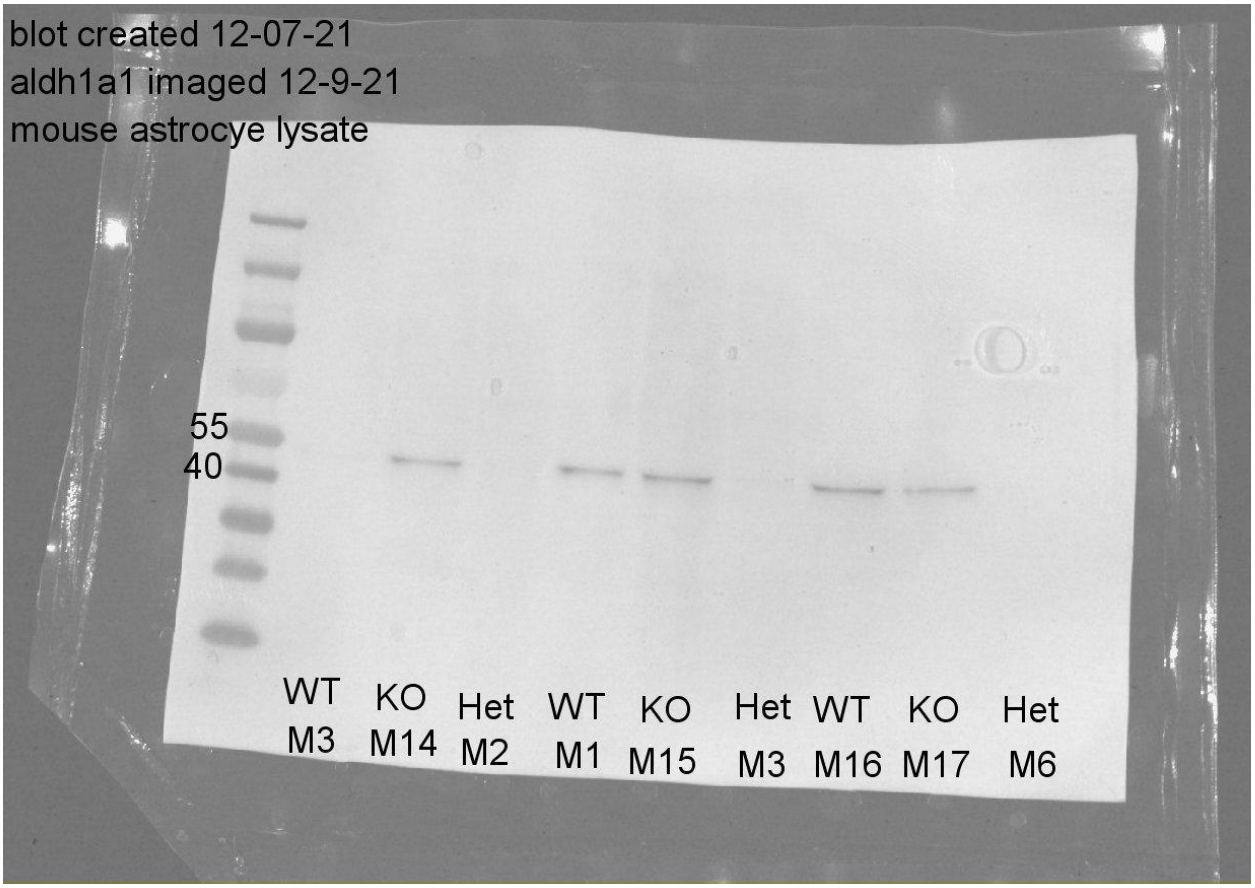

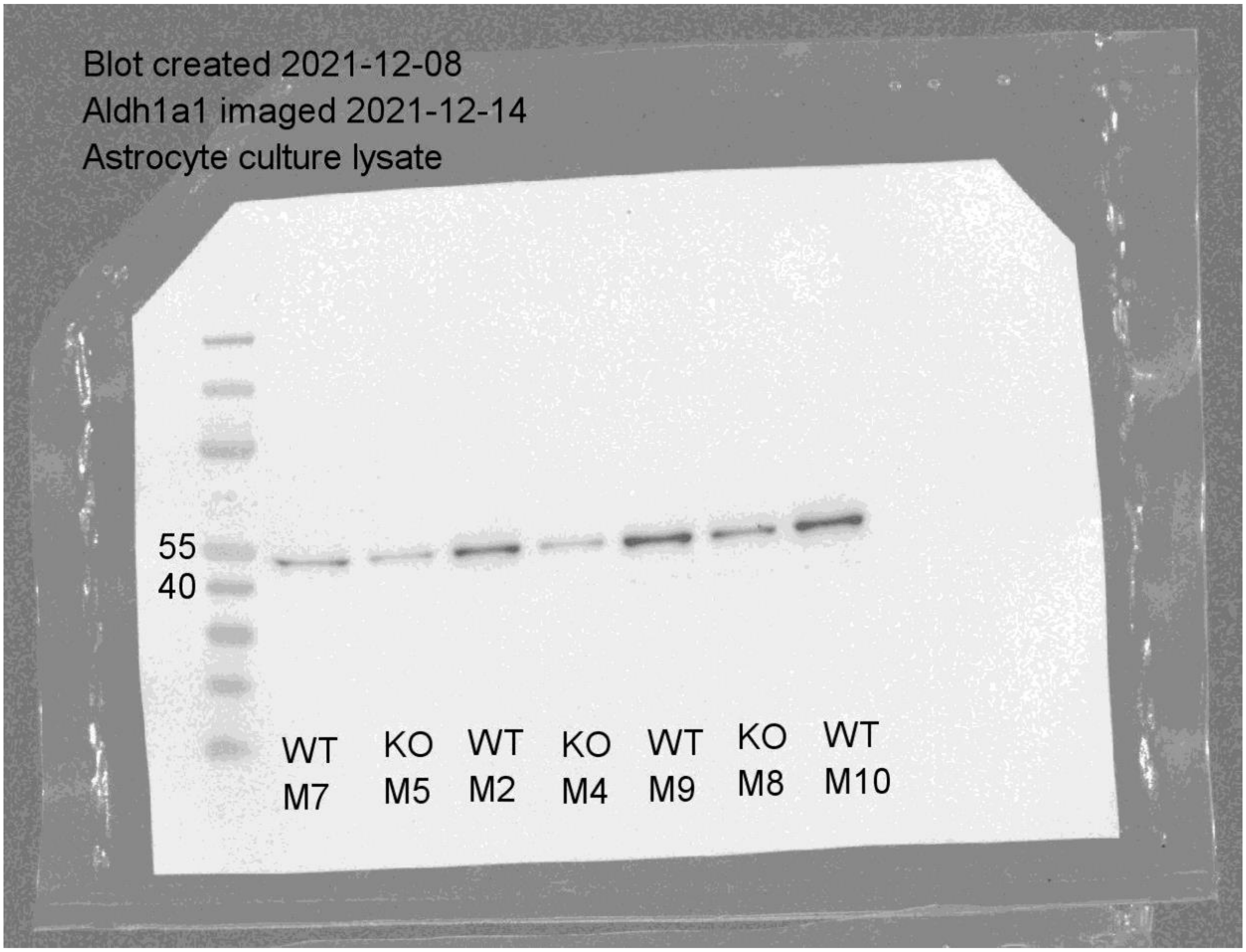
Blot created 2021-12-08
Aldh1a1 imaged 2021-12-14
Astrocyte culture lysate
55
40
WT M7
KO M5
WT M2
KO M4
WT M9
KO M8
WT M10

GAT3 blots for Figure 3G

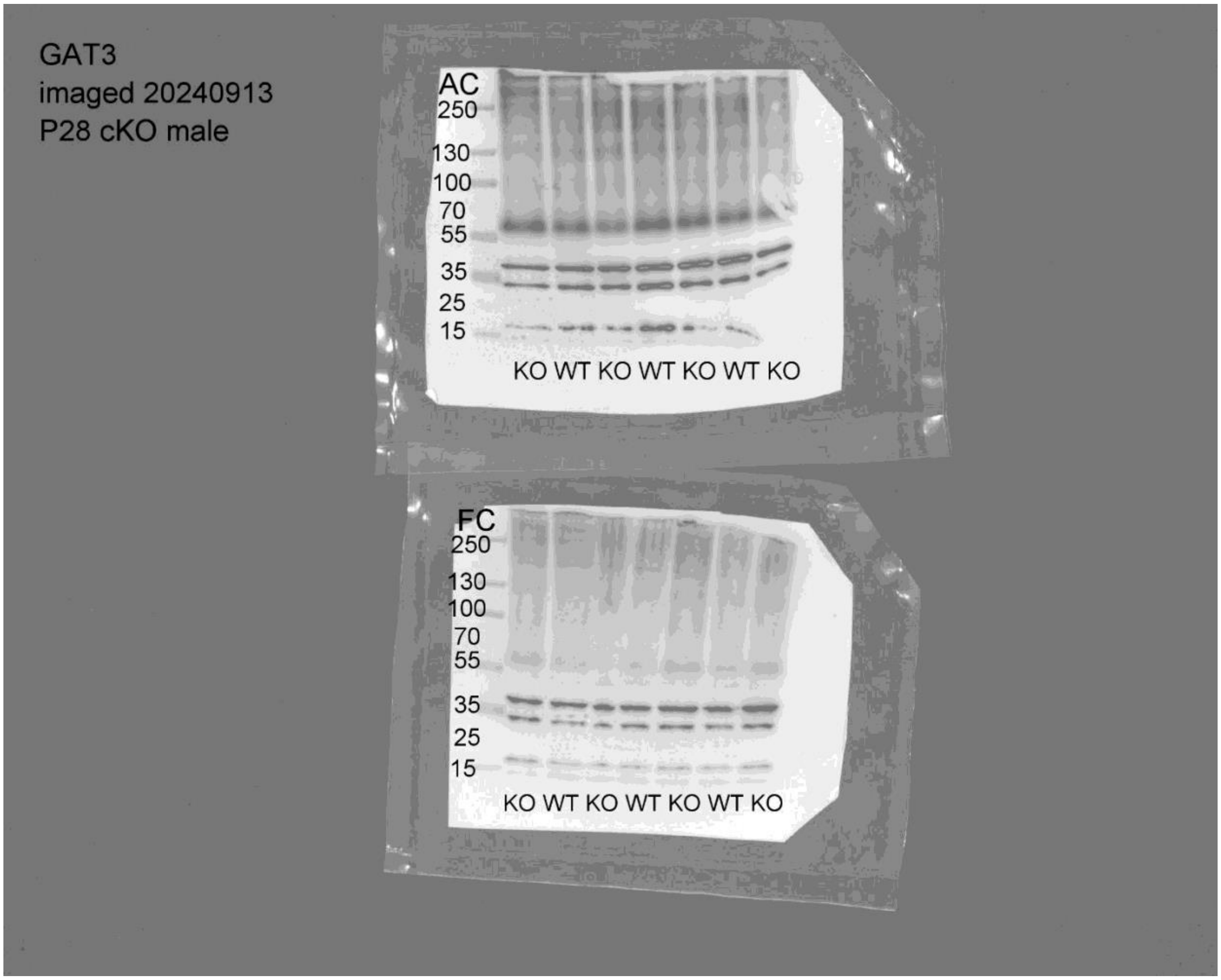

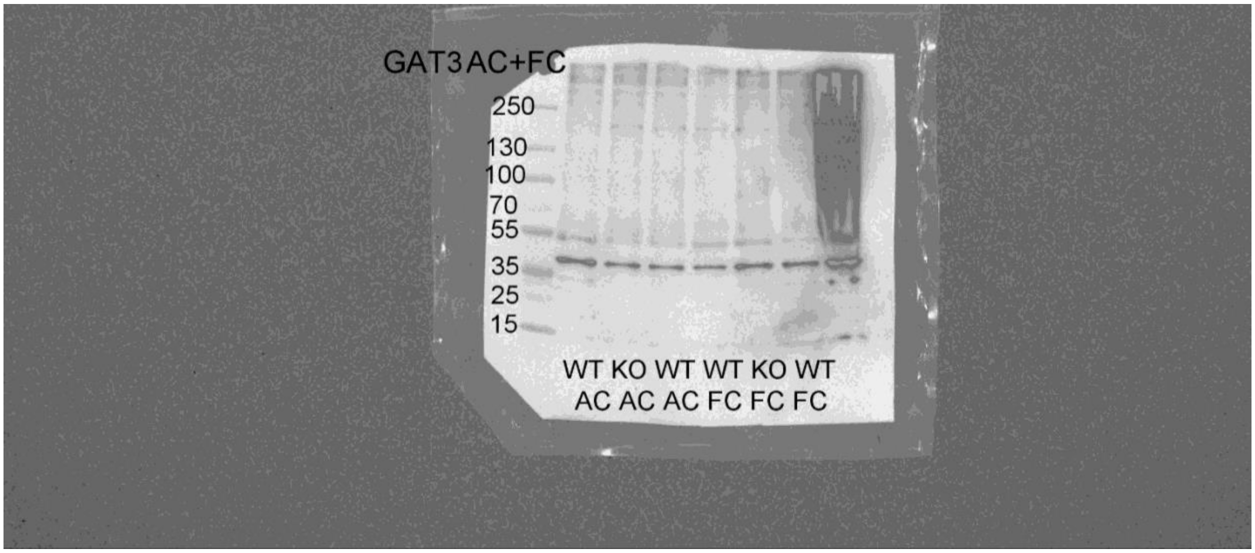

Beta-actin blot for Figure 3G, S2E

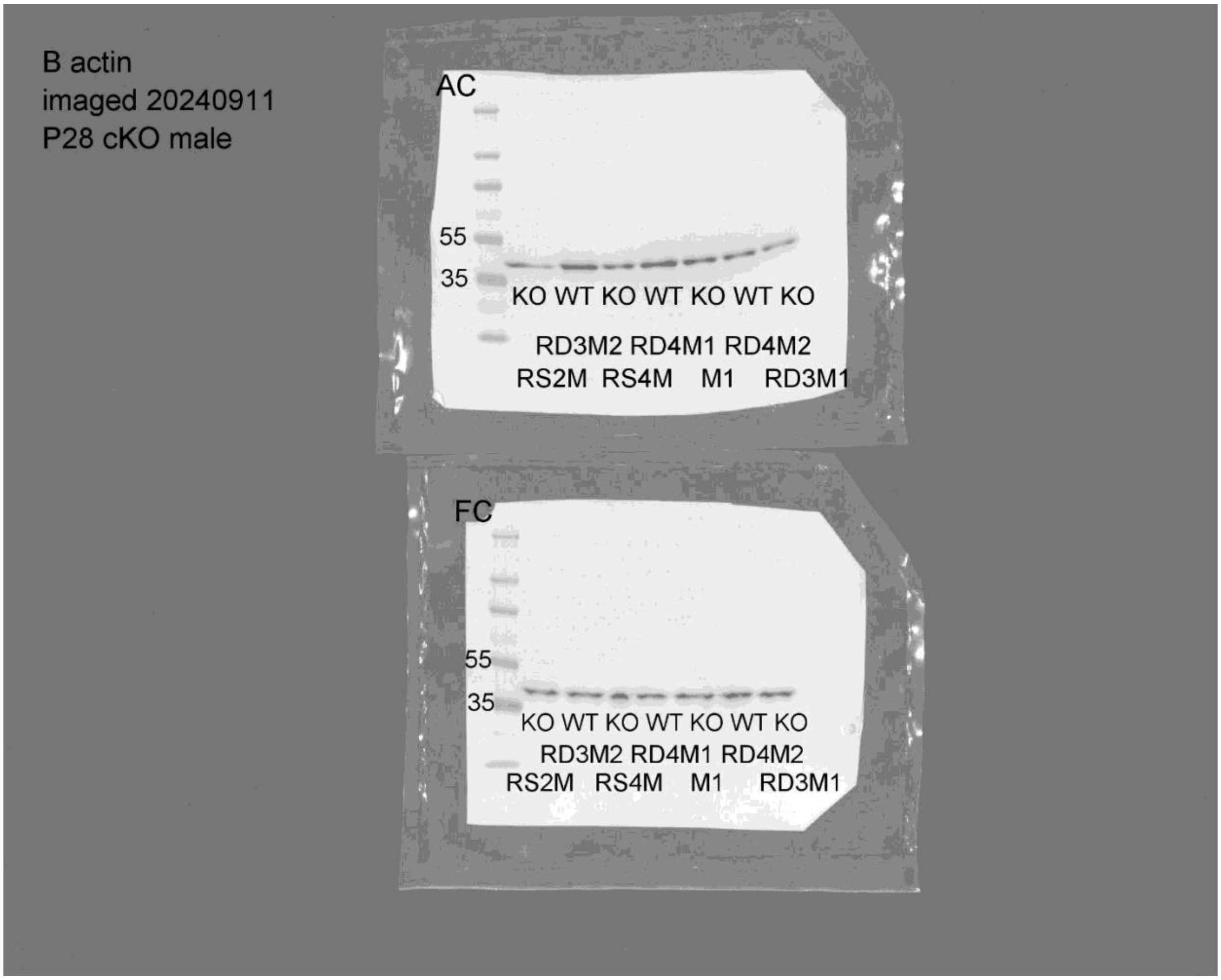

Beta-actin blot for Figure 3G

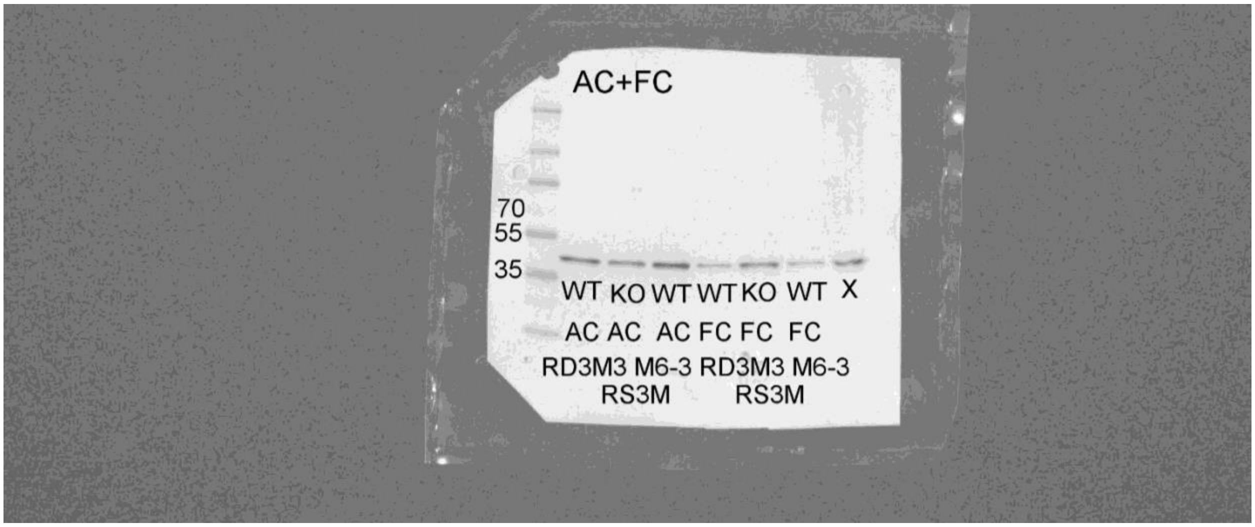

GABRg2 blots for Figure S2E

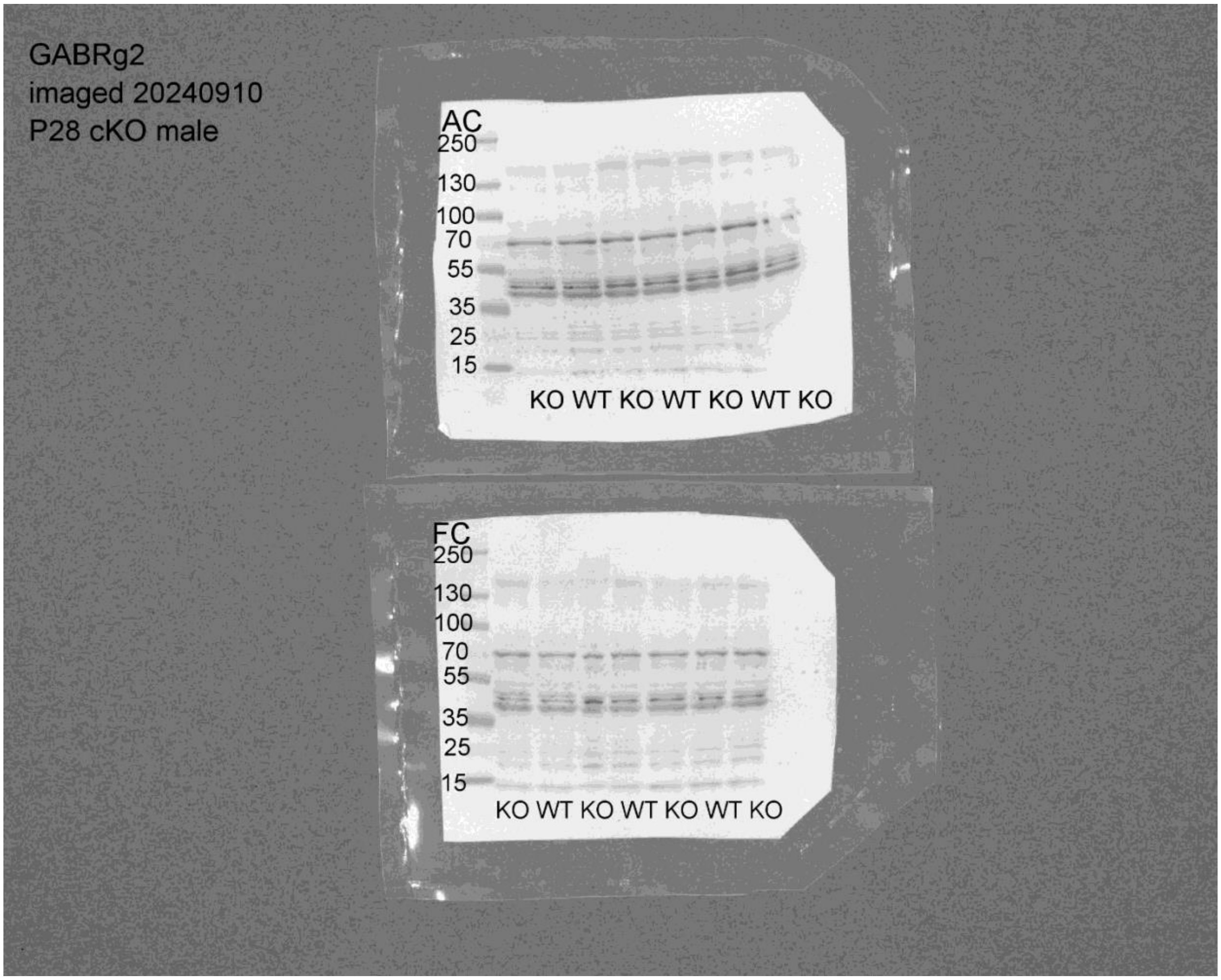

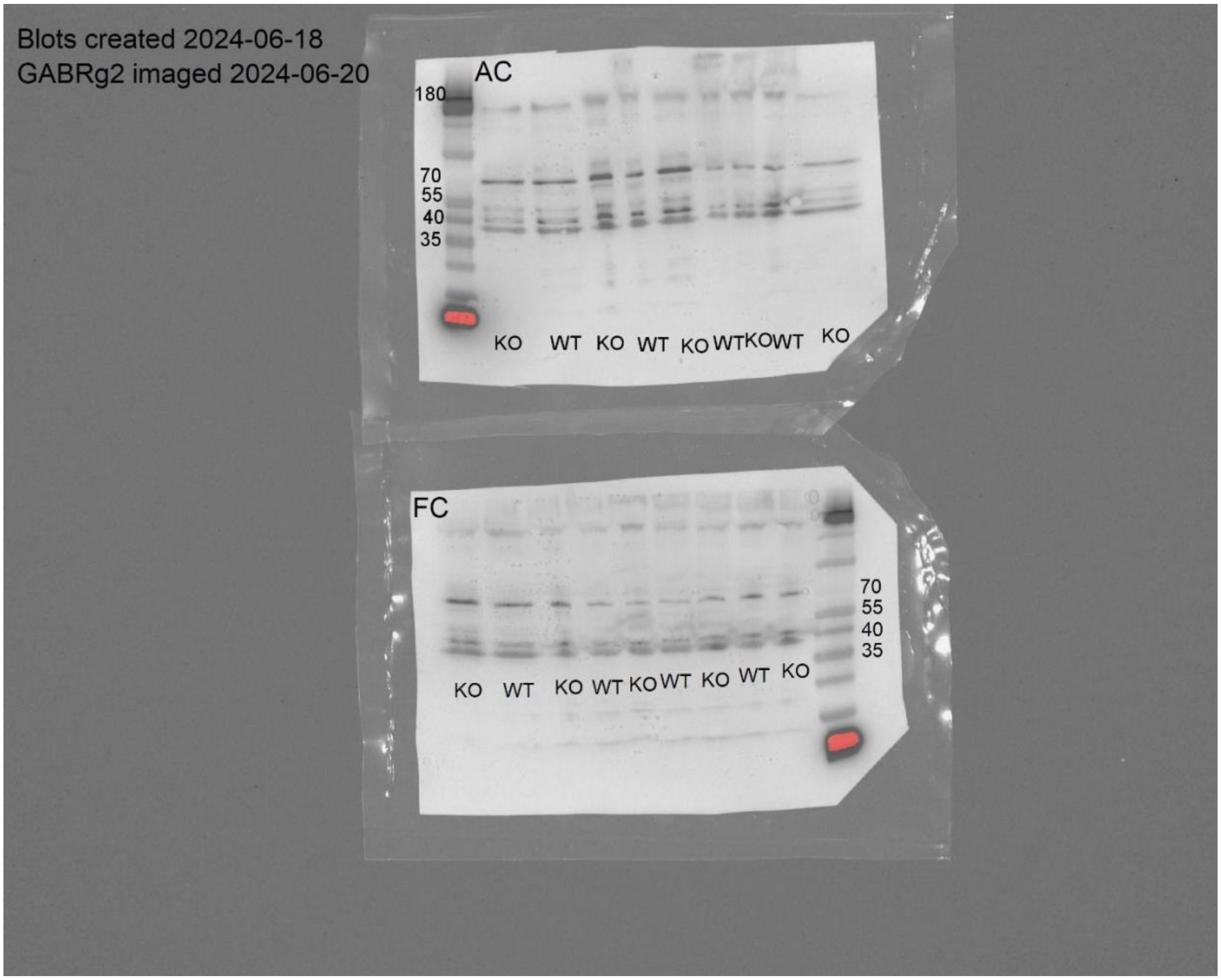
Blots created 2024-06-18
GABRg2 imaged 2024-06-20
AC
180
70
55
40
35
KO WT KO WT KO WTKOWT KO
FC
70
55
40
35
KO WT KO WT KO WT KO WT KO

Beta-actin blot for Figure S2E

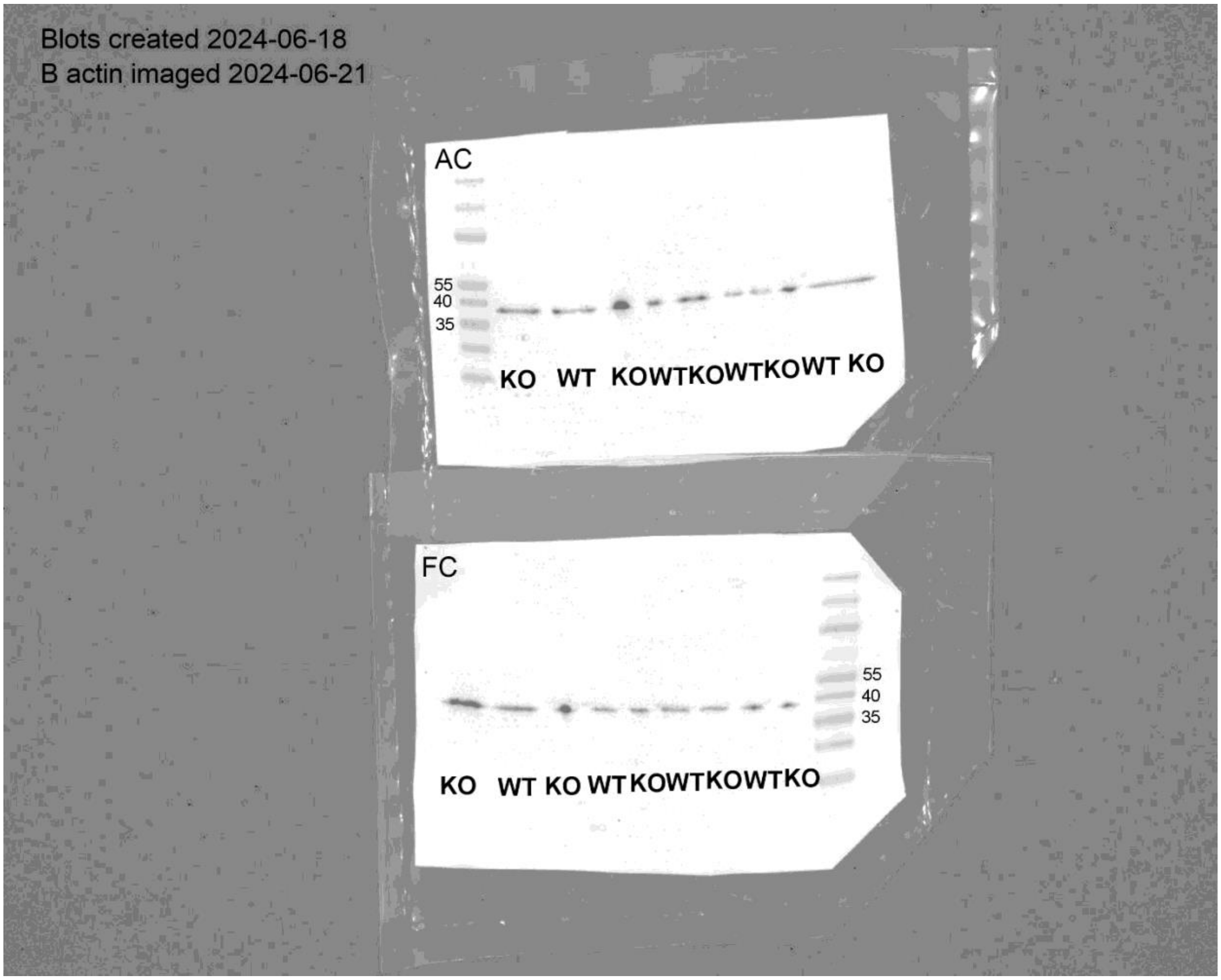

GABRa5 blots for Figure S2F(AuC first, FC second)

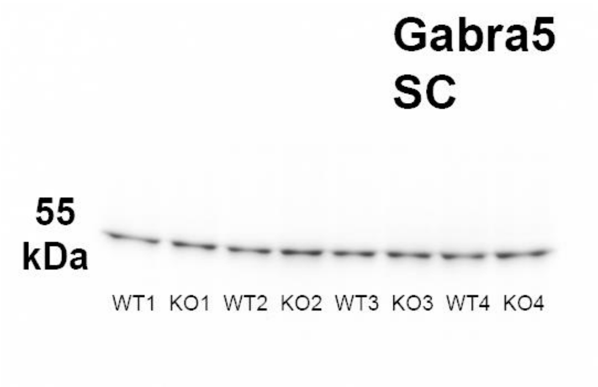

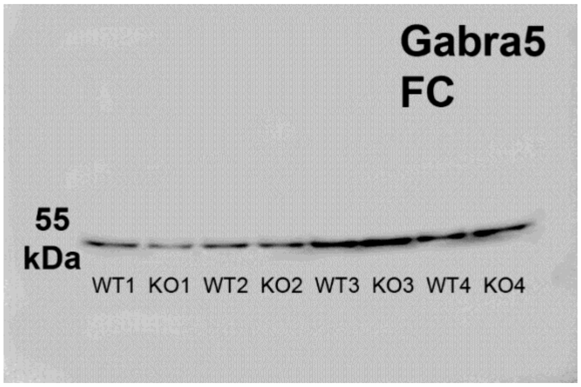

Beta-actin blots for Figure S2F (AuC first, FC second)

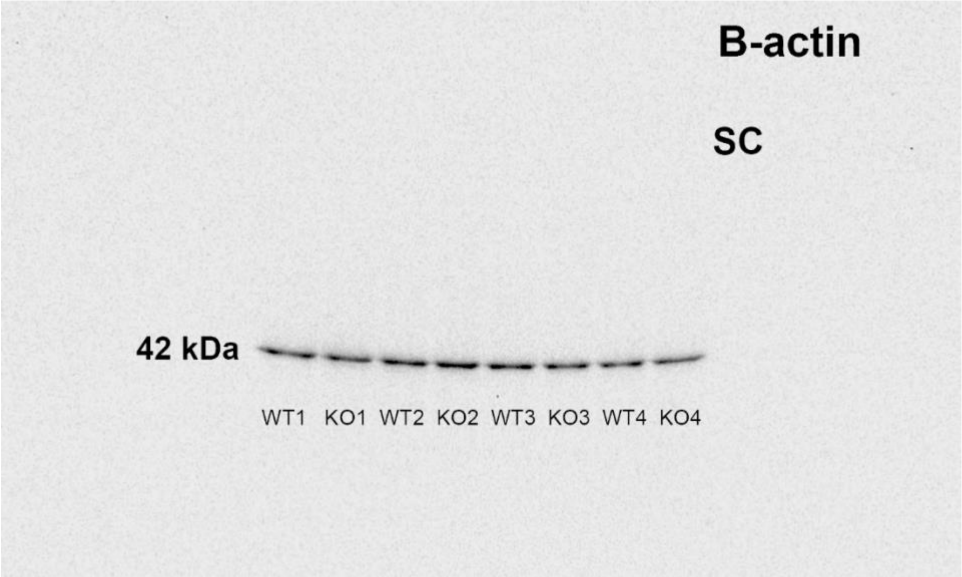

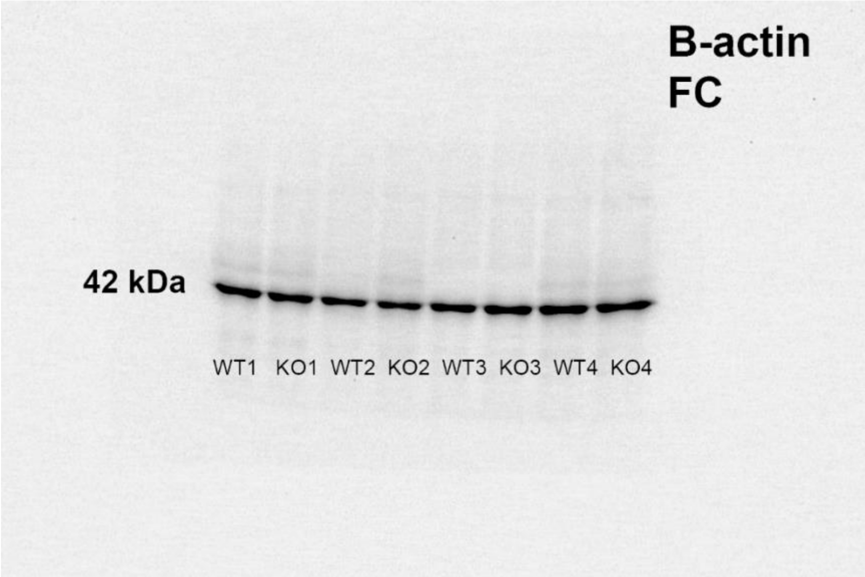
